# Unlocking Elementary Conversion Modes: ecmtool unveils all capabilities of metabolic networks

**DOI:** 10.1101/2020.06.06.137554

**Authors:** Tom J. Clement, Erik B. Baalhuis, Bas Teusink, Frank J. Bruggeman, Robert Planqué, Daan H. de Groot

## Abstract

The metabolic capabilities of cells determine their biotechnological potential, fitness in ecosystems, pathogenic threat levels, and function in multicellular organisms. Their comprehensive experimental characterisation is generally not feasible, particularly for unculturable organisms. In principle, the full range of metabolic capabilities can be computed from an organism’s annotated genome using metabolic network reconstruction. However, current computational methods cannot deal with genome-scale metabolic networks. Part of the problem is that these methods aim to enumerate all metabolic pathways, while computation of all (elementally balanced) conversions between nutrients and products would suffice. Indeed, the elementary conversion modes (ECMs, defined by Urbanczik and Wagner) capture the full metabolic capabilities of a network, but the use of ECMs has not been accessible, until now. We extend and explain the theory of ECMs, implement their enumeration in `ecmtool`, and illustrate their applicability. This work contributes to the elucidation of the full metabolic footprint of any cell.

## Introduction

Metabolism underlies most cellular behaviours. Which chemical compounds a microbe can exploit for growth, which products it can make and at which yields, is essential information for understanding the microbe’s roles in ecosystems, its responses to varying conditions, and its potentials for biotechnology and bioremediation. In the case of pathogens, metabolic capabilities are informative about the niches in which they can thrive. The functioning of multicellular organisms relies on how the capabilities of different cell types complement each other. A computational method that can enumerate all metabolic capabilities of any cell, from its annotated genome sequence, is therefore of key importance.

In the pre-genomic era, a cell’s metabolic capabilities were investigated using experimental physiological information and elemental balancing of nutrients and products. Cellular metabolism was seen as a black box: without having knowledge of the metabolic details, so-called macrochemical equations were calculated which specify the stoichiometry of the conversion of nutrients into biomass (cells) and byproducts [1, 2, 3, 4, 5, 6]. Precise measurements of heat exchange, nutrient uptake and product formation could be used to develop thermodynamic theories of cellular growth [7], which led to the improvement of biotechnological processes [5, 8]. These methods could not always be applied: they were not exhaustive, and required experimental data and basal knowledge of metabolic pathways. This information is often lacking, in particular for unculturable and extremophile microorganisms, or for cells that only survive in multi-species communities or as part of a multicellular organism. In addition, when several substrates can be consumed or multiple byproducts can be produced, a unique macrochemical equation can not be derived, and the methods need to be augmented with experimental data [3]. Nowadays, in the post-genomic era, in which the genome of any organism can be sequenced, the potential exists for comprehensive and unsupervised enumeration of all macrochemical equations of any cell. Yet, despite its great benefits, no such method is currently used, partially because most efforts focus on computation of a *highly* redundant capability set.

All metabolic reactions that a cell can catalyse can be determined from the metabolic-gene annotations of its genome. This allows for the reconstruction of the metabolic network, which can nowadays almost be done purely computationally [9] (see [10] for a recent review). The resulting genome-scale metabolic networks, or genome-scale stoichiometric models, have been determined for thousands of species. Since such a model specifies all metabolic reactions, it determines all possible pathways from substrates to products, which are conveniently described by the set of all elementary flux modes (EFMs) [11, 12, 13, 14, 15, 16]. The enumeration of all EFMs of large metabolic networks is not possible, due to a severe combinatorial explosion in their number [17], so that most research has focused on calculating only subsets of EFMs [18, 19, 20, 21, 22, 23, 24, 25, 26, 27, 28, 29, 30]. However, since many EFMs share the same overall substrates-to-products conversion and, therefore, indicate the same metabolic capability, their enumeration is not always required. Instead, for many applications it suffices to focus on all possible overall conversions that a cell can catalyse.

The complete metabolic capabilities of a cell can thus be studied by focussing on all conversions from substrates to products. An exhaustive list of these is obtained by enumeration of the Elementary Conversion Modes (ECMs), defined in 2005 by Urbanczik and Wagner [31]. ECMs are not defined in terms of the metabolic routes through the network; rather, they are defined in terms of the end results only: the feasible stoichiometries between substrates and products - the net conversion (see Box 1 for explanation). Thus, ECMs focus on the connection of an organism with its environment rather than on the metabolic pathways with which it achieves this.

ECMs can be seen as analogous objects to EFMs: the ECMs form a minimal set that generates all steady-state substrate-to-product conversions, i.e., all macrochemical equations, while the EFMs form the minimal set that generates all steady-state flux distributions. However, the set of ECMs is much smaller than the set of EFMs. First, because many different EFMs map to the same overall conversion. Second, because ECMs are objects in the lower-dimensional space of external metabolite changes, rather than in the space of reaction rates. For these reasons, the combinatorial explosion that prohibited the enumeration of all EFMs on a genome-scale network might disappear when enumerating ECMs.

Although ECMs were already defined in 2005 [31], and despite their potential for broad applicability, we could find only one study in which they were used [32]. This might be because the concept was never made accessible for a broad audience, even though it was rigorously defined mathematically. Mostly, it might be due to the absence of a readily usable computational tool that computes ECMs for general metabolic networks.

In this work, we unlock the potential of ECMs by making the theory accessible and enumeration possible for any systems biologist. We reformulate and extend the ECM theory of Urbanczik and Wagner, provide additional explanations in Boxes 1–3, and supply extensive Supporting Information where all enumeration steps are explained and mathematically supported. Most importantly, we present a Python-based enumeration program: `ecmtool`. Our software accepts metabolic models in the SBML-format as input [33], and gives an exhaustive and exact list of ECMs as output. `Ecmtool` provides both an indirect and a direct method. The indirect method is based on the algorithm proposed in [31] and is fast for small to medium-scale networks; the direct method uses a novel algorithm that lends itself to massive parallelisation and is therefore scalable to much larger networks. We validate the correctness of the computed ECMs on the medium-scale `e_coli_core`-network [34], and test the scope of `ecmtool` by enumerating the ECMs of networks of various sizes and complexity. In addition, we provide a `hide`-method that allows focusing on the conversions between a user-defined subset of the external metabolites.

This method enables the enumeration of ECMs on genome-scale models. Finally, in a collaborative, parallel study on rhizobial bacteroids, we showed that ECMs can now truly be applied to gain biological insight [35].

This work contributes to closing the gap between any cell’s genotype and phenotype. It offers a computational toolkit for the exhaustive determination of metabolic capabilities, and should be particularly valuable when experimental characterisation is impossible because cells cannot be cultured in isolation.

### Box 1: Definition of ECMs

ECMs are the minimal building blocks of all net conversions by metabolic networks, and were defined by Urbanczik and Wagner [31]. To explain their definition, we start with the stoichiometry matrix *N* of a metabolic network. Each column of *N* captures for one reaction which metabolites are consumed, which are produced, and in what ratios. To facilitate the exposition, we here assume that all reversible reactions are split into a forward and backward reaction, so that all reactions in *N* are irreversible. Some metabolites are internal to the cell and some metabolites are external; metabolites that occur both inside and outside the cell are considered as two metabolites: one internal and one external. We denote the index set of internal metabolites by *Int*. The product of the stoichiometric matrix with the vector of reaction rates ***v***, gives the rates of change of all metabolite concentrations, i.e., the conversion: ***ċ*** = *N**v***. Metabolism is assumed to be in steady state, so that all internal metabolite concentrations are constant: ***ċ***_*i*_ =0 for all *i* ∈ *Int*. The space of all steady-state conversions, and thus of all metabolic capabilities, is given by

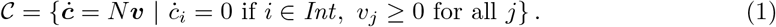

This space is called the *conversion cone*, and should not be confused with the *flux cone* which comprises all steady-state fluxes. In fact, the conversion cone is the result of multiplying all points in the flux cone with the stoichiometric matrix, see SI Section 2 for more explanation.

#### Definition 1.

*The set of* Elementary Conversion Modes (ECMs) *is the minimal set of conversions* {*ecm*_1_,…, *ecm_K_*} *such that each steady-state conversion can be written as a positive sum of ECMs, without the production of any external metabolite being cancelled in that sum*.

Some readers might note that this definition of ECMs is similar to the definition of EFMs. This is because both can be defined as elementary vectors [36, 37]: ECMs are the elementary vectors of the conversion cone, while EFMs are the elementary vectors of the flux cone. The values in an ECM indicate the changes of metabolite concentrations, while the values in an EFM indicate reaction rates.

We will explain the two parts of Definition 1 using the toy network example of Figure 1, in which the external metabolites *A, B, BM* are interconverted via internal metabolites *C, D*, and *E*.

**Figure 1:**
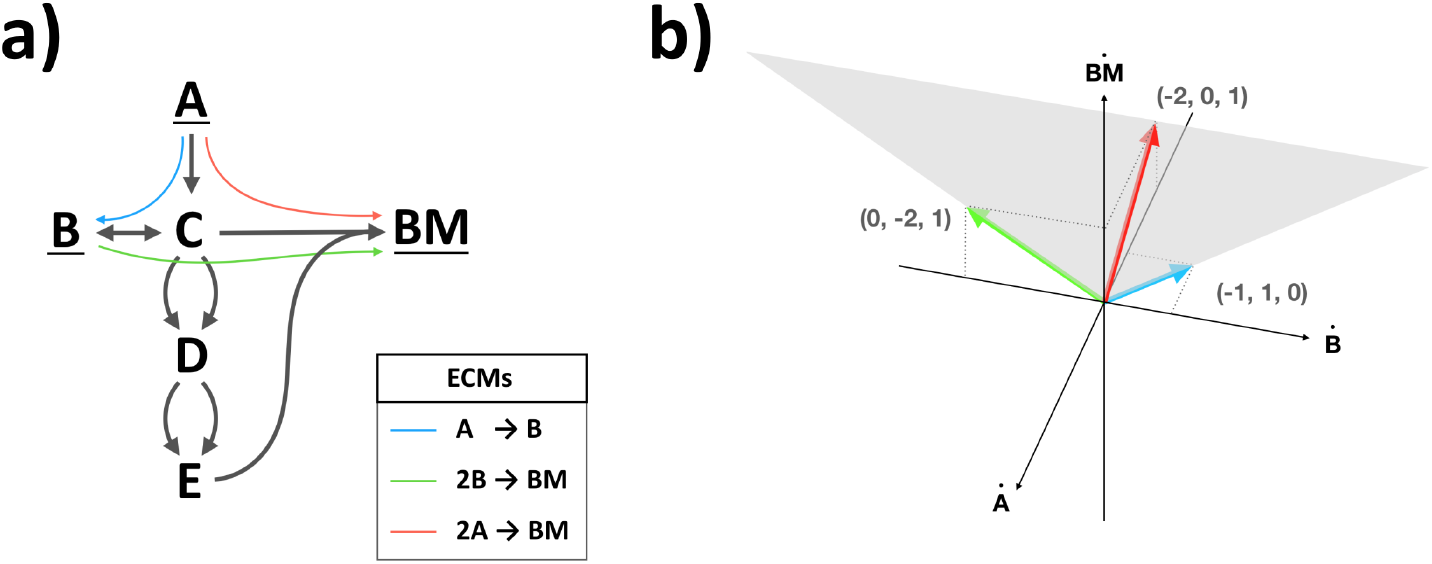
**a)** The Elementary Conversion Modes for a small network are shown in blue, green and red. Notice that the red ECM can be written as a positive combination of the blue and green ECM, but that this cancels the production of *B*. **b)** The cone of steady-state conversions is shown in gray, and is spanned by the blue and green ECM. The red ECM lies in the interior of the cone on the intersection with the 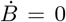 -plane.

All steady-state conversions together form the conversion cone, which is a “convex polyhedral cone’’ (shaded area in 1**b**)). As a consequence, the steady-state conversions can be fully described by the extreme rays of this cone (blue and green in the figure). Indeed, any steadystate conversion can be written as a positive sum of the extreme rays. By the first part of Definition 1, this means that these extreme rays are Elementary Conversion Modes. The example therefore has at least two ECMs: A → B (blue) and 2B → BM (green).

It is important to note that any *positive* sum of steady-state conversions is again a steadystate conversion. This makes sense in biological terms: a conversion lies in the cone if there exists a set of reactions that gives rise to the conversion, and satisfies the irreversibility and steady-state constraints from Equation (1). So, if we have several sets of reactions that correspond to conversions, their sum will correspond to the summed conversion. However, a sum in which some extreme conversions are added *negatively* does not necessarily result in a feasible conversion, because the resulting conversion might not be feasible without using an irreversible reaction in the negative direction.

Now consider the conversion 2A → BM (red). This conversion can be written as a positive sum of the two previously found ECMs: 2(A → B) + (2B → BM)=(2A → BM). However, in summing these ECMs, the metabolite *B* is cancelled, since it is produced *and* consumed. Since, the ECMs are intended to capture a complete set of minimal building blocks of biologically realistic conversions, taking only the extreme rays does not suffice: we also want to describe the possibility of producing *BM* from *A* without simultaneously excreting and consuming *B*. The second part of the ECM-definition therefore ensures that these conversions are added to the set of ECMs as well: the ECMs should generate all conversions *without cancellation of the production of any metabolite*. If we would take a combination of the blue and green extreme conversions, this would always (partly) cancel the production of *B*, since *B* is produced in the blue conversion and consumed in the green conversion. Therefore, the red conversion is also an ECM: since this conversion does not produce or consume *B*, a positive combination with the other ECMs does not induce a cancellation. In total, we thus have three conversions, as listed in Figure 1**a)**.

In mathematical terms, one could obtain the full set of ECMs by calculating the extreme conversions per orthant, and then taking the union of all these extreme conversions. The requirement that no metabolite production is cancelled, implies that all steady-state conversions can be written as a positive sum of ECMs in which each metabolite is either produced by all ECMs in the sum, or consumed by all ECMs in the sum. In this manner, cancellations no longer occur (see [37]).

## Results

### Cells have orders of magnitude fewer metabolic capabilities (ECMs) than flux routes (EFMs)

The number of ECMs increases much slower with metabolic-network size than the number of Elementary Flux Modes (Figure 2). For example, the number of elementary modes in the `e_coli_core` model [34] reduces from 100,274 EFMs to 689 ECMs. The number difference is likely even greater for larger genome-scale metabolic networks. This makes ECM visualization possible, which facilitates their exploration and analysis (Figure 3). This illustrates that it is more direct and efficient to enumerate ECMs, which are the metabolic capabilities of a cell, instead of EFMs, which are flux routes that often have an identical metabolic capability, i.e. net conversion of cellular nutrients into products (Figure 2A).

**Figure 2:**
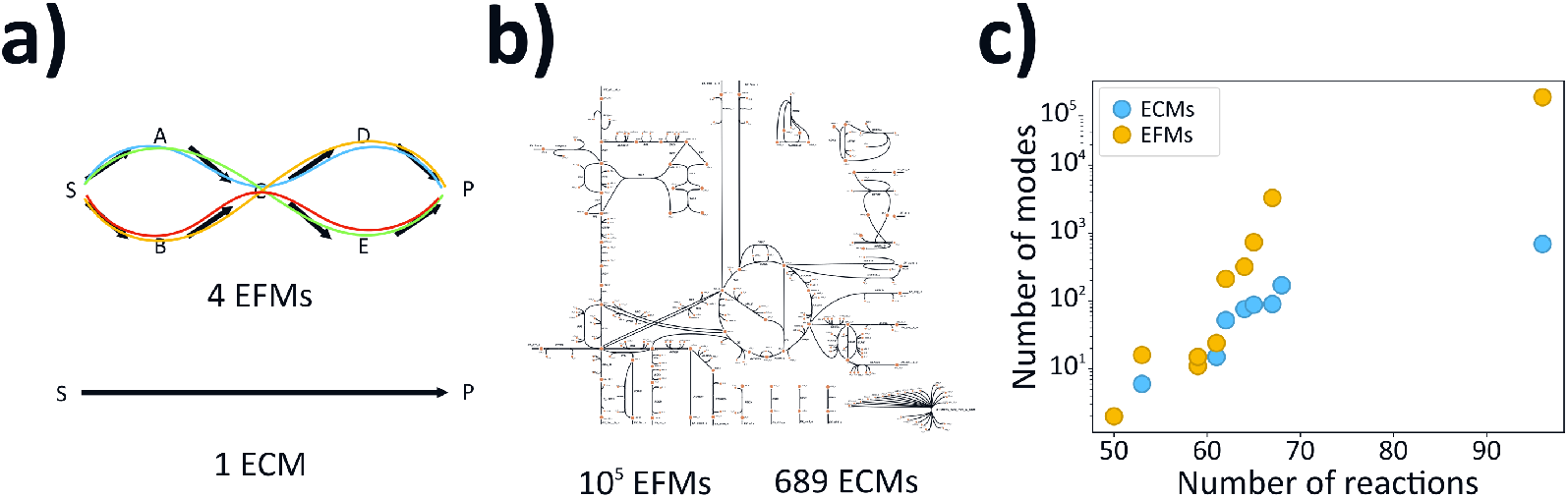
The number of ECMs remains orders of magnitude lower than the number of EFMs. **a)** Because many different EFMs refer to the same overall metabolic capability, the number of ECMs is much lower than the number of EFMs. **b)** EFM-vs-ECM numbers in the `e_coli_core`-network. **c)** Subnetworks of the `e_coli_core`-network were selected (see SI 10.1) to illustrate how the number of ECMs and EFMs scales with network size.

**Figure 3:**
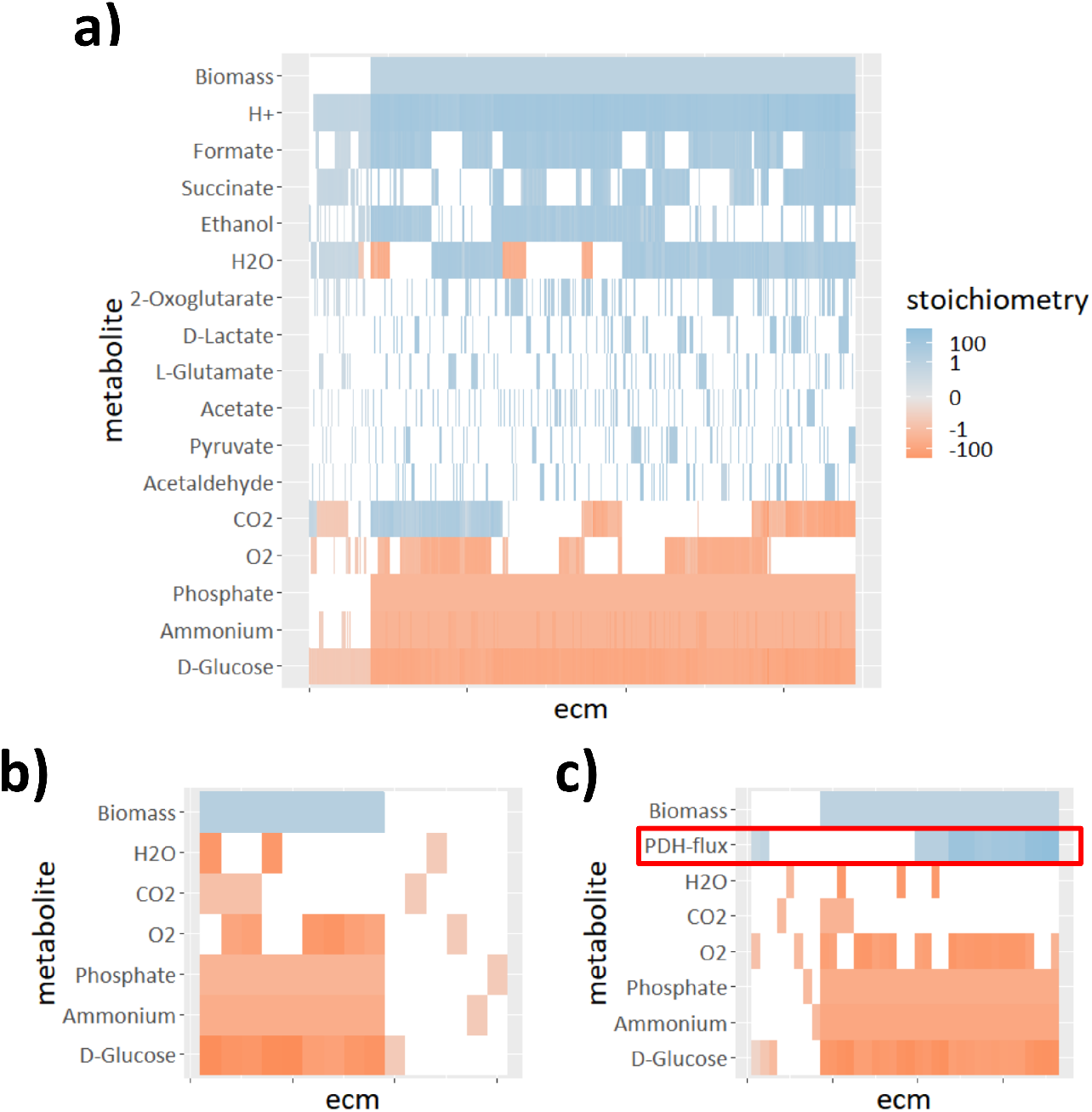
The full metabolic potential of the e_coli_core-model. **a)** The ECMs of the full model are shown as the different columns; each row corresponds to a different external metabolite. The colourscale indicates the stoichiometric coefficient of the metabolite in the conversion: blue for production, and red for consumption. The coefficients were log-transformed to allow for visualization of differences in both large and small coefficients (details and `R`-code can be found in SI 10.2); small values are shown in gray, while zero values are white. Of the 689 elementary conversions, 613 lead to the production of biomass. These ECMs were normalised to fix the biomass production at 1, while the other ECMs were normalised such that the sum of absolute coefficients is 1. **b)** If we use the `hide`-method, explained in Box 2, to hide the production of metabolites, we get 15 ECMs that span all possible ratios in which substrates can be converted into biomass. This smaller set of ECMs is easier to compute and easier to explore, while the steady-state assumption is still satisfied in the whole network. So, even though the secretion of products is not reported, it has been implicitly taken into account, so that all relations between substrates shown in **a)** are captured in **b)**. **c)** If we use the `tag`-method, also explained in Box 2, to report the activity of the pyruvate-dehydrogenase (PDH) reaction, we find 36 ECMs that summarize all possibilities. It can be seen that the PDH-reaction is not essential for growth, but that it seems to be necessary for efficient growth on glucose since the uptake of glucose is generally lower when PDH is active.

A major advantage of ECMs is that they can be computed for metabolic networks for which EFM- enumeration is not possible. For example, we found 874,236 possible ECMs for the pathogen *Helicobacter pylori* in a minimal medium (`iIT_341` [38] 485 metabolites and 554 reactions), while EFM enumeration ran into memory errors, most likely due to the enormous number of EFMs in this model (the full set of ECMs is available as a supplementary file). We note that a set of hundreds of thousands of ECMs might appear hard to analyze, but the user can easily filter out a relevant subset once such a set is obtained (see Supplemental Figure 1 for an example).

Summarizing, the enumeration of ECMs by `ecmtool` allows for the determination of all the metabolic capabilities of metabolic networks for which EFM enumeration is no longer feasible. We did find that the number of ECMs in the genome-scale *E. coli* network `iJR904` [39] (761 metabolites, 1075 reactions) is still too large to be computed by `ecmtool`. However, even for models of this size `ecmtool` still provides useful information. In Box 2 and Figures 5 and 6, we show how focussing on essential information allows networks of this size to be analysed.

**Figure 4:**
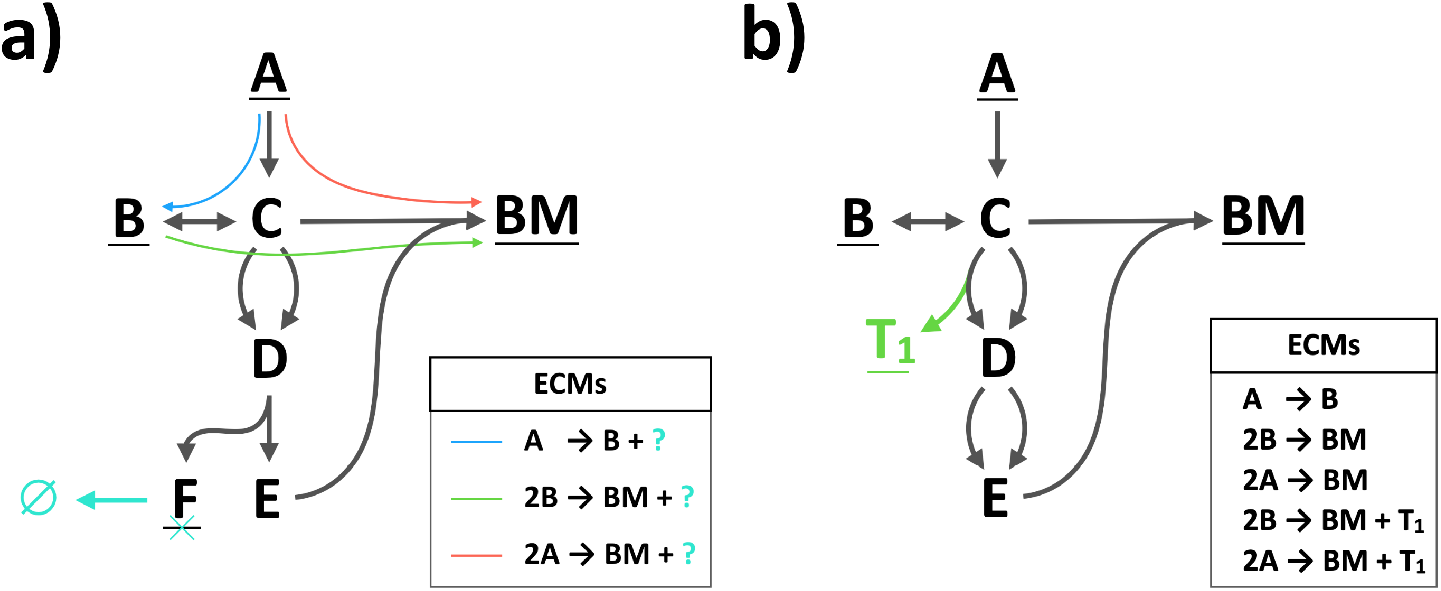
**a)** Information about the production of F can be ignored if F is marked as internal and a virtual reaction (cyan) is added that converts metabolite F into nothing. This strategy aids in fast computation, because the resulting set of ECMs is generally smaller (see also the worked-out example in the Methods-section). **b)** Information about the usage of a reaction can be uncovered by coupling the production of a virtual metabolite (T_1_, shown in green). The coefficient of T_1_ in the resulting ECMs denotes the rate of the reaction of interest.

**Figure 5:**
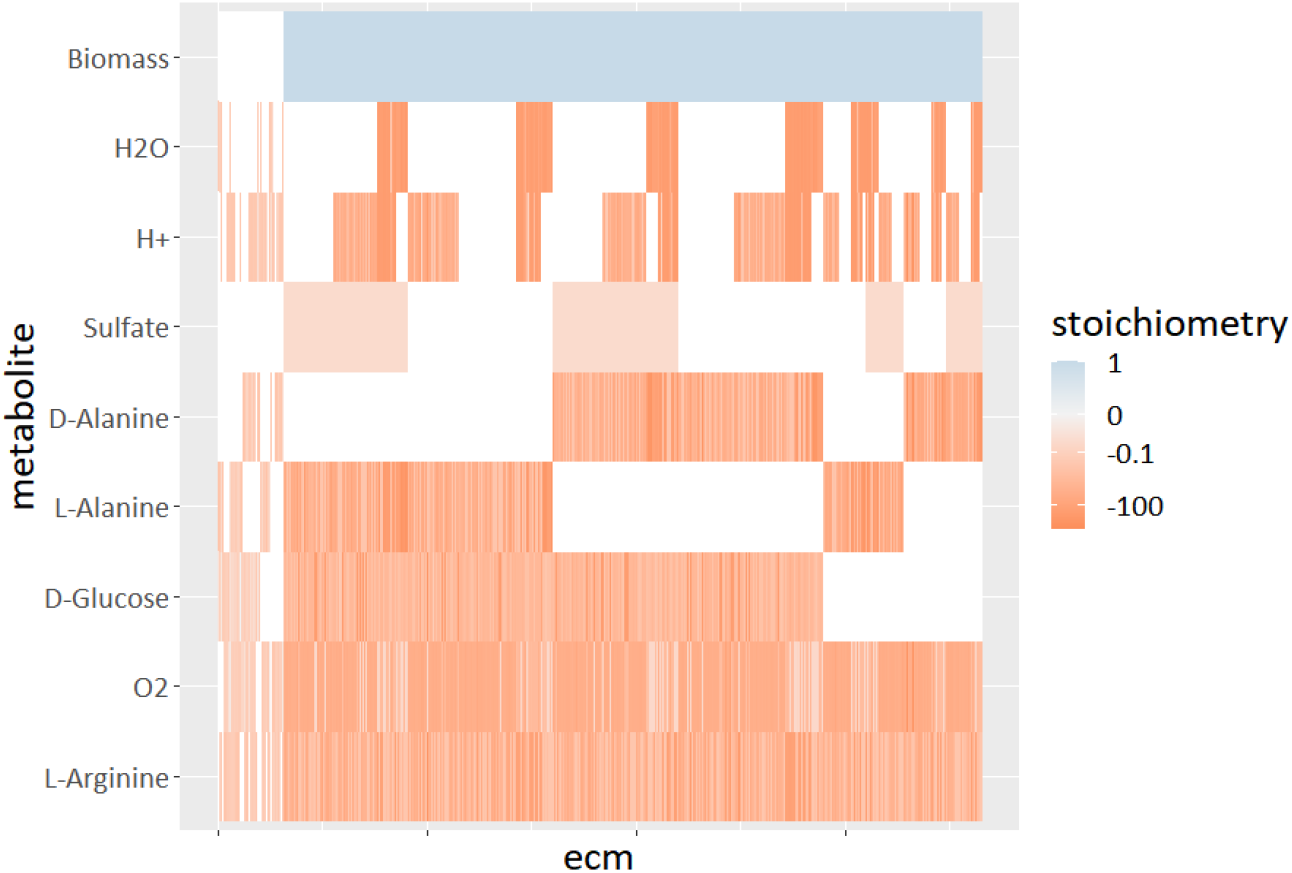
Focusing on substrate uptake shows the minimal needs of *Helicobacter pylori*. We computed the ECMs, shown as different columns, for the `iIT341`-model by allowing for the uptake of all metabolites of a supposedly minimal medium proposed by the developers of the model (MinII from [38]). All output metabolites were hidden, using the `hide`-method outlined in Box 2. The uptake of nine substrates is not shown here because these were equal for all ECMs, indicating that these are directly coupled to biomass formation. The colourscale indicates the log-transformed coefficients of the metabolites in the conversion, where metabolite production is shown in blue and consumption in red (details and `R`-code can be found in the SI, Section 10.2). The ECMs are normalised such that biomass production, if nonzero, is 1, otherwise the sum of the absolute coefficients is fixed to 1. The ECMs were clustered using hierarchical clustering. The block-like ordering of the ECMs indicates that substrate usage of *H. pylori* is largely modular: the uptake of one substrate seems independent of the uptake of another.

**Figure 6:**
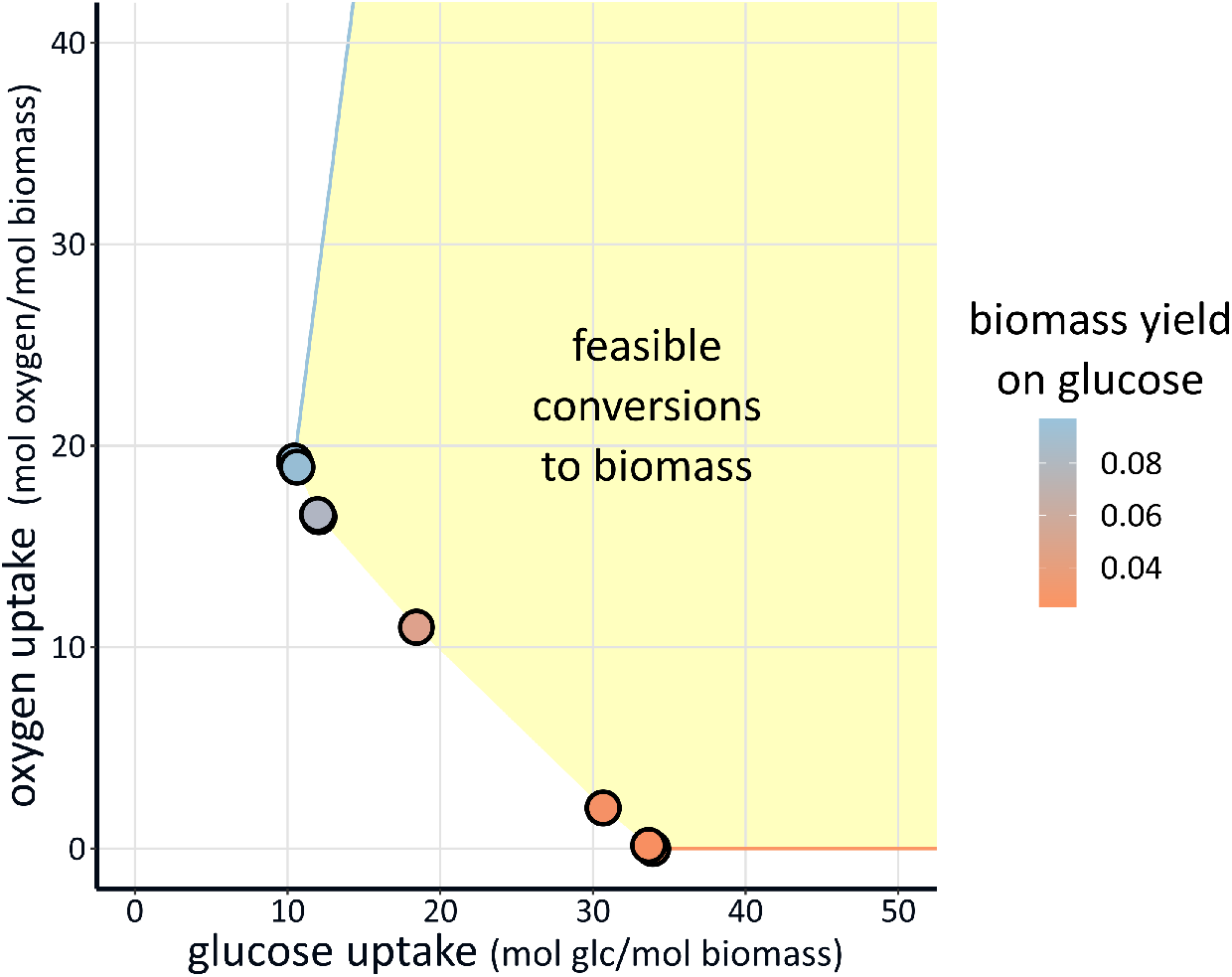
Few conversions from glucose and oxygen to biomass cover *E. coli*’s full flexibility. We calculated the ECMs for the genome-scale *E. coli*-model `iJR904` [39] by hiding all external metabolites except for glucose, oxygen and biomass. This gives 12 ECMs that span all possible biomass-yields on glucose and oxygen. The dots show, for the 10 ECMs that produce biomass, the necessary glucose and oxygen uptake to produce one unit biomass. The other 2 ECMs give the most extreme conversions from glucose and oxygen to non-biomass products, consuming only glucose (red arrow), or consuming the most oxygen per glucose (blue arrow). The convex combinations of biomass-producing ECMs combined with positive multiples of the non-biomass-producing ECMs, give all feasible ways to produce one unit biomass (yellow area).

### Validation of ecmtool for ECM enumeration

We validated the results of `ecmtool` in several ways. First, we have computed the ECMs on many small models for which we could still check the correctness and completeness of the results by hand. Second, we used the `e_coli_core`-model, for which we could still use the set of EFMs enumerated by `efmtool`, to validate our results. The `Matlab`-code that we used for this validation is provided as a supplementary file.

The correct set of ECMs should satisfy three properties: 1) each ECM must be a steady-state conversion, 2) each ECM must be an elementary vector, and 3) each steady-state conversion must be a positive combination of ECMs without metabolites being cancelled in the sum.

We confirmed that all computed ECMs are steady-state conversions by checking that the net production of internal metabolites equals zero, and that there exists a combination of metabolic reactions that gives rise to the ECM. Then, according to the definition of ECMs given in Box 1, we proved that each ECM is elementary by showing that it cannot be written as a positive sum of the other ECMs without the production of any external metabolite being cancelled.

The third property was harder to validate, because how can we prove for *all* steady-state conversions that they can be written as a combination of ECMs? We chose to use the set of Elementary Flux Modes calculated by `efmtool`. This set spans all possible steady-state flux combinations the metabolic network allows. For each EFM, we then calculated its overall conversion, and tried to write this conversion as a combination of ECMs. If we allowed for an error of 10^-7^, then each conversion could be decomposed into ECMs. This error margin was necessary because the results from `efmtool` are affected by round-off errors. The computed ECMs do not suffer from round-off errors because the computation by `ecmtool` uses fractions only. Although this slows down many of the calculations, this is necessary to maintain the accuracy of the computed ECMs. For example, for the Double Description method it is known that round-off errors can grow to a non-negligible size [40].

Above, we explained and validated that `ecmtool` finds all Elementary Conversion Modes, given an *annotated* genome. The annotation is necessary for the reconstruction of the metabolic network. Strictly speaking, this minimally requires the annotation of the *metabolic* genes. Since the annotation of a genome is not always complete, we cannot guarantee that all metabolic capabilities encoded on the genome are found. We can guarantee that all conversions are found of the genome-derived metabolic network.

#### Box 2: Hiding and tagging enables focusing on the most important metabolic conversions

In `ecmtool`, the user can choose to compute only the stoichiometric relations between a subset of the external metabolites by ‘hiding’ the other external metabolites. The resulting set of ECMs still gives a full summary of these relations, and complies with the steady-state assumption on the full metabolic network. The consumption and production of the hidden metabolites still occurs, but is not reported. As a result, the reported ECMs are not necessarily mass-balanced, which is emphasised by the question marks in Figure 4**a)**. An ECM computed with the `hide`-method thus gives a ratio in which the non-hidden metabolites can appear in a conversion, but it does not give any information about which hidden metabolites are consumed or produced in such a conversion. In return, the `hide`-method facilitates ECM-enumeration on much larger networks, because fewer ECMs are needed to describe all conversion relations between the smaller set of non-hidden metabolites. Therefore, ECM-enumeration with hidden metabolites can take an organism’s full metabolic complexity and summarize its metabolic capabilities regarding a few variables that are of interest.

In Figure 4**a)** we show how metabolite F can be hidden in the ECM-computation by adding a reaction that converts it into nothing (sometimes called a demand reaction). In general, a metabolite is hidden by adding a reaction that creates it from ‘nothing’, turns it into ‘nothing’, or both, depending on whether the metabolite can only be consumed, only produced, or both, respectively. Then, the metabolite is marked as an internal metabolite, so that the steady-state assumption is imposed. The added reaction can always make sure that the net consumption or production of the metabolite is zero. As a result, the hidden metabolite will vanish from the computations in an early stage of the enumeration, thereby reducing computation time. We illustrate this in the worked-out enumeration example in Box 3.

In the example of Figure 4**a)**, we obtain the conversions between non-hidden metabolites A, B and BM, ignoring the information about whether F is produced during these conversions or not. If metabolites are hidden, the computed conversions should be interpreted with care, acknowledging that the reported conversions are possibly not elementally balanced (since the hidden metabolites are excluded from this report). In the example, we emphasize that we do not know whether *F* was produced in the conversion by adding question marks on the right side of the conversion notation. If metabolites that can be consumed by the cell are also hidden, then question marks should be placed on the left side as well.

Besides hiding metabolites of minor importance, we can keep track of reaction rates of major importance. The `tag`-method, suggested in [31], adds a virtual external metabolite that is produced whenever the reaction of interest is used. As a result, one unit of virtual metabolite is produced when the tagged reaction runs at a rate of one. Since an ECM reports the stoichiometric coefficients of all metabolites in the conversion, the coefficient of the virtual metabolite in the ECM reflects the rate at which the tagged reaction must run to produce the conversion. This method will show to which conversions the reaction of interest contributes, possibly providing valuable information about the essentiality of that reaction. In Figure 4**b)**, we show an example of such reaction tagging. One of two reactions from C to D is extended to produce virtual metabolite T_1_, resulting in the reaction C → D + T_1_. Any conversion that uses the reaction of interest produces T_1_, and its coefficient in the conversion is equal to the reaction rate.

In Figure 3 we illustrate the `hide`- and `tag`-methods in the `e_coli_core` model to respectively highlight the different possible combinations of growth substrates, and the necessity of the pyruvate-dehydrogenase reaction in these conversions.

### Focusing on subsets of metabolites enables genome-scale calculation of metabolic capabilities

Focusing on the stoichiometric relations between metabolites of major importance by hiding external metabolites of minor importance is a powerful way to scale up the size of metabolic networks that can be dealt with in `ecmtool`. The ECMs that are obtained now span all possible relations between the non-hidden metabolites, but no longer give information about what happens to the hidden metabolites (see Box 2 for a more elaborate explanation). Importantly, the steady-state assumption remains satisfied and all hidden metabolites can be produced or consumed, even though this production or consumption is not reported.

To illustrate how the `hide`-method can help focusing on the most important metabolic capabilities of a network, we focused on the minimal growth strategies that the pathogen *Helicobacter pylori* can employ.

We took the `iIT341`-model for which we already calculated the full set of ECMs (see Supplemental Figure 1), and hid all information about product secretion (Figure 5). The 3652 ECMs that were obtained thus span all possible proportions in which the different nutrients can be consumed. The results show that the only mutual dependency between the uptake of different nutrients is between D-alanine and L-alanine, one of which should always be consumed. This independence indicates a modular design of the nutrient uptake system of *H. pylori*, which might benefit its flexibility when living in the human stomach.

If we focus only on the conversion of glucose and oxygen into biomass, we could even compute the ECMs for a genome-scale model of *E. coli*: iJR904 [39], containing 761 metabolites, 1075 reactions (Figure 6). According to their definition, the resulting ECMs form a minimal spanning set of all feasible conversions from glucose and oxygen to biomass. This implies that the set of ECMs contains the most ‘extreme’ conversions. Therefore, we can use them to draw the full Pareto front between the biomass yield on glucose and on oxygen, extending a method used by Carlson and Srienc to genome-scale models [41]. It turns out that this Pareto front is completely determined by 12 ECMs. For each of these ECMs, we can find a combination of reaction rates that gives rise to this conversion. In doing so, we obtain twelve states of metabolism that fully determine *E. coli*’s flexibility to optimise its growth rate in glucose- and oxygen-limited conditions. A Flux Balance Analysis where glucose and oxygen uptake is constrained and biomass production is maximized, will always result in a combination of these metabolic states.

### Case study: a metabolic capability study of an unculturable *rhizobia* strain with ecmtool

Rhizobia are soil bacteria that can induce formation of nodule structures on plant roots, in which they differentiate into non-dividing bacteroids. Bacteroids fix atmospheric nitrogen into ammonia and make this available to the plant in exchange for carbon in the form of dicarboxylates [42]. Although a metabolic network was reconstructed, physiological information about rhizobial bacteroids is lacking, because they are difficult to isolate and extremely fragile [43]. In addition, analyzing the metabolic network with an optimization approach like Flux Balance Analysis [44] is unfavourable because it is unclear what the optimization objective would be. After personal correspondence, `ecmtool` was used by Schulte et al. to enumerate the metabolic capabilities of *Rhizobium leguminosarum* [35]. This aided in exposing the role of oxygen supply in the observed amino acid secretion and carbon polymer synthesis by bacteroids, and in quantitatively reproducing the carbon cost of biological nitrogen fixation.

## Discussion

### Relevance of ECMs

Our method enumerates and quantifies, for any organism for which a metabolic reconstruction has been made, all possible stoichiometric relations between substrates, products and biomass. This method does not rely on any optimality assumption, nor does it require experimentally obtained physiological information. It uncovers the full metabolic capability of an organism, and with that the metabolic footprint that an organism may leave in its environment.

ECM enumeration stands in a long tradition of methods that pursue this goal [45]. Some of these methods attempt to find an exhaustive list of reaction pathways that a cell is capable of, for example calculating Extreme Currents [46], Elementary Flux Modes [11], or Elementary Pathways [47]. These methods all have in common that scaling to genome-scale metabolic networks is impossible because of the rapid growth of the number of pathways with network size [17]. Other methods try to view the cell as a black box and focus on what is consumed and what is produced, leading to the concepts of macrochemical equations [3, 5], direct overall reactions [2], and eventually to Elementary Conversion Modes [31]. ECM enumeration is the only method that provides a complete set of metabolic capabilities, takes reaction irreversibility into account, and scales to genome-scale networks.

### Applications of ECM enumeration

The enumeration of ECMs facilitates the exploratory study of metabolic networks: investigation of the ECMs could spark new hypotheses and show unexpected connections. It therefore complements optimization approaches like Flux Balance Analysis [44] that are efficient at answering questions that are known beforehand. Even in the case that optimization approaches are more efficient, elementary mode analysis provides additional insight. For example, EFM-analysis was used to understand an adaptive growth strategy of *Lactobacillus plantarum* that was observed experimentally and predicted by Flux Balance Analysis (FBA) [48]. In this specific case, the analysis could be restricted to primary metabolism which facilitated the EFM-computation, but this restriction is often biologically unreasonable. In the future, ECMs could replace EFMs, such that this approach can be more generally applied. Carlson and Srienc [41] used the set of Elementary Flux Modes in a relatively small *E. coli* model to investigate optimized *E. coli* growth in carbon- and oxygen-limited conditions. Using this approach, they could simplify their analysis by selecting four Elementary Flux Modes that together determined all optimal growth strategies in different glucose- and oxygen-limited conditions. In Figure 6 we showed that with `ecmtool` this approach can be generalized to genome-scale models.

Most analyses of metabolic networks require *a priori* physiological information that is often not available. For example, it is often required to impose constraints on exchange fluxes, to choose a reaction rate that needs to be optimized, or at least to know which metabolites can be produced [3]. This hinders the investigation of species that are insufficiently characterized and difficult to culture. Moreover, for many organisms it is doubtful whether reaction rates are optimized at all, for example for pathogens or the composing cells of higher eukaryotes. ECMs do not require extensive information, solely a reconstructed metabolic network. The decisive role that ECM enumeration can play in the study of unculturable and non-optimized organisms is exemplified by the recent application of `ecmtool` to investigate the symbiotic relationship of unculturable bacteroids with plants [35].

An overview of all feasible overall reactions might furthermore be useful when studying interacting species, such as crossfeeding species, host-pathogen interactions, or multi-species communities. The possible interactions are determined by what is consumed and produced by the individual species, which is exactly the information offered by the ECMs. Indeed, knowing the capabilities of one and the incapabilities of another might lay bare dependencies on which a stable community is built.

### Methods to scale ECM computation even further

Although ECM-computation increases the size of models for which metabolic capabilities can be charted, all ECMs of genome-scale networks with thousands of reactions can still not be computed. We hope that this last scaling step can be made in the future. Even if this step cannot be made, the `hide`-method described in Box 2 enables focusing on the most relevant set of external metabolites while the steady-state constraints are still satisfied in the whole network. In Figures 3 and 5 we illustrate with an *E. coli* core model and a genome-scale *H. pylori*-model that this method can be used to obtain a much smaller set of conversions that spans all stoichiometric couplings between the user-defined external metabolites. This has not been possible with any other method. Moreover, when we focussed only on the relations between glucose, oxygen and biomass production, the `hide`-method allowed us to scale ECM computation to the genome-scale *E. coli* model (Figure 6).

ECM enumeration ignores all information about the activities of reaction rates. If the `hide`-method is used, even the consumption and production of the hidden metabolites is ignored. For example, if we hide everything but glucose, oxygen and biomass, the ECMs show that the cell is capable of converting glucose and oxygen into biomass in the reported ratios. However, we get no information about which other metabolites can be consumed and produced during this conversion. Therefore, the ECMs obtained while hiding metabolites are generally not elementally balanced, illustrated by the question marks in Figure 4**a)**. This might limit their use if one is for example interested in the thermodynamic properties of the conversion. However, for each ECM of interest, some flux distributions that lead to it can be reconstructed. These flux distributions can then be used to determine the overall conversion. The reconstruction could be done by imposing the conversion ratios from the ECM as equality constraints on the model. Then, solving an FBA-problem would give one candidate flux distribution, performing a Flux Variability Analysis [49] would give the feasible ranges of all fluxes, and it might even be possible to find all elementary pathways that lead to this ECM by computing the Elementary Flux Vectors [50, 51]. In addition, if one is particularly interested in the activities of a certain set of reactions in the conversions, this can be reported by using the `tag`-method, which is explained in detail in Box 2. In Figure 3**c)** we used the `tag`-method to highlight the use of the pyruvate-dehydrogenase reaction in the `e_coli_core`-network.

## Conclusion

In this work we presented `ecmtool`, a computational tool that calculates all overall chemical conversions that a cell might catalyse – all its metabolic capabilities – from its metabolic network alone. We hope that ECM enumeration will in the future become a standard step after metabolic network reconstruction, so that the metabolism of all known organisms will be fully characterized.

## Methods

Here, we will describe only the most important conceptual steps of the Elementary Conversion Modes computation. The method that was implemented in `ecmtool` is more elaborately described and explained in the Supplementary Information. In developing this method we strongly benefited from the pioneering work by Urbanczik and Wagner, who did not only define ECMs, but also described many of the enumeration steps. Unfortunately, their enumeration tool, implemented in a mixture of `Mathematica, Matlab`, and `C` does no longer function, but many of the ideas could be used. In the following, we will mention which conceptual steps were based on ideas from Urbanczik and Wagner, and which were added by us.

### The minimal ingredients for computing ECMs

To start the computation of ECMs we need the following ingredients

1. a stoichiometry matrix,
2. reversibility information of all reactions,
3. information on which metabolites are external or internal,
4. information on whether external metabolites can be produced, consumed or both

Our Python-implementation can automatically extract these from an `SBML`-file (Systems Biology Markup Language [33]). In the case that it is not clear whether a reaction is reversible or not, the reaction can be assumed to be reversible. Incorrectly marking a reaction as reversible can only lead to some ‘false positives’: computed ECMs that are in fact not possible, but not to ‘false negatives’. Since marking a metabolite as *internal* or *external* is sometimes ambiguous and context-dependent, we here use our own definition: a metabolite is internal whenever the steady state assumption should be met, so that its production and consumption should balance out.

### Splitting external metabolites into inputs and outputs enables ECM computation by extreme ray enumeration

The ECMs, formally defined in Box 1, can be described as the elementary vectors [?] in the space of all steady-state conversions. This space is given by:

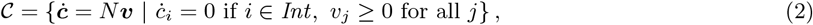

where *N* is the stoichiometry matrix, and *Int* the index set of internal metabolites. We have, for simplicity, assumed all reactions to be irreversible, but this is not necessary.

We first consider the case that the space of steady-state conversions is contained in one orthant, i.e., that for each dimension, *i*, all conversions are either nonnegative (*ċ_i_* ≥ 0), or nonpositive (*ċ_i_* ≤ 0). In that case, the elementary vectors coincide with the spanning rays: a well-defined minimal set of vectors with which we can generate the cone by taking conical combinations (weighted sums with positive weights). Enumerating the extreme rays of a polyhedral cone is a known mathematical problem described for example by Fukuda [52]. However, the set 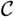 is generally not contained in one orthant, because some external metabolites can be used as an input (*ċ_i_* < 0) in some conversions, and as an output (*ċ_i_* > 0) in others. This adds ECMs that are not spanning rays of 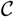, so that extreme ray enumeration is no longer enough.

We devised a new method to solve this problem: we extend the network slightly to make a new 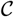 that is contained in one orthant. Let *A*_ex_ be an external metabolite that is both an input and an output. We connect *A*_ex_ to two virtual metabolites: *A*_ex,in_ and *A*_ex,out_, through two irreversible reactions: *A*_ex,in_ → *A*_ex_ and *A*_ex_ → *A*_ex,out_. Finally, we mark *A*_ex_ itself as an internal metabolite, such that it has to be kept in steady-state. As a consequence, conversions in which *A*_ex_ was produced must now produce *A*_ex,out_ to maintain the steady-state assumption. Likewise, conversions in which *A*_ex_ was consumed must now consume *A*_ex,in_. As such, all information about *A*_ex_ is stored in the production of *A*_ex,out_ and the consumption of *A*_ex,in_, while these new external metabolites can only be produced or consumed. Therefore, the new space of steady-state conversions is contained in a single orthant, so that we can proceed the ECM-computation by enumerating the spanning rays of this space. After the calculation we can then undo the splitting of metabolites so that we obtain the full set of ECMs (we prove this in Supplemental Information Section 3.3).

### Finding the ECMs is finding a generator representation of 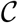

The space of steady-state conversions 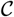 is a so-called pointed polyhedral cone. Such a cone can be described in two ways: with an *inequality representation* or with a *generator representation* [53].

The inequality representation is a set of vectors {***a***_1_,…, ***a***_*M*_} that give the bounds that constrain the cone.

All elements in the cone: 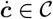, must satisfy ***a***_*i*_ · ***ċ*** ≥ 0 for all *i*. Or, as a matrix equation:

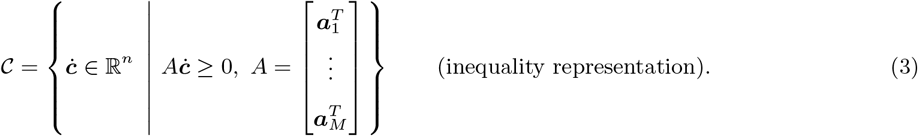

In the generator representation one gives a set of vectors, {***r***_1_,… ***r***_*K*_}, with which all elements in the cone can be generated by taking conical combinations:

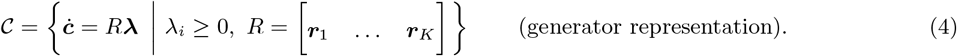

Since we have split the external metabolites into inputs and outputs before, computing the ECMs now amounts to obtaining a minimal generator representation of 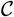, because the generators, ***r***_*i*_, are then precisely the ECMs.

### The main computation step: impose equality constraints on a large set of generators

Following Urbanczik et al. [31], we will start the computation with a cone that is too large, but for which we already have a generator representation. To be precise, we will start with the cone generated by the columns of the stoichiometry matrix:

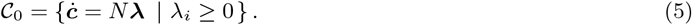

This cone is the space of all conversions that can result from combinations of reactions of the metabolic network, no matter if these conversions meet the steady state requirement or not. Therefore, this cone does contain the steady state conversion cone, 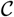, because it contains all possible conversions in steady-state. However, to get a good description of 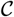, we should still impose the steady state constraint. To compute the ECMs, we should therefore keep track of how our set of generators changes while we impose the steady-state equalities *ċ_i_* = 0 for each internal metabolite.

Concluding: we start with a set of generators of the cone 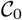, we impose the set of equalities given by *ċ_i_* = 0, and are then interested in the generators of the resulting cone. We have implemented two methods for this main part of the computation: an *indirect method* which was extended from suggestions in literature [31, 52], and a *direct method* which we developed ourselves. These methods are described elaborately in SI Sections 7 and 8, but we will also shortly explain both below. We chose to implement both methods because their merits complement each other. The indirect method is fast on small- to medium-scale networks, and might therefore be preferred over the direct method. The method is called indirect, because it first computes a large intermediate result which is then used to compute the ECMs. However, the intermediate result might be much larger than the final result, so that the indirect method can run into memory issues while calculating the intermediate result, even though the final result is not that large. Our newly developed direct method performs better on larger networks, and especially when many metabolites are hidden using the `hide`-method, because it avoids such large intermediate results.

### The indirect method

As we will explain below, the indirect method twice uses the Double Description (DD) method [54, 40]. The DD-method computes a minimal set of generators from an inequality representation of a cone. This part of the computation is done using `polco` [55].

Although our actual starting point is a generator representation of 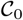, it is useful for now to imagine that we already have an inequality representation of 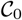. We will later explain how we obtain this representation. This inequality representation would be a set of vectors ***h***_1_,… ***h**_M_*, such that

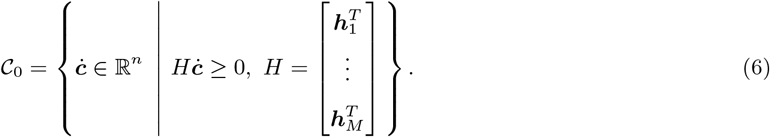

Given this representation, it is easy to impose a steady-state constraint *ċ_i_* = 0, by adding the elementary unit vector *ê_i_* = [0,⋯, 0,1, 0,⋯, 0]^*T*^ both positively and negatively to the set of inequalities. This enforces 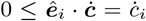, and 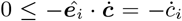, such that we have actually imposed *ċ_i_* = 0. In our implementation, we have sped up the computation by imposing the steady-state constraint through removing the *i*^th^ column from the inequality constraint matrix *H*. We prove in SI Section 7.3.1 that this is equivalent. Removing these columns can make many of the rows in the constraint matrix redundant. For this, we have developed a redundancy removal algorithm that minimizes the size of the inequality constraint matrix, see SI Sections 5.6 and 7.3.3.

Comparing (2) and (6), we see that by imposing these steady state constraints for all internal metabolites we go from an inequality representation for 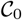 to an inequality representation of the cone of steady-state conversions 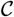. From this, we can use the Double Description method to compute a minimal set of generators for this cone, yielding the ECMs.

It remains to be shown how we obtain an inequality representation of 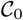 from the generator representation we start with. For this, we use that 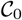 has a dual cone associated to it: 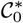, which has two important properties (see SI 1.3 for more information and explanation):

1. the dual of the dual cone is again the cone: 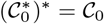,
2. the vectors in the generator representation of a cone form an inequality representation of the dual cone, and vice versa: 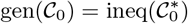.

Our computation starts from a generator representation of 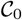 (5), but by property 2 this is also an inequality representation of its dual 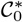. By applying the Double Description method on this inequality representation, we can find a generator representation of 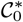. This generator representation is, again by property 2, an inequality representation of the dual of this dual cone. By property 1, we have thus obtained an inequality representation of 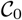, which was exactly what we needed. The steady-state constraints can now be imposed and the ECMs computed, as explained above.

Note that this indirect method heavily relies on the Double Description method. We found that polco [55] functions well and is reasonably fast, but can run into memory issues when the networks for which we try to compute the ECMs get too large. We found that these memory issues were caused by the size of the inequality representation needed to describe 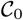, i.e., the issues arise in the first application of the DD-method. This therefore causes a computational limitation even though the generator representation of 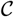 (which we are eventually after) can be much smaller. This lack of control of the size of our intermediate results forms an important disadvantage of the indirect method. Therefore, we developed the direct method for the computation of ECMs for larger networks.

### The direct method

Just as the indirect method, the direct method starts with the cone 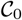 introduced in (5), generated by the columns of the stoichiometry matrix *N*. We collect these generators in a matrix *R*^(0)^. Then, we iteratively impose the steady-state constraints, *ċ_i_* = 0, for all internal metabolites *i*. Imposing such a steady-state constraint means that we take the intersection of the cone 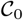 with the hyperplane *ċ_i_* = 0. The intersection is again a cone, called 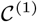, which is generated by a new set of generators that we collect in a matrix *R*^(1)^. Proceeding with *R*^(1)^ and imposing more steady-state constraints, we will eventually end up with a set of generators for the steady-state conversion cone 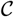.

One such iteration thus starts with a set of generators of *C*^(*i*−1)^, collected in *R*^(*i*−1)^. Now, we distribute these generators in three groups: a plus-group, a zero-group and a minus-group, depending on if the generators have *ċ_i_* > 0, *ċ_i_* = 0, or *ċ_i_* < 0, respectively. The generators in the plus- and minus-groups do not satisfy the steady-state constraint, and should therefore be dropped. However, each combination of a plus-generator with a minus-generator can provide a candidate generator that does satisfy *ċ_i_* = 0. These candidates, combined with the generators that were already in the zero-group, must contain all generators of 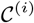.

However, when we combine all generators from the plus-group with the minus-group to create new generators, we will not get a *minimal* set of generators. In other words, the cone 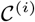 could also be generated by a smaller number of generators. This might not seem like a large problem, but the number of unnecessary generators (also called redundant generators) grows exponentially with the number of iterations, quickly causing computational infeasibility. Therefore, we have developed an *adjacency test*. This test determines for each candidate, i.e., an appropriate combination of a plus-generator with a minus-generator, if it is redundant. It does so by checking if the candidate can be written as a combination of other generators. If so, then the candidate is redundant, and should be left out of *R*^(*i*)^. This test is implemented by performing a linear optimization for each candidate. In SI Section 8.1 we have added figures to explain our method. There, we also elaborate on the linear optimization, and explain how we optimized it to be fast enough.

Although performing many linear optimizations is in principle a very slow process, there is an important advantage: the different optimizations can be done completely independently. Therefore, we were able to parallelise this direct method so that it can now be run on large computation clusters.

### Network compression facilitates ECM computation on large networks

ECM theory focuses on the overall conversions between external metabolites, instead of on how these conversions come about internally. This distinction can be exploited to simplify the network even before we start the main computation steps described above. We have implemented several compression steps that together bring large networks back to a workable size. Most of these compression steps were suggested by Urbanczik et al. [31]. We have added the removal of cycles, the removal of redundant reactions, and part of the removal of infeasible reactions. In the Supplementary Information (Section 5) we provide proofs and more extensive explanations.

#### Infeasible reactions can be removed

The flux vectors that give rise to the ECMs should satisfy the steady state and the irreversibility constraints. If we can prove that a reaction can never be active in a solution that meets both of these constraints, then this reaction can be safely removed. In principle, the feasibility of a reaction can be tested by running a linear optimisation: for reaction *i* we would maximise *v_i_* such that *v_j_* ≥ 0 for irreversible reactions *j*, and such that *N_int**v**_* = 0, where *N_int_* is the part of the stoichiometry matrix corresponding to internal metabolites. If the optimal solution does not give *v_i_* strictly larger than zero, then reaction i is infeasible and can be removed from the network. In Section 5.1 of the SI we describe a computationally efficient way of achieving the same.

#### Redundant reactions can be deleted

We can delete redundant reactions; a reaction is called redundant if it can be written as a conical combination of other reactions. Because the function of these reactions can always be replaced by the combination of the other reactions, it does not add functionality to the network and can therefore be removed. Redundant reactions in systems with fewer than about 10,000 reactions can be removed using a program called `redund` from *lrslib* [56], so that this suffices during this compression step. As we have mentioned above, we also apply redundancy removal during both the direct and the indirect method, and here the number of reactions can become much larger than 10,000. This is the reason that we also developed our own parallelisable redundancy test (see SI 5.6 for an explanation).

#### Reversible reactions can be used to cancel a reaction and a metabolite

Each reversible reaction can be used to cancel itself and one metabolite it connects to. Say that a reversible reaction, *R*_1_, produces an internal metabolite *A*, and say that there are several other reactions producing or consuming *A*. We prove in Section 5.3 of the SI that we can, without changing the ECMs of the network, add or subtract reaction *R*_1_ to these other reactions such that the production or consumption of *A* is cancelled. After doing this for all reactions connected to *A, R*_1_ is the only reaction left that produces *A*. This implies that no reaction flux is possible through *R*_1_ in a steady-state solution, because the production of *A* cannot be compensated by another reaction. Therefore, we can delete both *R*_1_ and *A* from the network without affecting the ECM-results.

#### Dead-end metabolites and connecting reactions can be deleted

Sometimes an internal metabolite can only be produced and not consumed, or vice versa. In this case, the reaction flux through the reactions connected to this metabolite has to be zero in any steady-state solution. Therefore, we can delete the metabolite and all connecting reactions without affecting the set of ECMs.

#### Reactions with a unique function can be used to cancel a reaction and a metabolite

Say that we have a reaction *R*_1_ which is the sole reaction that produces a metabolite *A*, but that there are several reactions that consume *A*. Then, again without affecting the set of ECMs, we can add *R*_1_ to these consuming reactions such that the consumption of *A* is exactly cancelled. The reaction *R*_1_ is now the only reaction left that produces *A*, and can therefore not be active in a steady-state solution. We can thus cancel both *R*_1_ and *A*.

#### Cycles of *k* reactions can cancel *k* – 1 reactions and metabolites

A cycle is a set of reactions that can be used in a certain ratio such that nothing is produced nor consumed. Say that the reactions *R*_1_,…, *R_k_* form a cycle, so that with appropriate weights λ_*i*_ we have 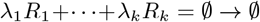. In addition, say that *R*_1_ produces an internal metabolite *A*. In Section 5.5 of the SI we show that we can now use a trick similar to what we used with the reversible reactions. We use λ_1_*R*_1_ as the forward reaction, and λ_2_*R*_2_ + ··· + λ_*k*_*R_k_* as the backward reaction to cancel the production and consumption of *A*. After doing this, *R*_1_ will again be the only reaction producing *A*, so that we can delete both *R*_1_ and *A* from the network. Since we compensated for the action of *R*_1_ in the rest of the network, we will be left with a cycle using the reactions *R*_2_,…, *R_k_*, on which we can use the same trick again. In this way, we can delete *k* – 1 reactions and *k* – 1 metabolites.

##### Box 3: A worked-out example of ECM enumeration with the direct method

To show how ECM enumeration works in practice, we will here work out some steps of the computation of ECMs for the network given by the following stoichiometry matrix:

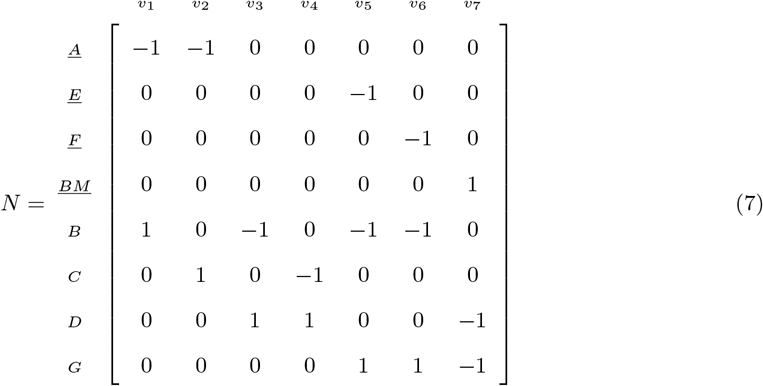

which is also shown as the first network of Figure 7**a)**. All reactions are assumed irreversible, external metabolites *A, E* and *F* can only be used as inputs, and *BM* can only be used as an output. For the enumeration, we will use the direct intersection method, and we will not apply any of the network compression steps (examples of these steps can be found in SI 5).

**Figure 7:**
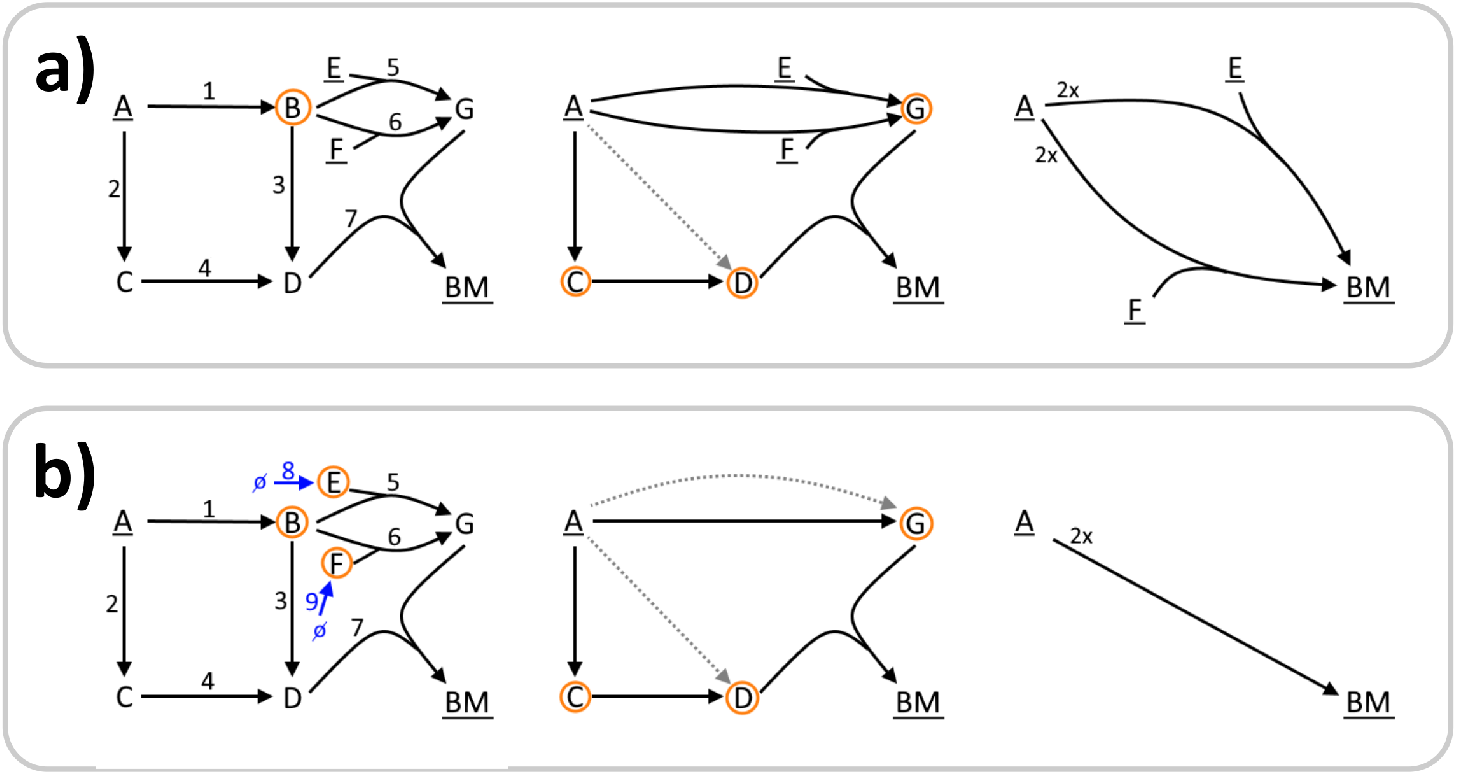
A worked-out example of ECM enumeration on a small network. All steps are described in the main text. Metabolites that are underlined are marked as external. The metabolites for which we impose the steady-state constraint in the next step are circled. Dotted arrows indicate conversions that were found to be redundant, and are thus deleted.

The stoichiometry matrix gives a list of generators that generates all conversions before we have imposed the steady-state constraints: *R*^(0)^ = *N*. On this collection of generators, we impose the steady state constraint for metabolite *B*, i.e., 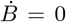. In the stoichiometry matrix we can see that there are three reactions, *v*_2_, *v*_4_, *v*_7_, that do not produce or consume *B*, and therefore already satisfy this constraint. Of the other reactions, *v*_1_ produces *B* and *v*_3_, *v*_5_, *v*_6_ consume *B*. Each pair of a producing and a consuming reaction generates a candidate that satisfies the steady state constraint, so this gives us 1 **×** 3 = 3 candidates:

- *v*_1_ + *v*_5_: <u>*A*</u> + <u>*E*</u> → *G*,
- *v*_1_ + *v*_6_: *<u>A</u>* + *<u>F</u>* → *G*,
- *v*_1_ + *v*_3_: *<u>A</u>* → *<u>D</u>*.

All candidates are tested for redundancy by the adjacency test described in SI Section 8.1. This test indicates if the candidate can be written as a positive combination of already existing reactions. The first two reactions are non-redundant, and thus added to the next list of generators, but the third reaction can be written as a sum of *v*_2_ and *v*_4_, and is therefore not added. We get:

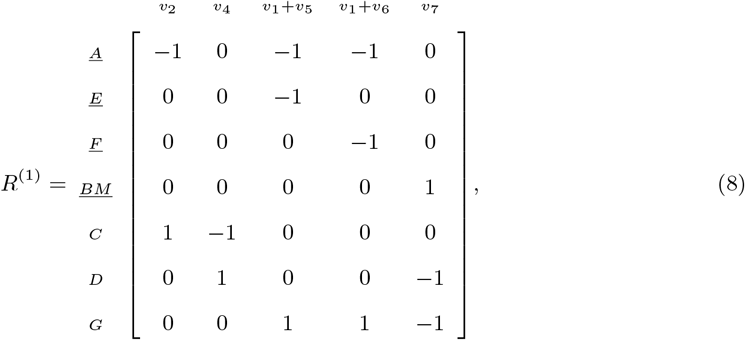

which is depicted as the second network in Figure 7**a)**.

This process is then repeated for internal metabolites *C, D* and *G*, eventually giving

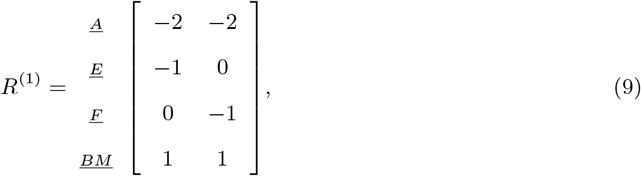

containing all ECMs, namely 2 *<u>A</u>* + *<u>E</u>* → <u>*BM*</u>, and 2*<u>A</u>* + *<u>F</u>* → *<u>BM</u>*.

In Figure 7**b)** we illustrate the ECM enumeration when we use the `hide`-method to ignore the consumption of *E* and *F*. Hiding these metabolites is done by extending the metabolic network with reactions that create *E* and *F* from nothing, and marking the metabolites as internal. We thus get:

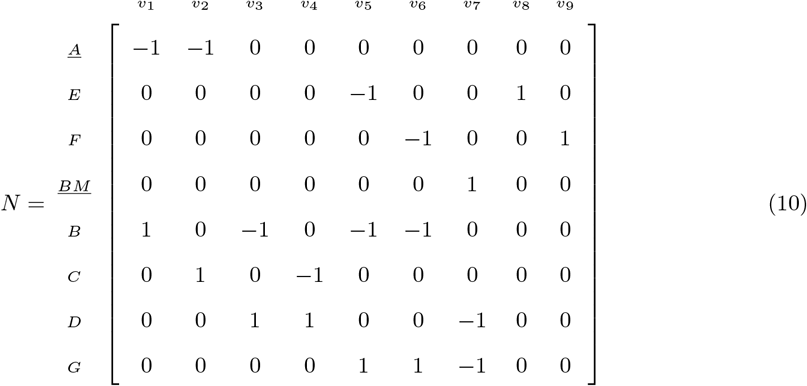

When we now start by imposing the steady state constraints for metabolite *E*, we see that only *v*_5_ and *v*_8_ do not satisfy this constraint. Combining these reactions gives the candidate: *v*_5_ + *v*_8_: *B* → *G*, which is added to the new list of generators. When we then impose the steady state constraint for metabolite *F*, we get the same candidate *v*_6_ + *v*_9_: *B* → *G*, but since this is not a new conversion it is not added to the list of generators. It can thus be seen that hiding metabolites *E* and *F* immediately reduces the computational complexity, because now only one reaction from *B* to *G* remains, while without the `hide`-method there were two such reactions. Moreover, after imposing the remaining steady state constraints, we find only one ECM: 2 *A* + ?? → *G*, where the question marks indicate that we do not know whether more metabolites are consumed because this information is hidden.

Although it would not give problems in this example, we can in general not hide a metabolite by simply removing it from the network. This is because information about whether the metabolite can be used as an input, as an output, or both, would be lost from the computation. With the current method, this information is stored in the directionality of the added reaction.

### The ECM-computation was implemented in Python

We implemented our algorithms in a publicly available Python-program called `ecmtool`. It is freely available on GitHub at https://github.com/SystemsBioinformatics/ecmtool, and can additionally be installed through the Python package manager `pip`. The direct and indirect computation method are both available within the program. A manual is available as Section 11 of the SI, and some worked-out examples are provided as supplementary files.

## Supporting information

Theoretical background ecmtool-methods, including user manual

## Author’s contributions

Conceptualization, D.H.dG. and T.J.C.; Software, T.J.C., E.B.B. and D.H.dG.; Validation, T.J.C., E.B.B. and D.H.dG.; Formal Analysis, T.J.C., E.B.B., R.P. and D.H.dG.; Resources, F.J.B. and B.T.; Writing - Original Draft, D.H.dG.; Writing - Review & Editing, T.J.C., R.P., F.J.B., B.T., and D.H.dG.; Visualization, T.J.C. and D.H.dG; Supervision, R.P., F.J.B., B.T. and D.H.dG; Funding Acquisition, F.J.B., B.T.

## Acknowledgements

We thank Brett Olivier and Leen Stougie for helpful discussions and Carolin Schulte for showing us the scope of the potential applications of `ecmtool`. Bas Teusink and Daan de Groot were supported by NWO VICI grant 865.14.005 (https://www.nwo.nl/).

## Supplementary Files

1. Document with supplementary figure, and detailed theoretical background for all methods used in `ecmtool`
2. User manual
3. Matlab-scripts used for validation of ECMs
4. Python-script used for creating subnetworks of the e_coli_core-network
5. R-scripts used for clustering ECMs

