## Supplementary material for "Unlocking Elementary Conversion Modes: ecmtool unveils all capabilities of metabolic networks": Theoretical background ecmtool-methods, including user manual

#### Supplementary figures

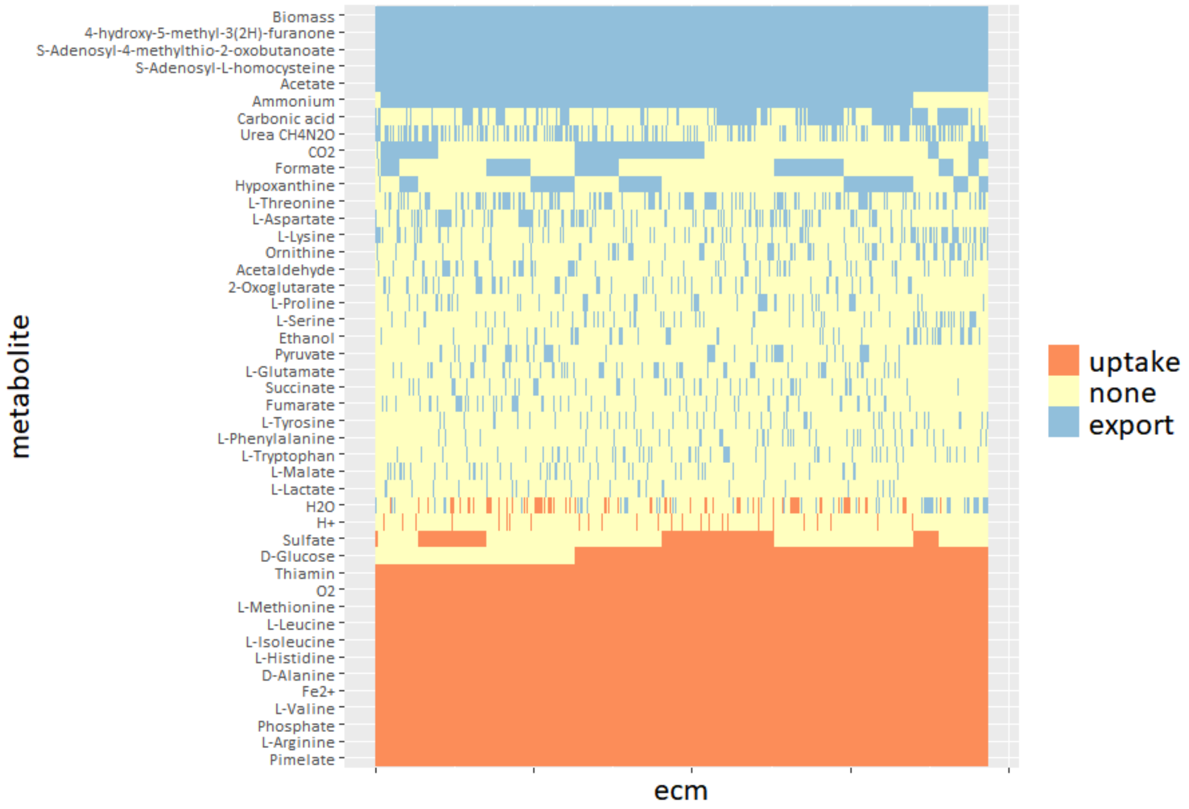

Figure S1: **All Elementary Conversion Modes for the iIT_341-model of *Helicobacter pylori* could be computed.** We show the ECMs that support growth on a defined medium proposed by the model developers (see minII in [?]). For visualization purposes, we only show which metabolites are consumed or produced rather than their exact stoichiometric coefficient. In addition, we show only the ECMs that in which L-alanine was not taken up, and in which acetate and biomass was produced, because the full set of 874 236 ECMs was simply too large to show. After these simplifications, there were still 7740 unique ECMs. The full set of ECMs is available as a supplementary file.

#### Notation and conventions

In the following, we will denote matrices by capital letters ($A$), and vectors by bold lowercase letters ($\boldsymbol{v}$) (or bold uppercase letters if the vectors are rows or columns from a matrix). The requirement that all components of a vector $\boldsymbol{v} \in \mathbb{R}^n$ are nonnegative, $v_i \geq 0,\ i \in \{1, ..., n\}$, will be written as $\boldsymbol{v} \geq \mathbf{0}$. Time derivatives are indicated by a dot, $\dot{c} = \frac{dc}{dt}$. All single vectors in this work denote column vectors, unless otherwise stated. The $i^{\text{th}}$ row of a matrix $A$ is indicated as $\boldsymbol{A}_{i\bullet}$. Column vectors of $A$ are sometimes denoted as $\boldsymbol{A}_{\bullet j}$.

In biochemical networks, an underlined metabolite ($\underline{\mathrm{S}}$) is assumed to be 'external', which here means that the steady-state constraint does not have to be satisfied for that metabolite. To simplify the following analysis, we will assume all reactions to be irreversible, unless explicitly stated otherwise. We can do this without loss of generality since all reversible reactions can be decomposed into an irreversible forward and an irreversible backward reaction.

### 1 Background information on polyhedral computation

We will shortly review the concepts from linear algebra that we will need for the definition and enumeration of ECMs (see for example [**?**]).

#### 1.1 Inequality and generator descriptions of polyhedral cones

A subset $S$ of $\mathbb{R}^d$ is called a *convex cone* if any conical combination of two vectors $\boldsymbol{v}, \boldsymbol{w} \in S$ is still a vector in $S$, i.e., for all $\alpha, \beta \in \mathbb{R}_{\geq 0}$ we have $\alpha \boldsymbol{v} + \beta \boldsymbol{w} \in S$. $S$ is called a *polyhedral convex cone* if there is some constraint matrix $A$ such that

$$S = \left\{ \boldsymbol{x} \in \mathbb{R}^d \mid A\boldsymbol{x} \geq \boldsymbol{0} \right\}, \qquad \text{(inequality representation)}. \tag{1}$$

This way of describing $S$ is called the inequality-, or $H$-representation. The Minkowski-Weyl theorem tells us that every polyhedral cone can be generated by taking conical combinations of a finite set of vectors, $\boldsymbol{r_1}, \ldots, \boldsymbol{r_n} \in \mathbb{R}^d$. If we collect these vectors as the columns of a matrix $R$, this gives us a second representation of $S$ called the generator-, or the $V$-representation:

$$S = \{ \boldsymbol{x} = R\boldsymbol{\lambda} \mid \lambda_i \geq 0 \}, \qquad \text{(generator representation)}. \tag{2}$$

Such representations of $S$ are denoted by ineq($S$) and gen($S$), respectively.

#### 1.2 Pointedness

Vectors $\boldsymbol{v} \in S$ for which $-\boldsymbol{v} \in S$ as well, are called linealities. The space that comprises all such vectors is called the lineality space. Since by the definition of $S$, we would have $A\boldsymbol{v} \geq \boldsymbol{0}$ and $A(-\boldsymbol{v}) = -A\boldsymbol{v} \geq \boldsymbol{0}$, we can describe the lineality space as the null-space of the constraint matrix that was used in the inequality representation of $S$, i.e.,

$$\text{Lin}(S) = \left\{ \boldsymbol{x} \in \mathbb{R}^d \mid A\boldsymbol{x} = \boldsymbol{0} \right\}. \tag{3}$$

A polyhedral cone is called *pointed* if its lineality space contains only the zero vector. A pointed cone has a unique generator representation.

In one of the steps of ECM-enumeration, one can encounter a non-pointed polyhedral cone. The computation, however, can only proceed with a pointed cone. In such cases we can use that any cone is the direct sum of its lineality space and a pointed cone [?]. To be precise, for any cone $S$, we have

$$S = \mathrm{Lin}(S) \oplus (S \cap \mathrm{Lin}(S)^{\perp}). \tag{4}$$

The pointed part of the polyhedral cone is thus given by all vectors in the cone that are perpendicular to the lineality space. Using a set of basis vectors $\boldsymbol{n}_1, \ldots, \boldsymbol{n}_k$ of the lineality space $\mathrm{Lin}(S)$, we can even get an inequality representations for the two parts

$$S = \left\{\boldsymbol{x} \in \mathbb{R}^d \mid A\boldsymbol{x} = \boldsymbol{0}\right\} \oplus \left\{\boldsymbol{x} \in \mathbb{R}^d \mid A\boldsymbol{x} \geq \boldsymbol{0}, \quad N\boldsymbol{x} = \boldsymbol{0}\right\}, \tag{5}$$

where $N = \begin{bmatrix} \boldsymbol{n}_1 & \cdots & \boldsymbol{n}_k \end{bmatrix}^T$.

#### 1.3 Dual cones

Let $S$ be a polyhedral cone in $\mathbb{R}^d$. Its *dual cone* $S^*$ is defined as

$$S^* = \{\boldsymbol{u} \in \mathbb{R}^d \mid \boldsymbol{u} \cdot \boldsymbol{v} \geq 0, \boldsymbol{v} \in S\}. \tag{6}$$

The dual of the dual of a convex polyhedral cone is equal to itself if $S$ is convex and closed.

In Figure S2 we illustrate an important property of dual cones: a generating set of $C$, $\mathrm{gen}(C)$, forms the inequalities representation of $C^*$, $\mathrm{ineq}(C^*)$, and vice versa [?, ?]:

$$\mathrm{gen}(C) = \mathrm{ineq}(C^*), \tag{7}$$

$$\mathrm{ineq}(C) = \mathrm{gen}(C^*). \tag{8}$$

#### 1.4 Adjacent rays

Assume that we have an $m \times n$ inequality representation $A$ of the polyhedral cone

$$\mathcal{C} = \{\boldsymbol{x} \in \mathbb{R}^n : A\boldsymbol{x} \geq \boldsymbol{0}\}$$

and corresponding ray representation $R$:

$$\mathcal{C} = \{\boldsymbol{x} \in \mathbb{R}^n : \boldsymbol{x} = R\boldsymbol{\lambda}, \boldsymbol{\lambda} \geq \boldsymbol{0}\}.$$

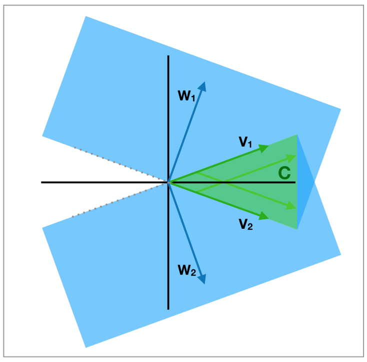

(a)

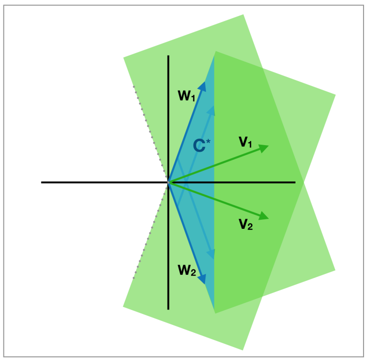

(b)

Figure S2: **Cones are spanned by inequality constraints of their dual.** (a) A 2-dimensional cone $\mathcal{C}$ (green) spanned by generating vectors $v_1$ and $v_2$, meaning that any nonnegative linear combination of them (light-green arrows) is also in $\mathcal{C}$. $\mathcal{C}$ can additionally be described by the intersection of the two halfspaces $\boldsymbol{w}_1 \cdot \boldsymbol{x} \geq \boldsymbol{0}$ and $\boldsymbol{w}_2 \cdot \boldsymbol{x} \geq \boldsymbol{0}$ (blue). Here, $\boldsymbol{w}_1$ and $\boldsymbol{w}_2$ form the inequality representation of $C$. (b) The dual of $\mathcal{C}$, $\mathcal{C}^*$ (blue), is spanned by the inequality representation vectors of $\mathcal{C}$. Thus, we can see that the inequality representation of a cone forms generating vectors of its dual, and its generating vectors form the inequality representation of its dual. Both cones, $\mathcal{C}$ and $\mathcal{C}^*$, extend to infinity; only a section is drawn.

For each extreme ray $\boldsymbol{r}_j$ of the cone (i.e., the $j$-th column of $R$), define the *zero set*

$$Z(\boldsymbol{r}_j) = \{i : \boldsymbol{A}_{\boldsymbol{i}\bullet}\boldsymbol{r}_j = 0\}.$$

Two distinct extreme rays $\boldsymbol{r}_{j_+}, \boldsymbol{r}_{j_-} \in R$ are *adjacent* if for any extreme ray $\boldsymbol{r}_j$, we have

$$Z(\boldsymbol{r}_{j_+}) \cap Z(\boldsymbol{r}_{j_-}) \subseteq Z(\boldsymbol{r}_j) \implies \boldsymbol{r}_j \sim \boldsymbol{r}_{j_+} \text{ or } \boldsymbol{r}_j \sim \boldsymbol{r}_{j_-},$$

where $\sim$ means that the two vectors are scalar multiples of each other. In other words, two vectors $\boldsymbol{r}_{j_+}$ and $\boldsymbol{r}_{j_-}$ are adjacent if there are no other extreme rays that satisfy the constraints with equality that are satisfied with equality by both $\boldsymbol{r}_{j_+}$ and $\boldsymbol{r}_{j_-}$ from $A$.

Geometrically, two distinct extreme rays are adjacent if the minimal face of $C$ containing both contains no other extreme rays. For more details, see Proposition 7 in [**?**].

#### 1.5 The double description method

The Double Description method is an algorithm originally suggested by Motzkin et al. [**?**] to translate between the two representations of a cone. For a more comprehensive description and proofs of the propositions, see [**?**].

A pair $(A, R)$ is called a **double description pair** or DD pair if $A$ is an inequality representation and $R$ is a generator representation of the same cone. That is,

$$\{\boldsymbol{x} \in \mathbb{R}^n : A\boldsymbol{x} \geq \mathbf{0}\} = \{\boldsymbol{x} \in \mathbb{R}^n : \boldsymbol{x} = R\boldsymbol{\lambda}, \boldsymbol{\lambda} \geq \mathbf{0}\}.$$

The following proposition shows that an algorithm that can translate in one direction automatically also solves the inverse direction.

**Proposition 1.** *$(A, R)$ is a DD pair if and only if $(R^T, A^T)$ is a DD pair.*

The double description method provides an algorithm to find an $R$ based on a given $A$ such that $(A, R)$ is a DD pair. As we described in the previous section, $R^T$ is an inequality representation for the dual cone of the cone represented by the inequalities in $A$. If we want to find a DD pair starting from a given $R$, we can take $R^T$ and treat it as the inequality representation of a cone, then apply the double description method to find $A^T$ (hence giving $(A, R)$).

The core of the double description algorithm is an incremental procedure. We start with an $m \times n$ matrix $A$, giving the inequality representation of a cone that we will denote by $P(A)$. Let $K \subset \{1, \ldots, m\}$ be a subset of the row indices of $A$ and $A_K$ the submatrix consisting of the corresponding rows. Suppose we already have a ray representation for the cone $P(A_K)$, i.e., we have a DD pair $(A_K, R_K)$. To perform the incremental step, select any index $i$ not in $K$; we will denote the corresponding row by $\boldsymbol{a}_i$. We will add $\boldsymbol{a}_i$ to $A_K$ to form $A_{K\cup\{i\}}$ and use $R_K$ to construct a DD pair $(A_{K\cup\{i\}}, R_{K\cup\{i\}})$.

First, we partition the column index set $J$ of $R_K$ into three parts:

$$J^+ = \{j \in J : \boldsymbol{a}_i R_{\bullet j} > 0\},$$
$$J^0 = \{j \in J : \boldsymbol{a}_i R_{\bullet j} = 0\},$$
$$J^- = \{j \in J : \boldsymbol{a}_i R_{\bullet j} < 0\}.$$

To make the matrix $R_{K\cup\{i\}}$ we keep all the columns $R_{\bullet j}$ from $R_K$ with $j \in J^+$ or $j \in J^0$. These rays satisfy the inequality given by the row $\boldsymbol{a}_i$ already: $\boldsymbol{a}_i R_{\bullet j} \geq 0$ and so they are in $P(A_{K\cup\{i\}})$ unchanged. The columns corresponding to $J^-$ are not in the new cone, but they give rise to new generating rays in the following way.

Let $\boldsymbol{r}_{jj'} = (\boldsymbol{a}_i R_{\bullet j})R_{\bullet j'} - (\boldsymbol{a}_i R_{\bullet j'})R_{\bullet j}$ for each $(j, j') \in J^+ \times J^-$. Note that $\boldsymbol{a}_i \boldsymbol{r}_{jj'} = 0$ for each choice of $(j, j')$. These columns are also added to $R_{K\cup\{i\}}$.

To sum up, $R_{K\cup\{i\}}$ is the $d \times |J'|$ matrix with columns given by the index set $J' = J^+ \cup J^0 \cup (J^+ \times J^-)$,

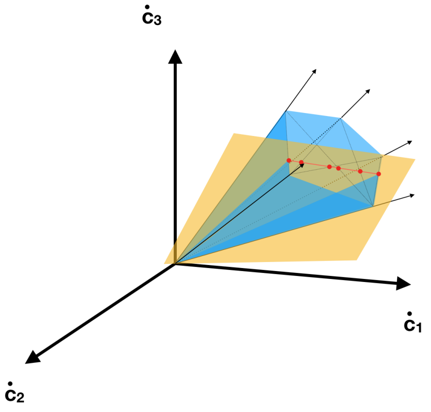

Figure S3: A cone (blue) spanned by generators (black) is being intersected with an inequality constraint (orange). All generators on one side of the constraint can remain, and those on the other side are replaced by the non-redundant the red vectors. The red vectors are conical combinations of each pair of extreme rays on both sides of the halfspace, such that the red vector exactly saturates the constraint. This process is repeated for all inequality constraints, until only the generating set of the final cone remains.

where $\boldsymbol{r}_{jj'} = (\boldsymbol{a}_i R_{\bullet j}) R_{\bullet j'} - (\boldsymbol{a}_i R_{\bullet j'}) R_{\bullet j}$. In Figure S3, one intersection with an inequality constraint is illustrated.

**Proposition 2.** $(A_{K\cup i}, R_{K\cup\{i\}})$ *is a DD pair.*

Repeating this step, eventually all the rows of $A$ will be included, giving a DD pair $(A, R)$ as intended. What is left is to find is a starting point. A good option is starting with $d$ linearly independent rows of $A$ to form $A_{K_0}$. Then we can find an inverse matrix, and $A_{K_0}\boldsymbol{x} = \boldsymbol{\lambda} \geq \boldsymbol{0}$ implies that $\boldsymbol{x} = A_{K_0}^{-1}\boldsymbol{\lambda}$, so that $(A_{K_0}, A_{K_0}^{-1})$ is a DD pair.

##### 1.5.1 Redundant rays

Although the Double Description method as described above gives a ray representation $R$, it is not necessarily a *minimal* representation: there can be redundant columns in $R$ that are not extreme rays of the cone. In fact, the number of unnecessary rays in practice increases very fast and goes beyond any tractable limit [?].

One possible solution is to discard all redundant vectors, preferably between each step of the Double Description method. This can be done, for example, with the program `redund` from `lrslib` [?], which looks for redundant vectors by trying to find matching conic combinations of other rays (see also Section 5.6). However, this quickly becomes computationally intensive for large sets of vectors.

Alternatively, for each $\boldsymbol{r}_{jj'}$ that we created with $(j, j') \in J^+ \times J^-$, we can test the originating rays for adjacency, which we defined above in Section 1.4. It turns out the new ray $\boldsymbol{r}_{jj'}$ is redundant if and only if its parent rays were not adjacent (Lemma 8 in [?]). The Double Description method, including an adjacency test, is efficiently implemented in `Polco` [?]. In Section 8.1.2 we will return to this adjacency test, because we need an adapted version for our direct intersection method.

#### 2 Defining Elementary Conversion Modes

A metabolic network with $r$ reactions and $m$ metabolites can be summarized in an $m \times r$ stoichiometry matrix $N$. The entries in this matrix denote how much of which metabolite (rows) are used in each reaction (columns). The stoichiometric matrix can be multiplied by a vector of reaction rates, a *flux vector* $\boldsymbol{v}$, to yield the net production and consumption of metabolites by this combination of reaction rates: $\dot{\boldsymbol{c}} = N\boldsymbol{v}$. We will call $\dot{\boldsymbol{c}}$ a *conversion*. We can assume without loss of generality that all reactions in the metabolic network are irreversible, meaning that $\boldsymbol{v} \geq \mathbf{0}$, and we will do so unless otherwise stated.

We often impose a steady-state constraint on a part of the metabolites, enforcing that the net production of these metabolites is zero. In the following, we will call the metabolites for which we impose the steady-state *internal metabolites*, and we collect their indices in an index set Int. We can now define two important sets, the *steady-state flux cone*:

$$\mathcal{F} = \{\boldsymbol{v} \in \mathbb{R}^r \mid N_{\text{Int}}\boldsymbol{v} = \mathbf{0},\ v_i \geq 0 \text{ if reaction } i \text{ irreversible}\}, \tag{9}$$

and the *steady-state conversion cone*:

$$\mathcal{C} = \{\dot{\boldsymbol{c}} = N\boldsymbol{v} \mid N_{\text{Int}}\boldsymbol{v} = \mathbf{0},\ v_i \geq 0 \text{ if reaction } i \text{ irreversible}\}. \tag{10}$$

Both of these sets are convex polyhedral cones and thus have both an inequality representation, and a generator representation. The definitions of Elementary Flux Modes and Elementary Conversion Modes are closely related to the generator representations of these cones. We here give a definition of EFMs that is not conventional, but which clarifies the analogy with ECMs better.

**Definition 1.** *The set of* Elementary Flux Modes (EFMs) *is the minimal set* $\{efm_1, \ldots, efm_K\} \subset \mathcal{F}$ *of flux vectors such that each steady-state flux vector can be written as a positive sum of EFMs, without the flux of any reaction being canceled in that sum.*

The first part of this definition implies that all vectors in the generator description of the steady-state flux cone $\mathcal{F}$ are EFMs. However, in case that we have reversible reactions, these do not form the complete set of EFMs. Rather, we should add conical combinations of these generators that make an additional reaction rate equal to zero. The complete set of EFMs may be found by taking the union of all generator sets of the intersections of $\mathcal{F}$ with the orthants. In practice, it is easier to just split all reversible reactions in a forward and a backward reaction. As such, no reaction can be canceled and the EFMs are exactly the generators of the new flux cone $\mathcal{F}$.

**Definition 2.** *The set of* Elementary Conversion Modes (ECMs) *is the minimal set of conversions* $\{ecm_1, \ldots, ecm_L\} \subset \mathcal{C}$ *such that each steady-state conversion can be written as a positive sum of ECMs, without the production of any external metabolite being canceled in that sum.*

This definition is further explained and illustrated in Box 1 of the main text. It is important to note that a reasoning can be used here that is similar to what was used for the EFMs above. That is, there are two ways of calculating ECMs as the generators of a convex polyhedral cone. First, we can take the union of all generators of the cones obtained by intersecting $\mathcal{C}$ with the orthants. Second, we can split all metabolites that can be produced and consumed into two virtual metabolites: one that is consumed, and one that is produced. We will take the latter approach in this work.

### 3 Pre-processing of the metabolic networks

To facilitate the enumeration of ECMs, `ecmtool` goes through some pre-processing steps after reading in a metabolic network. We will here review these steps shortly. Although `ecmtool` works well with most models in the `SBML`-format, we recommend the user to always review the success of these preprocessing steps. We have added arguments called `--print_metabolites` and `--print_reactions` to facilitate this review. With the printed lists of parsed metabolites and reactions, the user can check if the correct metabolites are marked as external, and if their directionality is as intended.

#### 3.1 Deleting exchange reactions and determining directionality of external metabolites

`Ecmtool` first detects external metabolites by using functionalities from the `cbmpy`-package (`http://cbmpy.sourceforge.net/`). Metabolites are marked external when their metabolite-id has a specific suffix (the suffix for the external compartment is set to `e`, but this can be changed with the argument `--external_compartment`), or when the metabolite is attached to a 'dead-end reaction', which is a reaction that involves only one metabolite.

These dead-end reactions are virtual reactions called exchange reactions. Exchange reactions are present in models to allow the steady-state assumption to hold even for the external metabolites, since these external metabolites are often assumed fixed. In the case of ECMs, however, we are interested in the production and consumption of these external metabolites, and will therefore delete all exchange reactions.

Based on the directionality, reversibility and constraints of the exchange reactions in the model, `ecmtool` will detect if the external metabolite can be used as an input, an output, or as both. This conclusion can be overruled easily by using the `--inputs`- and `--outputs`-arguments.

##### 3.2 Adding an objective metabolite

Many models contain an objective reaction, often a virtual reaction that uses all components that are needed for biomass production in the right proportions. Although ECMs in principle do not report any reaction rates, the rate of the objective reaction is most often of interest. Therefore, `ecmtool` by default adds an external 'objective metabolite' to the model that is produced in the objective reaction. In this manner, the rate of the objective reaction is reported in the production of the objective metabolite. The addition of this objective metabolite can be disabled by the `--add_objective_metabolite`-argument.

##### 3.3 Splitting metabolites

As was discussed in Section 2, not all ECMs are generating vectors of the steady-state conversion cone defined in (10), unless this cone is contained in one orthant. However, the conversion cone might not be contained in one orthant since some metabolites can both be consumed $\dot{c}_i < 0$ and produced $\dot{c}_i > 0$. In the Method-section of the main text, we explained how this can be remedied by splitting external metabolites into input- and output-metabolites. We will here make this explanation more precise.

We start with a stoichiometric matrix $N$, and possibly overlapping index sets *Input* and *Output*, indicating which metabolites can be taken up and which can be produced. We now create a new metabolite for each metabolite in *Input* and for each metabolite in *Output*. For convenience, let us give index $I_i$ to the virtual metabolite that is added if $i \in \mathit{Input}$, and the virtual metabolite added if $i \in \mathit{Output}$ index $O_i$. An extended stoichiometry matrix $\bar{N}$ is created by adding rows and columns for these metabolites. The new columns are of the shape $\hat{\boldsymbol{e}}_i - \hat{\boldsymbol{e}}_{I_i}$ and $-\hat{\boldsymbol{e}}_i + \hat{\boldsymbol{e}}_{O_i}$, where $\hat{\boldsymbol{e}}_i$ indicates the $i$-th elementary unit vector. These new columns model the irreversible 'reactions' $c_{I_i} \to c_i$ and $c_i \to c_{O_i}$. The metabolite $c_i$ is marked as internal, so that the steady-state assumption should be satisfied. We then enumerate the generator description of the conversion cone associated with this extended metabolic network. The results are mapped back to our original metabolic network by the unsplitting rule: $\dot{c}_i = \dot{c}_{I_i} + \dot{c}_{O_i}$.

**Theorem 3.** *Given a metabolic network, we follow the outlined recipe above to create an extended metabolic network. All Elementary Conversion Modes of the original metabolic network can be found as* *the generator representation of the conversion cone of the extended metabolic network.*

*Proof.* Let us first show that there is a bijection $f$ from the space of conversions of the original metabolic network, $\mathcal{C}$, to its image in the space of conversions of the extended metabolic network $\bar{\mathcal{C}}$. Let $\dot{\boldsymbol{c}}$ be a steady-state conversion of the original metabolic network. We can map the conversion to an extended conversion by defining

$$\begin{aligned} f_{I_i}(\dot{\boldsymbol{c}}) &= \dot{c}_i \text{ if } \dot{c}_i < 0 \text{ and } i \in \textit{Input}, \\ f_{O_i}(\dot{\boldsymbol{c}}) &= \dot{c}_i \text{ if } \dot{c}_i > 0 \text{ and } i \in \textit{Output}, \\ f_j(\dot{\boldsymbol{c}}) &= 0 \text{ otherwise.} \end{aligned}$$

This is a steady-state conversion for the extended metabolic network. Its inverse is then given by the unsplitting rule:

$$g_i(\bar{\boldsymbol{x}}) = \bar{x}_{I_i} + \bar{x}_{O_i}.$$

This map is a bijection between the original steady-state conversion cone and its image in the extended cone. This shows that any conversion in the original steady-state conversion cone is still present in the new conversion cone. Now, we only have to show that any ECM of the original metabolic network becomes a generator of the new conversion cone, and vice versa.

We thus first want to show that if $\dot{\boldsymbol{c}}$ is an ECM in the original metabolic network, then $f(\dot{\boldsymbol{c}})$ is a minimal generator of $\bar{\mathcal{C}}$. We will prove this by its contrapositive: assume that $f(\dot{\boldsymbol{c}}) \in \bar{\mathcal{C}}$ is *not* a minimal generator. Then, we know that there are two steady-state conversions $\bar{\boldsymbol{x}}, \bar{\boldsymbol{y}} \in \bar{\mathcal{C}}$ that are not multiples of each other, such that $f(\dot{\boldsymbol{c}}) = \lambda_1 \bar{\boldsymbol{x}} + \lambda_2 \bar{\boldsymbol{y}}$, with $\lambda_1, \lambda_2 > 0$. We can now apply the inverse mapping $g$ to find a similar decomposition in $\mathcal{C}$. According to the definition of ECMs, this shows that $\dot{\boldsymbol{c}}$ is not an ECM, unless for all $i$ we have $\lambda_1 g_i(\bar{\boldsymbol{x}}) = \lambda_2 g_i(\bar{\boldsymbol{y}})$, or if for some $i$ we have $\lambda_1 g_i(\bar{\boldsymbol{x}}) = -\lambda_2 g_i(\bar{\boldsymbol{y}})$. So, we have either $\lambda_1(\bar{x}_{I_i} + \bar{x}_{O_i}) = \lambda_2(\bar{y}_{I_i} + \bar{y}_{O_i})$, or $\lambda_1(\bar{x}_{I_i} + \bar{x}_{O_i}) = -\lambda_2(\bar{y}_{I_i} + \bar{y}_{O_i})$. In addition, we know for each $i$ by the definition of the mapping that either $f_{I_i}(\dot{\boldsymbol{c}}) = 0$, or $f_{O_i}(\dot{\boldsymbol{c}}) = 0$. Let us assume the former without loss of generality, then we know that $\bar{x}_{I_i} = \bar{y}_{I_i} = 0$. Adding the pieces together, we find that either $\lambda_1 \bar{x}_{O_i} = \lambda_2 \bar{y}_{O_i}$, or $\lambda_1 \bar{x}_{O_i} = -\lambda_2 \bar{y}_{O_i}$. Since $\bar{x}_{O_i}, \bar{y}_{O_i} \geq 0$, the latter is impossible, implying that for all $i$, $\lambda_1 \bar{x}_{O_i} = \lambda_2 \bar{y}_{O_i}$. But this is in direct contradiction with $\bar{\boldsymbol{x}}$ and $\bar{\boldsymbol{y}}$ not being multiples of each other, and $\dot{\boldsymbol{c}}$ can not be an ECM. We conclude that any ECM of the original metabolic network must map to a minimal generating vector of the extended conversion cone.

It remains to be shown that for all vectors $\bar{\boldsymbol{x}}$ in the minimal generating set of $\bar{\mathcal{C}}$, $g(\bar{\boldsymbol{x}})$ is an ECM. Let's again use contrapositivity, such we assume that $g(\bar{\boldsymbol{x}})$ can be written as $g(\bar{\boldsymbol{x}}) = \lambda_1 \dot{\boldsymbol{c}} + \lambda_2 \dot{\boldsymbol{d}}$ so that no metabolite is cancelled in the sum. Even stricter, if we choose appropriate convex combinations of

$\lambda_1\dot{\boldsymbol{c}}$ and $\lambda_2\dot{\boldsymbol{d}}$, we can find two vectors that we will call $\dot{\boldsymbol{c}}'$ and $\dot{\boldsymbol{d}}'$, for which we have $g(\bar{\boldsymbol{x}}) = \dot{\boldsymbol{c}}' + \dot{\boldsymbol{d}}'$ and that both lie in the same orthant as $g(\bar{\boldsymbol{x}})$. Since these two vectors are not multiples of each other and lie in the same orthant, the mapping $f$ will map them to two vectors that are also not multiples of each other, and for which we still have $\bar{\boldsymbol{x}} = f(\dot{\boldsymbol{c}}') + f(\dot{\boldsymbol{d}}')$. This shows that if $g(\bar{\boldsymbol{x}})$ is an ECM, then $\bar{\boldsymbol{x}}$ cannot be in the minimal generating set of $\bar{\mathcal{C}}$. So, by contrapositivity this means that all vectors in this minimal generating set map to ECMs of the original metabolic network. This completes the proof. □

As we will mention later, the splitting of metabolites can be postponed to later in the calculation if the 'indirect method' is used. This is sometimes beneficial, decreasing both memory usage and computation time, but not always. We have not found a clear indication of when it is useful to postpone the splitting. Therefore, we have added an argument `--splitting_before_polco` that determines when the splitting is done, so that the user can try out which method works best for the model of interest.

##### 3.4 Converting the stoichiometic coefficients into fractions

The ECM enumeration that we implemented works entirely with fractions. This is necessary to keep round-off errors from accumulating which would lead to reporting too many ECMs. Most metabolic networks have some reactions of which the stoichiometric coefficients are decimal numbers: usually the biomass reaction, and sometimes some 'maintenance' reaction. Each number with a finite number of decimals can be exactly converted to a fraction. This is what we do in this step.

It is important to note that coefficients with many decimal numbers will require the numerator and denominator of the fractions to be large numbers. These large integers can slow down the computations. The user might therefore choose to round off the decimal numbers in the input SBML-file if these do not matter too much.

##### 3.5 (Optional): Hiding metabolites and tagging reactions

We have discussed the `--hide`- and `--tag`-methods in Box 2 of the main text. Comma-separated lists can be given to `ecmtool` indicating of which metabolites the production and consumption should be ignored, and of which reactions the rates should be reported. When metabolites are hidden, virtual sink or source reactions are added at this point, and the metabolite is itself marked as internal. When reactions are tagged, virtual metabolites are added that are produced during these reactions.

### 4 A custom LP-solver

During the enumeration of ECMs we will often encounter Linear Programs of a specific type. We developed a customized LP-solver for this type of problems based on the conventional Revised Simplex Method, see for example [?]. Our solver outperformed other solvers on our type of Linear Programs by both speed and accuracy. In this section we describe the Revised Simplex Method and how we have optimized it.

#### 4.1 Problem description

In the ECM enumeration, we often encounter linear problems where an initial feasible solution is known, but where we want to know if there are alternative solutions. Specifically, let $\boldsymbol{\lambda}_{Init}$ be the initial solution, and let *Init* be the index set of its support. We want to know if there is a solution that uses more than only these reactions. We can use the following Linear Program

$$\begin{aligned} \underset{\boldsymbol{\lambda}}{\text{maximize}} \quad & \sum_{j \notin Init} \lambda_i \\ \text{subject to} \quad & R\boldsymbol{\lambda} = R\boldsymbol{\lambda}_{Init} \\ & \lambda_i \geq 0. \end{aligned} \tag{11}$$

A normal Linear Program would keep searching until the maximizer is found, but we are not interested in the maximizer. Rather, we are interested in *whether* an alternative solution exists. We thus developed an LP-solver with an 'early exit'. Moreover, we used that in this type of problems an initial solution is always known beforehand. Lastly, many of the problems that we will encounter are highly degenerate, as defined below. So, in summary, our problems have three specific features:

- Solver should stop when first alternative solution is found
- Initial feasible solution is known
- Problem is highly degenerate

Before, we can explain how our LP-solver exploits these features, we give a short introduction to Linear Programming. Then, we will describe the revised simplex method, and after that describe how we used and adapted this method.

#### 4.2 Some background on Linear Programming

Let us consider a Linear Program in the form of (11). Let $m \times n$ be the dimensions of $R$. It is safe to assume $n \geq m$; if this is not the case, there are more linear constraints than variables, and some of them will be redundant.

A *basis* for the LP is any set $B$ of $m$ indices from $\{1, \ldots, n\}$ such that the corresponding columns of $R$ form a basis of the column space of $R$. In other words, a basis is a $B$ such that the $m \times m$ submatrix $R_B$ of $R$ is non-singular.

We say that a feasible solution $\boldsymbol{\lambda}$ is a *basic feasible solution (BFS)* with basis $B$ if all the non-zero elements of $\boldsymbol{\lambda}$ are also elements of $B$ ($B$ may be a larger index set, though). Note that any basis $B$ has at most one corresponding BFS: since $R_B$ is nonsingular, $R_B\boldsymbol{\lambda} = \boldsymbol{x}$ has a unique solution $\boldsymbol{\lambda_B}$. This $\boldsymbol{\lambda_B}$ is a BFS if and only if it satisfies $\boldsymbol{\lambda_B} \geq \mathbf{0}$. The converse is not true: one BFS can have multiple bases. This occurs when there is a BFS $\boldsymbol{\lambda}$ that has fewer than $m$ non-zero elements. This is called a *degenerate* solution. We will often encounter degenerate solutions. In fact, the initial feasible solution that we will use is almost always degenerate.

Degenerate points can cause *cycling*, a situation where iterative solution methods such as the (revised) simplex method visit the exact same point more than once. When this happens, the method will again start the same cycle, and will thus never terminate. We will overcome this by perturbing the Linear Program, see Section 4.7.

#### 4.3 The revised simplex method

The revised simplex method uses the idea of a basis for a linear program to find an optimal solution. Essentially the method is based on the Karush-Kuhn-Tucker (KKT) conditions, which are necessary and sufficient conditions for the optimality of a Basic Feasible Solution. More information is available in the book *Numerical Optimization* [**?**], specifically chapter 12 and 13.

Suppose we have an LP problem in standard form:

$$\begin{aligned} \underset{\boldsymbol{x}}{\text{minimize}} \quad & \boldsymbol{c}^T\boldsymbol{x} \\ \text{subject to} \quad & A\boldsymbol{x} = \boldsymbol{b} \\ & x_i \geq 0, \end{aligned} \tag{12}$$

where we assume $A$ to have size $m \times n$ and rank $m$. The KKT optimality conditions are:

$$
\begin{aligned}
A\boldsymbol{x} &= \boldsymbol{b}, && (13)\\
A^T\boldsymbol{\pi} + \boldsymbol{s} &= \boldsymbol{c}, && (14)\\
\boldsymbol{x} &\geq \boldsymbol{0}, && (15)\\
\boldsymbol{s} &\geq \boldsymbol{0}, && (16)\\
\boldsymbol{s}^T\boldsymbol{x} &= 0. && (17)
\end{aligned}
$$

These conditions derive from constrained optimization using Lagrange multipliers, where $\boldsymbol{\pi}$ is the Lagrange multiplier associated with the constraint $A\boldsymbol{x} = \boldsymbol{b}$, and $\boldsymbol{s}$ is the Lagrange multiplier associated with $\boldsymbol{x} \geq \boldsymbol{0}$. Practically it means that if we find a combination of $\boldsymbol{x}, \boldsymbol{\pi}$ and $\boldsymbol{s}$ such that all conditions are met, we have found the optimal solution to the LP. These conditions do not only tell us when we are done, but also induce an optimization strategy, which we describe below.

The revised simplex method starts in a Basic Feasible Solution $\boldsymbol{x}$ with feasible basis $B$ of length $m$. If the problem is non-degenerate, all entries in $\boldsymbol{x}_B$ will be strictly greater than zero. One can thus construct the feasible basis directly from a non-degenerate solution. However, we will often have an initial feasible point with fewer than $m$ nonzero's. In that case, fewer than $m$ columns of $A$ are needed to satisfy the constraints. To still be able to start with the revised simplex method, we must then supply these columns with additional columns from $A$ such that we get an invertible $m \times m$-matrix $A_B$. The way to find these additional columns is described in Section 4.5. Let us for now proceed by assuming that we have a basis $B$ of size $m$ corresponding to the feasible solution $\boldsymbol{x}$.

We denote by $N$ the indices not in $B$, and split $\boldsymbol{x}$, $\boldsymbol{c}$ and $\boldsymbol{s}$ according to $B$:

$$
\boldsymbol{x} = \begin{bmatrix} \boldsymbol{x}_B \\ \boldsymbol{x}_N \end{bmatrix} = \begin{bmatrix} A_B^{-1}\boldsymbol{b} \\ \boldsymbol{0} \end{bmatrix}, \qquad \boldsymbol{c} = \begin{bmatrix} \boldsymbol{c}_B \\ \boldsymbol{c}_N \end{bmatrix}, \qquad \boldsymbol{s} = \begin{bmatrix} \boldsymbol{s}_B \\ \boldsymbol{s}_N \end{bmatrix}.
$$

Then, we can also split the second KKT constraint in two parts:

$$
A_B^T\boldsymbol{\pi} + \boldsymbol{s}_B = \boldsymbol{c}_B, \tag{18}
$$

$$
A_N^T\boldsymbol{\pi} + \boldsymbol{s}_N = \boldsymbol{c}_N. \tag{19}
$$

In order to satisfy the last KKT condition, let $\boldsymbol{s}_B = \boldsymbol{0}$. Since $A_B$ and $\boldsymbol{c}_B$ are known, we can deduce from

(18) that

$$\boldsymbol{\pi} = (A_B^T)^{-1}\boldsymbol{c}_B. \tag{20}$$

Next, from (19) we can compute $\boldsymbol{s}_N$ according to:

$$\boldsymbol{s}_N = \boldsymbol{c}_N - A_N^T\boldsymbol{\pi}. \tag{21}$$

Now $\boldsymbol{x}$ satisfies the KKT conditions if and only if $\boldsymbol{s}_N \geq \mathbf{0}$, so we can tell whether or not the vertex $\boldsymbol{x}$ is optimal. If $\boldsymbol{x}$ is indeed optimal, we are done and the LP is solved. Otherwise, we perform a *pivot operation.*

#### 4.4 Pivoting: replacing one of the columns in the feasible basis

One step in the revised simplex method involves replacing one column index in $B$ by one from $N$.

To select the entering index, consider $\boldsymbol{s}_N$. At least one element is negative, otherwise the previous $\boldsymbol{x}$ would have been optimal. We will see later that the entries in $\boldsymbol{s}_N$ capture how much the objective value would decrease when the corrsponding column is added to the basis. Now choose any $q$ with $s_q < 0$ as the entering index. The procedure for altering $B$ and changing $\boldsymbol{x}$ and $\boldsymbol{s}$ accordingly is as follows:

1. Increase $x_q$ from zero;
2. Keep all other components of $\boldsymbol{x}_N$ at zero;
3. Change the current basic vector $\boldsymbol{x}_B$ in such a way that $A\boldsymbol{x} = \boldsymbol{b}$ remains satisfied;
4. Keep increasing $x_q$ until one of the components of $\boldsymbol{x}_B$ (say $x_p$) reaches zero, or determine that no such component exists (then the LP is unbounded);
5. Remove $p$ from $B$ and replace it with $q$.

To perform step 3 we can use the following reasoning. Call the new vertex $\boldsymbol{x}^+$. Since both $A\boldsymbol{x} = \boldsymbol{b}$ and $A\boldsymbol{x}^+ = \boldsymbol{b}$, and since $\boldsymbol{x}_N = \mathbf{0}$ and $x_i^+ = 0$ for $i \in N \setminus \{q\}$, we have

$$A\boldsymbol{x}^+ = A_B\boldsymbol{x}_B^+ + \boldsymbol{A}_q x_q^+ = A_B\boldsymbol{x}_B = A\boldsymbol{x}. \tag{22}$$

Using that $A_B$ is non-singular, we can left-multiply with $A_B^{-1}$ to obtain

$$\boldsymbol{x}_B^+ = \boldsymbol{x}_B - A_B^{-1}A_q x_q^+ = \boldsymbol{x}_B - \boldsymbol{d}x_q^+, \tag{23}$$

showing that we should subtract from $\boldsymbol{x}_B$ the vector $\boldsymbol{d} = A_B^{-1}\boldsymbol{A}_q$. Step 4 tells us that we should subtract this vector until one of the entries in $\boldsymbol{x}_B^+$ becomes zero. This means that we must have

$$x_q^+ = \min\left\{\frac{(\boldsymbol{x}_B)_1}{d_1}, \ldots, \frac{(\boldsymbol{x}_B)_m}{d_m}\right\}, \tag{24}$$

and that the leaving column has index $p = \operatorname{argmin}\left\{\frac{(\boldsymbol{x}_B)_1}{d_1}, \ldots, \frac{(\boldsymbol{x}_B)_m}{d_m}\right\}$.

We can verify that this pivot operation always leads to a decrease in the objective $\boldsymbol{c}^T\boldsymbol{x}$.

**Theorem 4.** *The above described pivot operation will always lead to a decrease in the objective function. The change is always strictly less than zero if the original solution $\boldsymbol{x}$ is non-degenerate.*

*Proof.* We know that

$$\boldsymbol{x}_N^+ = (0, \ldots, 0, x_q^+, 0, \ldots, 0)^T, \tag{25}$$

so

$$\begin{aligned} \boldsymbol{c}^T\boldsymbol{x}^+ &= \boldsymbol{c}_B^T\boldsymbol{x}_B^+ + \boldsymbol{c}_N^T\boldsymbol{x}_N^+ \\ &= \boldsymbol{c}_B^T\boldsymbol{x}_B^+ + c_q x_q^+ \\ &= \boldsymbol{c}_B^T\boldsymbol{x}_B - \boldsymbol{c}_B^T A_B^{-1}\boldsymbol{A}_q x_q^+ + c_q x_q^+. \end{aligned} \tag{26}$$

From (20) we have $\boldsymbol{c}_B^T A_B^{-1} = \boldsymbol{\pi}^T$, and from (19) we get $\boldsymbol{\pi}^T\boldsymbol{A}_q = c_q - s_q$. Therefore,

$$\boldsymbol{c}_B^T A_B^{-1}\boldsymbol{A}_q x_q^+ = \boldsymbol{\pi}^T\boldsymbol{A}_q x_q^+ = (c_q - s_q)x_q^+, \tag{27}$$

so by substituting in (26) we obtain

$$\boldsymbol{c}^T\boldsymbol{x}^+ = \boldsymbol{c}_B^T\boldsymbol{x}_B - (c_q - s_q)x_q^+ + c_q x_q^+ = \boldsymbol{c}_B^T\boldsymbol{x}_B - s_q x_q^+. \tag{28}$$

Since $\boldsymbol{x}_{\boldsymbol{N}} = 0$, we have $\boldsymbol{c}^T\boldsymbol{x} = \boldsymbol{c}_B^T\boldsymbol{x}_B$ and therefore

$$\boldsymbol{c}^T\boldsymbol{x}^+ = \boldsymbol{c}^T\boldsymbol{x} - s_q x_q^+. \tag{29}$$

We chose $q$ such that $s_q < 0$, and since $x_q^+ \geq 0$, it follows that the step produces a decrease in the objective function $\boldsymbol{c}^T\boldsymbol{x}$. The change in objective value is strictly less than zero whenever $x_q^+$ is strictly larger than zero. Since $x_q^+ = \min\left\{\frac{(\boldsymbol{x}_B)_1}{d_1}, \ldots, \frac{(\boldsymbol{x}_B)_m}{d_m}\right\}$, we are sure to have a strict decrease whenever $\boldsymbol{x}_B > \boldsymbol{0}$, which is the definition of a non-degenerate solution. □

It is important to emphasize what happens when we have a degenerate solution. In that case, one of the entries in $\boldsymbol{x}_B$ is zero, and the pivot operation might thus lead to a step of size zero: column $p$

was used zero times in the original solution, and is now replaced by column $q$ which is also used zero times. In this case, and in this case only, the objective value does not strictly decrease during the pivot operation. This can be problematic, because the simplex method could later return to the same basis. If the same pivot step is again made at that point, one could end up in an infinite cycle. Such cycling is impossible in the non-degenerate case, since the strict decrease in objective value prevents the same basis from re-occurring. This problem can be prevented by keeping track of the bases that have already been visited, or by perturbing the system to get rid of degeneracy. We will describe in Section 4.7 that we choose the latter.

#### 4.5 Finding a starting basis efficiently

As mentioned in Section 4.1, we always already have an initial feasible solution for the Linear Programs that we must solve in this work. This initial $\boldsymbol{x}$ satisifies the constraints, but it is not yet a Basic Feasible Solution. For that we need an index set $B$ of length $m$ that contains at least all indices that correspond to the nonzero's of $\boldsymbol{x}$, and such that $A_B$ is non-singular. Because our initial feasible solutions are often degenerate, $\boldsymbol{x}$ generally comprises fewer than $m$ nonzero entries, so that we should supply $B$ with columns of $A$ that do not contribute to satisfying the constraints, i.e., the corresponding entry in $\boldsymbol{x}$ is zero.

To do this, we start by taking the indices in which $\boldsymbol{x}$ is non-zero. The columns corresponding to these indices should be linearly independent, otherwise we must find an alternative solution where one of the columns is no longer necessary. However, in the problems that we will encounter, the linear independence is guaranteed by the preceding steps. Then, we start iterating over the columns of $A$: we try to add a column and check to see if the resulting matrix is still of maximal rank. When this is not the case, the most recently added column is dropped again. After trying this for all columns of $A$, we are guaranteed to find $m$ total columns that are linearly independent, since $A$ has rank $m$.

Finding a basis in this manner is a time-consuming computation, but we do not have to go through this procedure each time that we do an LP. Indeed, as will become clear in Sections 7.3.3 and 8.1.2, we will need to do many LPs for the same constraint matrix $A$, but with a different initial feasible solution $\boldsymbol{x}$. This means that we can use the basis $B$ found in the previous LP as a starting point. We only have to make sure that the indices corresponding to the nonzero's in $\boldsymbol{x}$ are in $B$. Therefore, we use the following method.

Let $A_B$ be an invertible matrix, and $\boldsymbol{A}_q$ a column. We want to construct a new invertible matrix $\bar{A}_B$ by replacing one of the columns from $A_B$ by $\boldsymbol{A}_q$. We solve

$$A_B \boldsymbol{x} = \boldsymbol{A}_q, \tag{30}$$

which according to Cramer's rule gives

$$x_i = \frac{\det(A_B^{(i)})}{\det(A_B)}, \tag{31}$$

where $A_B^{(i)}$ is the matrix formed by replacing the $i$-th column of $A_B$ by $\boldsymbol{A}_q$. Since $\boldsymbol{A}_q$ is nonzero, there must be an $i$ for which $x_i \neq 0$, which implies that $\det(A_B^{(i)}) \neq 0$. We can thus find a new invertible matrix by replacing the $i$-th column of $A_B$ by $\boldsymbol{A}_q$.

#### 4.6 Detecting an early exit possibility

As mentioned in Section 4.1, the LP-solver may stop when it is clear that the initial solution is not the only possible solution. This means that we can stop the program at two events: 1) when the KKT conditions are met, meaning that the initial feasible solution is the only (and thus the optimal) solution, or 2) when the first non-zero step is made during a pivot operation, showing that an alternative solution exists.

From (24), we know that the step size is equal to $\min\left\{\frac{(\boldsymbol{x}_B)_1}{d_1}, \ldots, \frac{(\boldsymbol{x}_B)_m}{d_m}\right\}$, and that the index that leaves the basis $B$ is then equal to $\operatorname{argmin}\left\{\frac{(\boldsymbol{x}_B)_1}{d_1}, \ldots, \frac{(\boldsymbol{x}_B)_m}{d_m}\right\}$. Therefore, it is clear that a nonzero step is only made when the leaving index corresponds to a nonzero entry of $\boldsymbol{x}$. This means that we can stop the Linear Program when the leaving index corresponds to a nonzero entry of the initial solution $\boldsymbol{x}$.

#### 4.7 Perturbation to remove degeneracy

Degenerate vertices are problematic for the (revised) simplex method. When $\boldsymbol{x}$ is degenerate, the pivot operation described above might not cause any change in $\boldsymbol{x}$ at all. It is possible to make a number of degenerate pivots resulting in the same basis we had before. In this case the algorithm would start cycling and never terminate.

We use a perturbation strategy to circumvent this problem. We will call the original Linear Program $LP$ and the perturbed one $LP'$. The perturbation is applied after we have established a Basic Feasible Solution, with solution $\boldsymbol{x}$, and corresponding basis $B$. This solution of course satisfies the original constraint

$$A\boldsymbol{x} = \boldsymbol{b}. \tag{32}$$

However, we now perturb the right hand side of this equation to get

$$A\boldsymbol{x'} = \boldsymbol{b'} = \boldsymbol{b} + A_B\boldsymbol{\varepsilon}, \tag{33}$$

where $\boldsymbol{\varepsilon}$ is a vector of length $m$ with elements chosen uniformly at random from $[\delta/2, \delta]$, where $\delta > 0$ is a small constant. Note that $\boldsymbol{\varepsilon} > \mathbf{0}$. The solution corresponding to this new constraint is

$$\boldsymbol{x'} = A_B^{-1}\boldsymbol{b'} = A_B^{-1}(\boldsymbol{b} + A_B\boldsymbol{\varepsilon}) = \boldsymbol{x} + \boldsymbol{\varepsilon}. \tag{34}$$

Since we chose $\boldsymbol{\varepsilon} > \mathbf{0}$, this new solution still satisfies $\boldsymbol{x'} \geq \mathbf{0}$, and $B$ is still a feasible basis for the perturbed problem $LP'$.

**Theorem 5** (properties of perturbed LP)**.** *The following hold for $LP'$:*
*(a) $LP'$ is non-degenerate.*
*(b) If $B$ is a feasible basis of $LP'$, then $B$ is also a feasible basis of $LP$.*
*(c) If $B$ is an optimal basis of $LP'$, then $B$ is also an optimal basis of $LP$.*
*(d) If $x_q$ can leave and $x_p$ can enter in a pivot corresponding to $B$ in $LP'$, then the same holds in $LP$.*

See for example [**?**] for a proof. This theorem shows that we can use the perturbed version $LP'$ to do all the pivots, and once we find an optimal basis it is guaranteed that this basis is also optimal for the original problem. This shows that although the perturbation might affect the exact optimal value of the LP, it will never affect whether there is an alternative solution to the initial solution, or not. For our purposes, the perturbation of the LP thus has no disadvantages, while it does prevent the revised simplex method from entering an infinite loop.

### 5 Compression of the metabolic networks

To reduce the size of the main step in ECM-enumeration, the metabolic network can be compressed by various methods. The compression steps that we have used all leave the eventual set of ECMs unchanged. The amount by which the network can be compressed is one of the advantages of ECM-enumeration compared to EFM-enumeration: because the individual reaction rates are not reported, many reactions can be merged or even deleted. The first four compression steps are adapted from [**?**], the compression of cycles and the removal of redundant rays has been added by us.

These compression steps are not completely independent. If compression step A is executed, then another execution of A will not remove any more metabolites or reactions. However, after executing compression step B, A might again be able to compress the network further. Therefore, to maximise the compression of the network, the following sequence of compression steps is repeated until a complete sequence did not remove another metabolite or reaction.

#### 5.1 Removal of infeasible reactions

Some reactions can never be active in a steady state solution that satisfies the irreversibility constraints. Before we start the ECM enumeration, these reactions can be safely removed, since we are only looking for steady state conversions.

A first test if a reaction is feasible can be done by calculating the nullspace of the matrix $N_{int}$, denoting the part of the stoichiometric matrix corresponding to internal metabolites. Let $K$ be a matrix with as columns a basis for the nullspace, so that we have for each column $NK_{\bullet i} = 0$. Note that the nullspace contains the full steady state flux cone. An entry $k_{ji}$ of the matrix $K$ denotes the rate of reaction $j$ in the $i$-th vector of the nullspace. If $K$ contains a row of only zeros, this means that this reaction can never be active in a steady state solution. The reactions corresponding to zero rows of $K$ can thus be removed from the network.

The nullspace contains all solutions that satisfy the steady state constraint $N_{int}\boldsymbol{v} = 0$, but might also include solutions that do not satisfy the irreversibility constraints: $v_i \geq 0$ for all irreversible reactions $i$. Therefore, we can do another test, suggested by Urbanczik et al. [?] to check if reactions become infeasible if we take these irreversibility constraints into account.

All solutions that satisfy the steady state constraint can be written as $v = K\lambda$, because the columns of $K$ form a nullspace of $N$. Here, the rate of the $i$-th reaction in the solution is given by $v_i = K_{i\bullet}\lambda$. We can get a sum of reaction rates by left-multiplying $v = K\lambda$ by a vector: $\mu K\lambda = \sum_i \mu_i v_i$. To test for reaction feasibility, we select the submatrix $K_{irr}$ of $K$ given by all rows that correspond to irreversible reactions. Then, we check, with a Linear Program, if there exists a non-zero vector $\mu$ such that $\mu_i \geq 0$ for all $i$, and $\mu K_{irr} = 0$. If such a vector exists, this means that for any $\lambda$ we have $0 = \mu K_{irr}\lambda = \sum_{\text{irreversible}} \mu_i v_i$. In other words, for all steady state solutions, the sum of reaction rates with weights $\mu_i$ is zero. However, since all $\mu_i$ are positive, this implies that some of the $v_i$ must be negative, but this would violate the irreversibility constraints. Therefore, the only option is that all $v_i$ are zero, and thus infeasible. Concluding, all reactions $i$ such that $\mu_i$ is strictly greater than zero must be zero, are thus infeasible, and can be removed.

#### 5.2 Dead-end metabolites are deleted

Internal metabolites that can either only be produced or only be consumed, are sometimes called *dead-end metabolites*. The ECMs are calculated under the constraint that all internal metabolites are in steady-state. A dead-end metabolite can therefore not be produced/consumed at all, because there are no consuming/producing reactions to maintain the steady-state. Before ECM-computation we can therefore delete these metabolites and all (solely producing or consuming) reactions that connect to them.

##### 5.3 Cancelling metabolites with a reversible reaction

A reversible reaction can be used to delete an internal metabolite without changing the space of steady-state conversions, see Figure S4b. Let's say that we have a metabolic network with $r$ reactions and that the last one of those is reversible. We also assume that there is at least one internal metabolite involved in this reaction, say metabolite $i$. Now, we are going to cancel the production or consumption of metabolite $i$ from each reaction by adding or subtracting reversible reaction $r$. We denote by $\epsilon_j^r$ the number of times that we need to add reaction $r$ to reaction $j$ to cancel metabolite $i$. The resulting stoichiometric matrix, $\bar{N}$ thus has the same number of columns, but only column $r$ still has a non-zero $i$-th entry. To see that this change of stoichiometry does not change the steady-state conversion space, let's recall the definition:

$$\mathcal{C} = \{\dot{\boldsymbol{c}} = N\boldsymbol{v} \ | \ N_{\text{Int}}\boldsymbol{v} = \boldsymbol{0}, \ v_i \geq 0 \text{ if } i \text{ irreversible}\}. \tag{35}$$

Let $\dot{\boldsymbol{c}} \in \mathcal{C}$ be a conversion in the original metabolic network, and let $\boldsymbol{v}$ be a flux vector that leads to this conversion. Then we know that $\dot{\boldsymbol{c}} = N\boldsymbol{v} = \sum_j \boldsymbol{N}_{\bullet j} v_j$. We now prove that this conversion can still be generated with the new stoichiometric matrix $\bar{N}$. We have

$$\dot{\boldsymbol{c}} = \sum_j \boldsymbol{N}_{\bullet j} v_j = \sum_j (\bar{\boldsymbol{N}}_{\bullet j} - \epsilon_j^r \bar{\boldsymbol{N}}_{\bullet r}) v_j = \bar{N}\bar{\boldsymbol{v}}, \tag{36}$$

where $\bar{\boldsymbol{v}} = \boldsymbol{v} + \sum_j \epsilon_j^r \hat{\boldsymbol{e}}_r$. Since reaction $r$ was reversible, $v_r$ is allowed to be negative. Therefore, $\bar{\boldsymbol{v}}$ is certainly a feasible steady-state flux vector, and the original conversion $\dot{\boldsymbol{c}}$ is therefore still possible. This compression is not possible for irreversible reactions, because $\sum_j \epsilon_j^r$ might be negative.

The advantage of changing the stoichiometric matrix from $N$ to $\bar{N}$ is that we are now in the situation of Section 5.2: metabolite $i$ has become a dead-end metabolite. We can therefore now cancel both metabolite $i$ and reaction $r$. In Figure S4b we illustrate how reversible reaction $v_3$ can be used to cancel metabolite $x_2$.

##### 5.4 Cancelling singly produced or consumed metabolites

A metabolite that is either produced by only one reaction, or consumed by only one reaction, can be cancelled without changing the space of steady-state conversions, see Figure S4a. Let's say that reaction $r$ is the only reaction in the metabolic network that produces metabolite $i$. (The case in which metabolite $i$ is consumed by only one reaction is similar and will thus not be treated here.) We can now add reaction

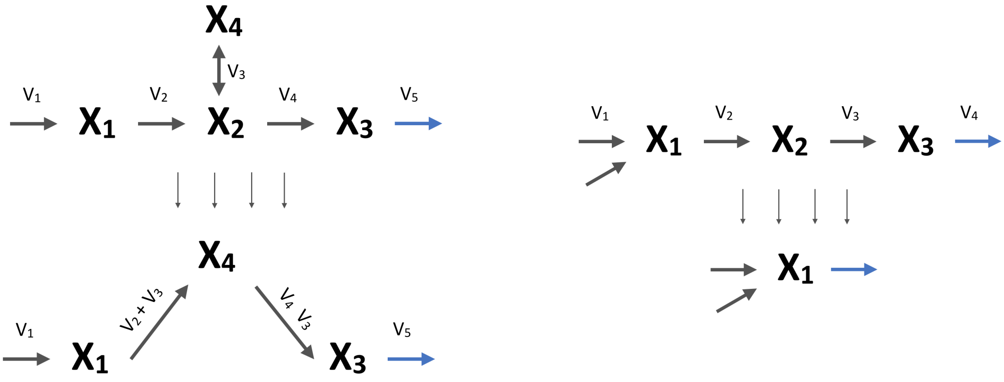

(a) (b)

Figure S4: **Conversion cones are invariant to network compression.** (a) Compression through addition of a reversible reaction to other reactions. $V_3$ is added to $V_2$ and subtracted from $V_4$. This sets the stoichiometry of $X_2$ to 0 in $V_2$ and $V_4$. Since $X_2$ is now only used in $V_3$, they can both be removed. (b) Compression through the removal of singly produced or consumed internal metabolites. As $X_2$ is only produced by $V_2$, $V_2$ can be added directly to $V_3$. Since $X_2$ is now only used in $V_2$, both $X_2$ and $V_2$ can be removed. The same can then be done for $X_3$.

$r$ to all reactions $j$ that consume metabolite $i$, such that the consumption of $i$ is cancelled exactly. We again denote by $\epsilon_j^r$ the number of times that we need to add reaction $r$ to reaction $j$ to cancel the consumption of metabolite $i$. The modified stoichiometric matrix is given by $\bar{N}$. Note that in this case, all $\epsilon_j^r$ are nonnegative. For a general conversion $\dot{\boldsymbol{c}}$ we again get

$$\dot{\boldsymbol{c}} = \sum_j \boldsymbol{N}_{\bullet j} v_j = \sum_j (\bar{\boldsymbol{N}}_{\bullet j} - \epsilon_j^r \bar{\boldsymbol{N}}_{\bullet r}) v_j = \bar{N} \bar{\boldsymbol{v}}, \tag{37}$$

where $\bar{\boldsymbol{v}} = \boldsymbol{v} + \sum_j \epsilon_j^r \hat{\boldsymbol{e}}_r$. Because all $\epsilon_j^r \geq 0$, this change will not violate the irreversibility constraints, and therefore all conversions remain feasible in the modified metabolic network.

Again we end up in the situation of Section 5.2: metabolite $i$ and reaction $r$ can be removed. In Figure S4a we illustrate how, for example, complete linear pathways can be reduced to just one reaction by this method.

#### 5.5 Removing cycles by cancelling metabolites

It is possible that the metabolic network contains *cycles*: combinations of reactions that have a combined production and consumption of zero. These combinations of reactions can also be viewed as vectors $\boldsymbol{v}$ that satisfy the irreversibility constraints and are in the nullspace of the stoichiometric matrix: $N\boldsymbol{v} = \boldsymbol{0}$. We will show that these cycles can in some sense be viewed as reversible reactions, and can therefore be used to cancel reactions and metabolites. Moreover, for the direct intersection method that we will describe below, it is necessary that the cone spanned by the columns of $N$ is pointed (see Section 1.2), which means that cycles may not exist. For the use of this intersection method, this compression step therefore must be applied.

We will detect cycles in $N$ by solving a Linear Program with our custom LP-solver described in Section 4. We will try to find a $\boldsymbol{\lambda} \geq \mathbf{0}$ that satisfies $N\boldsymbol{\lambda} = \mathbf{0}$ and $\boldsymbol{\lambda} \neq \mathbf{0}$; if such a $\boldsymbol{\lambda}$ exists, there are still cycles. The cycle finding LP is as follows:

$$\begin{aligned} \underset{\boldsymbol{\lambda}}{\text{maximize}} \quad & \sum_i \lambda_i \\ \text{subject to} \quad & N\boldsymbol{\lambda} = \mathbf{0} \\ & \lambda_i \geq 0 \\ & \lambda_i \leq 1. \end{aligned} \tag{38}$$

The LP is always feasible, since $\boldsymbol{\lambda} = \mathbf{0}$ is a solution with objective value 0. If this is the only feasible solution, then we are sure that cycles no longer exist in the metabolic network, so that this compression step is done. If another feasible solution exists, it will always be found by the Linear Program because it must result in a larger objective value.

Let's say that we have a cycle, then the optimal solution $\boldsymbol{\lambda}^*$ may be assumed to have at least one position equal to 1, say $\lambda_r^* = 1$. This is because any $\boldsymbol{\lambda}$ that induces a cycle satisfies $N\boldsymbol{\lambda} = \mathbf{0}$, so multiples of $\boldsymbol{\lambda}$ will satisfy that as well. Hence the constraint $\lambda_i \leq 1$ (for all $i$) is the only thing keeping $\lambda$ (and with that the optimal value) bounded. The corresponding column $\boldsymbol{N}_{\bullet r}$ gives the stoichiometry of a reaction that is involved in the cycle. This will be used to remove (part of) the cycle.

Let's say that metabolite $i$ is produced by reaction $r$. Because we have split each external metabolite into being only an input or only an output in Section 3.3, it is impossible for external metabolites to have non-zero coefficients in any ray that is part of a cycle, as there can be no circular flow through such external metabolites. We can thus be assured that metabolite $i$ is an internal metabolite.

Because reaction $r$ is part of a cycle, we have

$$\mathbf{0} = \sum_j \boldsymbol{N}_{\bullet j} v_j = \sum_{j \neq r} \boldsymbol{N}_{\bullet j} v_j + \boldsymbol{N}_{\bullet r} v_r = N\boldsymbol{v}^- + \boldsymbol{N}_{\bullet r} v_r, \tag{39}$$

where $\boldsymbol{v}^-$ are all reaction rates in the cycle except for reaction $r$: $\boldsymbol{v}^- = \boldsymbol{v} - v_r \hat{\boldsymbol{e}_r}$. Comparing this with the case in which we had a reversible reaction, reaction $r$ can now be viewed as the forward reaction, and the combination of the other reactions: $N\boldsymbol{v}^-$ as the backward reaction. Let us in fact add this combination of reactions as reaction $r+1$ to our network. We can now use these two 'reactions' to cancel the production and consumption of metabolite $i$ from all reactions in the network. If a reaction consumes metabolite $i$, then we will cancel this consumption by adding reaction $r$; let $\epsilon_j^r$ denote the number of times that reaction $r$ is added to reaction $j$. If reaction $j$ produces metabolite $i$, then we can cancel the production by adding reaction $r+1$; $\epsilon_j^{r+1}$ denote how many times this is necessary. We then get for any conversion that was feasible in the original metabolic network that

$$\dot{\boldsymbol{c}} = \sum_j \boldsymbol{N}_{\bullet j} v_j = \sum_j (\bar{\boldsymbol{N}}_{\bullet j} - \epsilon_j^r \bar{\boldsymbol{N}}_{\bullet r} - \epsilon_j^{r+1} \bar{\boldsymbol{N}}_{\bullet r+1}) v_j = \bar{N} \bar{\boldsymbol{v}}, \tag{40}$$

where $\bar{\boldsymbol{v}} = \boldsymbol{v} + \sum_j \epsilon_j^r \hat{\boldsymbol{e}}_r + \sum_j \epsilon_j^{r+1} \hat{\boldsymbol{e}}_{r+1}$. Since all $\epsilon$-values are now nonnegative, we now that this flux vector meets the irreversibility constraints, and thus that the conversion is still possible in the modified metabolic network.

Finally, we are left with a metabolic network in which metabolite $i$ is produced by reaction $r$ and consumed by reaction $r+1$, and these reactions are their exact reverse. It is clear that the steady-state constraint imposes $v_r = v_{r+1}$ which means that the net contribution of these reactions is always zero. Therefore, we can delete both reaction $r$, reaction $r+1$, and metabolite $i$ from the network.

After this procedure we have deleted (at least) one metabolite and one reaction from the orginal metabolic network. It is often the case that cycles still remain, so that the cycle removal procedure has to be repeated until no cycles are left.

##### 5.5.1 Example

In the metabolic network of Figure S5, there is one cycle: (B $\rightarrow$ C $\rightarrow$ F $\rightarrow$ E $\rightarrow$ B). The cycle finding LP described above would find the solution $\boldsymbol{\lambda^*} = (0, 0, 1, 1, 1, 1, 0, 0, 0, 0)$. So we learn that reactions 3, 4, 5 and 6 are involved in a cycle. We select reaction 3 and metabolite $B$ for the removal of the cycle. The only other reaction that uses $B$ is reaction 6. Therefore we add reaction 3 to reaction 6 to create a new ray that produces $C$ out of $E$. Then, reaction 3 can be removed. The resulting network can be seen in Figure S6. After three more steps, the network will be cycle-free (Figure S7).

#### 5.6 Removal of redundant reactions

We call reaction $r$ *redundant* if there is a feasible combination $\boldsymbol{w}$ of different reactions such that

$$\boldsymbol{N}_{\bullet r} = \sum_{j \neq r} \boldsymbol{N}_{\bullet j} w_j. \tag{41}$$

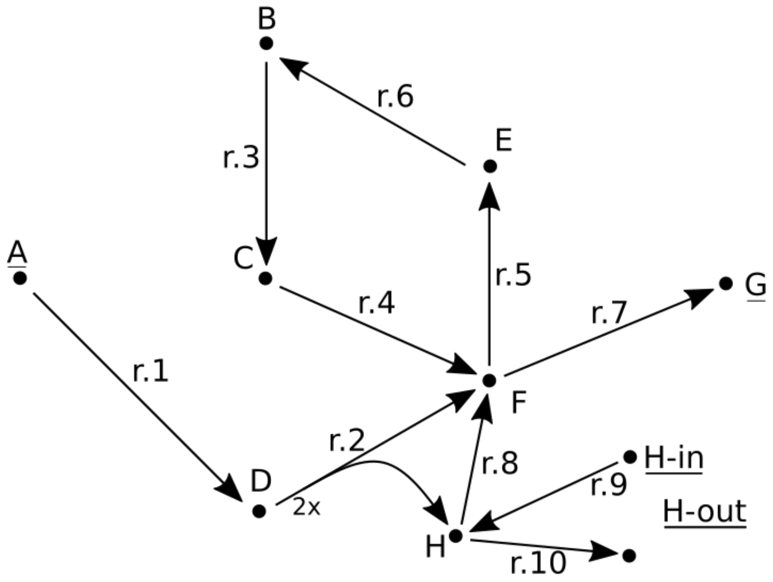

Figure S5: Example network for cycle removal.

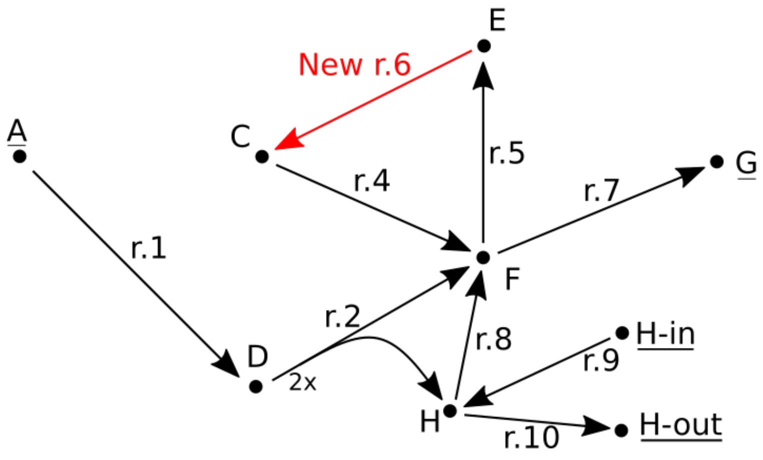

Figure S6: Network after one cycle removing step.

These redundant reactions can be removed without changing the steady-state conversion space. To see this, let $\dot{\boldsymbol{c}} \in \mathcal{C}$ be any conversion, then we have

$$\dot{\boldsymbol{c}} = N\boldsymbol{v} = \sum_j \boldsymbol{N}_{\bullet j} v_j = \sum_{j \neq r} \boldsymbol{N}_{\bullet j} v_j + \boldsymbol{N}_{\bullet r} v_r = \sum_{j \neq r} \boldsymbol{N}_{\bullet j} v_j + \sum_{j \neq r} \boldsymbol{N}_{\bullet j} w_j v_r = \sum_{j \neq r} \boldsymbol{N}_{\bullet j} (v_j + w_j v_r). \quad (42)$$

So, all steady state conversions remain feasible if we remove the redundant reaction from the network. The redundant reactions can be removed using `redund` from `lrslib` [?]. Although this program works well on relatively small sets of vectors (up to hundreds), it is very slow on larger sets. We have therefore

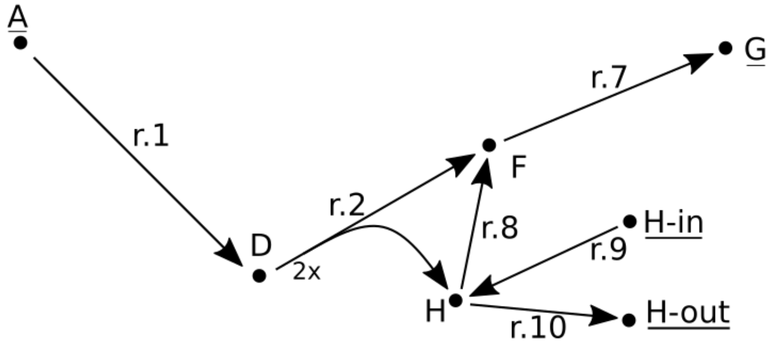

Figure S7: Network after all cycles have been removed.

developed our own redundancy-removal method that uses the following Linear Program:

$$\begin{aligned} \underset{\boldsymbol{v}}{\text{maximize}} \quad & \sum_{j \neq r} v_i \\ \text{subject to} \quad & N\boldsymbol{v} = \boldsymbol{N}_{\bullet r} \\ & v_i \geq 0. \end{aligned} \tag{43}$$

In this LP, we are not necessarily interested in the maximum, but rather in the question whether there is a solution besides $\boldsymbol{v} = \hat{\boldsymbol{e}}_{\boldsymbol{r}}$. This means that we can quit the program once it is clear that such an alternative solution exists. This is exactly the type of problem for which we designed the LP-solver described in Section 4. Note that this LP becomes unbounded whenever the cone generated by the columns of $N$ is non-pointed. In this case, our strategy would not work. Therefore, it is important to remove all cycles before removing redundant reactions (Section 5.5).

### 6 The starting point: generator representation of the (non-steady-state) conversion cone

After the metabolic network has been preprocessed (Section 3) and compressed (Section 5), we are now ready for the main computational task. Let us therefore recall the goal of ECM enumeration. We want to describe the *steady-state conversion cone*

$$\mathcal{C} = \{\dot{\boldsymbol{c}} = N\boldsymbol{v} \mid N_{\text{Int}}\boldsymbol{v} = \boldsymbol{0},\ v_i \geq 0 \text{ if reaction } i \text{ irreversible}\}. \tag{44}$$

Assuming that this cone is fully contained in one orthant, which can be assured by splitting metabolites (Section 3.3), the Elementary Conversion Modes can be found as a minimal set of generators of this cone.

Both the indirect method (Section 7) and the direct method (Section 8) start from a generator representation of a larger cone: the *(non-steady-state) conversion cone*,

$$\mathcal{C}_0 = \{\dot{\boldsymbol{c}} = N\boldsymbol{v} \mid v_i \geq 0 \text{ if reaction } i \text{ irreversible}\}, \tag{45}$$

which comprises all conversions, including those that do not satisfy the steady-state constraint. We can assume without loss of generality that all reactions in the metabolic network are irreversible, since they were either cancelled during the compression step, or they were split into a forward and backward reaction. In that case, (45) already gives a generator representation: the columns of the stoichiometry matrix, $N$, generate all possible conversions. The remaining task is to impose the steady-state constraints while keeping track of a generator representation. We will describe the two methods that we implemented to accomplish this.

The cone $\mathcal{C}_0$ is generally not contained in one orthant, nor is it necessarily pointed. In Section 3.3 we describe why it is beneficial to split external metabolites such that $\mathcal{C}$ is contained in one orthant, but this is not yet necessary for this initial cone $\mathcal{C}_0$. We will describe below that in the indirect method, external metabolites do not even have to be split already because it can be done later. The same holds for the pointedness: we cannot use the direct method on a non-pointed $\mathcal{C}_0$, while it will give no problems in the indirect method. Therefore, `ecmtool` always removes cycles before using the direct method, thereby making $\mathcal{C}_0$ pointed (Section 5.5).

### 7 The indirect method

The indirect method of ECM enumeration is based on the method introduced by Urbanczik et al. [?]. It is based on the fact that it is relatively easy to calculate a generator representation of a polyhedral cone when an inequality representation is known. This task can be accomplished by the Double Description (DD) method [?], and we use `Polco` for this [?]. Crucially, the indirect method depends on the connection between the generator representation of a cone and the inequality representation of its dual, see Section 1.3 for background information about this connection. More precisely, we will follow the following steps to go from the generator representation (45) to the set of ECMs:

$$\text{gen}(\mathcal{C}_0) = \text{ineq}(\mathcal{C}_0^*) \overset{\text{DD}}{\rightarrow} \text{gen}(\mathcal{C}_0^*) = \text{ineq}(\mathcal{C}_0) \overset{\text{SS constraints}}{\longrightarrow} \text{ineq}(\mathcal{C}) \overset{\text{DD}}{\rightarrow} \text{gen}(\mathcal{C}) = \text{ECMs} \tag{46}$$

One might at this point ask why the steady-state constraints cannot be added at the start, following the path $\text{gen}(\mathcal{C}_0) \rightarrow \text{gen}(\mathcal{C})$ at once. The answer is that this is indeed possible, and this is the strategy implemented by the direct method (Section 8). However, it is generally harder to impose equality constraints to a generator representation than to an inequality representation, where they can just be added as two inequality constraints to the existing representation. This is why this indirect route is in many cases, but not always, preferable, and this intersection method is therefore the default in `ecmtool`, which can be changed by setting `--direct True`.

In the following subsections we will give the details of the method, and where we have added optimization steps.

#### 7.1 $\mathbf{gen}(\mathcal{C}_0) = \mathbf{ineq}(\mathcal{C}_0^*)$

The columns of the stoichiometry matrix form a generator representation of the non-steady-state conversion cone (see (45)). In Section 1.3 we then explained that this gives an inequality representation to the dual cone: the constraint matrix being the transpose of the stoichiometry matrix. However, this dual cone does not need to be pointed, see Section 1.2. The problem with non-pointed cones is that generally they do not have a unique generator representation. The next step in the method, $\text{ineq}(\mathcal{C}_0^*) \overset{\text{DD}}{\to} \text{gen}(\mathcal{C}_0^*)$, would therefore not be well-defined. To still be able to apply the Double Description method, we decompose our cone in its lineality space and its pointed part, according to

$$\begin{aligned} \mathcal{C}_0^* &= \text{Lin}(\mathcal{C}_0^*) \oplus (\mathcal{C}_0^* \cap \text{Lin}(\mathcal{C}_0^*)^{\perp}), \qquad (47) \\ &= \left\{ \boldsymbol{x} \in \mathbb{R}^d \;\middle|\; N^T \boldsymbol{x} = \boldsymbol{0} \right\} \oplus \left\{ \boldsymbol{x} \in \mathbb{R}^d \;\middle|\; N^T \boldsymbol{x} \geq \boldsymbol{0}, \quad \text{Null}(N^T)^T \boldsymbol{x} = \boldsymbol{0} \right\}, \end{aligned}$$

where $\text{Null}(N^T) = \begin{bmatrix} \boldsymbol{n}_1 & \cdots & \boldsymbol{n}_k \end{bmatrix}$ is a matrix constructed from a basis of the nullspace of $N^T$. This shows that it is necessary to find a basis for the nullspace of $N^T$. First of all, because $\{\boldsymbol{n}_1, \ldots, \boldsymbol{n}_k, -\boldsymbol{n}_1, \ldots, -\boldsymbol{n}_k\}$ gives a generator representation of the lineality space (first part in (47)). Second, because we need this basis to complete the inequality representation of the pointed part of $\mathcal{C}_0^*$ (second part in (47)).

##### 7.1.1 An iterative symbolic nullspace calculation

It turns out that obtaining an exact basis for a nullspace for a large matrix with potentially large fractions is not a trivial computational task. We cannot resort to faster methods using floats, because this will induce round-off errors that might further accumulate during the ECM-enumeration. We have therefore implemented an iterative nullspace calculation which avoids memory issues. For this, we separate rows of $N^T$ into $K$ parts, denoting the first set of rows by $N_{(1)}$. We calculate a basis for its nullspace using a symbolic solver, and gather the basis vectors as the columns of a matrix $M$. We then know that any vector $\boldsymbol{x}$ in the nullspace of $N^T$ should be a linear combination of the columns in $M$, in other words: $\boldsymbol{x} = M\boldsymbol{\lambda}$, for some real vector $\boldsymbol{\lambda}$. We now take the second part of our matrix, $N_{(2)}$. We should have

$N_{(2)}\boldsymbol{x} = \mathbf{0}$, so that $\boldsymbol{\lambda}$ should satisfy $N_{(2)}M\boldsymbol{\lambda} = \mathbf{0}$. Therefore, we define a matrix $N_{(1:2)} := N_{(2)}M$, and calculate its nullspace. Again we gather a basis for this nullspace as the columns of $M$. Proceeding in this fashion, we will eventually obtain a matrix $M$ containing a basis for the nullspace of $N_{(1:K)} = N^T$.

#### 7.2 $\mathbf{ineq}(\mathcal{C}_0^*) \overset{\mathbf{DD}}{\rightarrow} \mathbf{gen}(\mathcal{C}_0^*)$

The dual of the non-steady-state conversion cone, $\mathcal{C}_0^*$ consists of the two parts given in (47). Fortunately, a set of generators of the entire space is just given by the union of the generators of both parts:

$$\begin{aligned}\text{gen}\left(\mathcal{C}_0^*\right) &= \text{gen}\left(\text{Lin}(\mathcal{C}_0^*)\right) \cup \text{gen}\left(\left(\mathcal{C}_0^* \cap \text{Lin}(\mathcal{C}_0^*)^{\perp}\right)\right), \\ &= \{\boldsymbol{n}_1, \ldots \boldsymbol{n}_k, -\boldsymbol{n}_1, \ldots \boldsymbol{n}_k\} \cup \text{DD}\left(\left\{\boldsymbol{x} \in \mathbb{R}^d \;\middle|\; N^T\boldsymbol{x} \geq \mathbf{0}, \quad \text{Null}(N^T)^T\boldsymbol{x} = \mathbf{0}\right\}\right)\end{aligned}$$

Note that the first part of this generator representation is not necessarily unique, because there is no unique basis for the nullspace. However, since this generator representation is only an intermediate result, this does not matter. What matters is that the cone described by this generator representation is unique.

The Double Description method that we use to get the second part of the generator representation is described in Section 1.5 and implemented in `Polco`[?]. This step, like all others, is done using fractions, so that our ECM-computations remain exact.

#### 7.3 $\mathbf{gen}(\mathcal{C}_0^*) = \mathbf{ineq}(\mathcal{C}_0) \overset{\textbf{SS constraints}}{\longrightarrow} \mathbf{ineq}(\mathcal{C})$

In the previous sections we have described how we obtain a generator representation of the dual of the (non-steady-state) conversion cone $\mathcal{C}_0^*$. This generator representation also forms an inequality representation of $\mathcal{C}_0$. Let us gather all these inequalities in a matrix $H$, then we have

$$\mathcal{C}_0 = \{\dot{\boldsymbol{c}} \mid H\dot{\boldsymbol{c}} \geq \mathbf{0}\}\,. \tag{48}$$

##### 7.3.1 Dropping columns corresponding to internal metabolites

We see that the columns of $H$ thus correspond to the metabolites in the metabolic network. We specify that we have $|\text{Ext}|$ external metabolites, $|\text{Int}|$ internal metabolites, and let us assume that $H$ is ordered such that all columns corresponding to external metabolites come first, i.e., $H = \begin{bmatrix} H_{\text{Ext}} & H_{\text{Int}} \end{bmatrix}$. To obtain an inequality description of the steady-state conversion cone, we should impose the steady-state constraints $\dot{c}_i = 0$ for $i \in Int$. This can be done easily by adding two inequality constraints per internal metabolites:

$$\mathcal{C} = \left\{ \dot{\boldsymbol{c}} \in \mathbb{R}^{|\text{Ext}|+|\text{Int}|} \mid H^{+}\dot{\boldsymbol{c}} \geq \mathbf{0} \right\}, \quad \text{where } H^{+} = \begin{bmatrix} H_{\text{Ext}} & H_{\text{Int}} \\ 0 & I_{|\text{Int}|} \\ 0 & -I_{|\text{Int}|} \end{bmatrix}, \tag{49}$$

with $I_{|\text{Int}|}$ being the ($|\text{Int}| \times |\text{Int}|$)-identity matrix. Although this is a valid description of $\mathcal{C}$, it is not a very efficient one. We prove in the following Theorem that we can just drop all columns corresponding to the internal metabolites.

**Theorem 6.** *Let $\mathcal{C} = \{\boldsymbol{x} \in \mathbb{R}^{|Ext|+|Int|} \mid H^{+}\boldsymbol{x} \geq \mathbf{0}\}$, and $\mathcal{C}' = \{\tilde{\boldsymbol{x}} \in \mathbb{R}^{|Ext|} \mid H_{Ext}\tilde{\boldsymbol{x}} \geq \mathbf{0}\}$. There is a bijection between cones $\mathcal{C}$ and $\mathcal{C}'$.*

*Proof.* Define a mapping between the two cones by keeping only the first $|\text{Ext}|$ coordinates:

$$\begin{aligned} f &: \mathcal{C} \to \mathcal{C}', \\ &: \boldsymbol{x} \mapsto \begin{bmatrix} x_1 & \cdots & x_{|\text{Ext}|} \end{bmatrix}^T. \end{aligned}$$

We first show that $f$ indeed maps into $\mathcal{C}'$. Take an arbitrary element $\boldsymbol{x} \in \mathcal{C}$. We know that $H^{+}\boldsymbol{x} \geq \mathbf{0}$. The last rows imply that $0\boldsymbol{x}_{\text{Ext}} + I_{|\text{Int}|}\boldsymbol{x}_{\text{Int}} \geq \mathbf{0}$, and $0\boldsymbol{x}_{\text{Ext}} - I_{|\text{Int}|}\boldsymbol{x}_{\text{Int}} \geq 0$, so that we must have $\boldsymbol{x}_{\text{Int}} = \mathbf{0}$. The first rows of the inequalities then imply $\mathbf{0} \leq H_{\text{Ext}}\boldsymbol{x}_{\text{Ext}} - H_{\text{Int}}\mathbf{0} = H_{\text{Ext}}\boldsymbol{x}_{\text{Ext}}$, which means that $f(\boldsymbol{x}) = \boldsymbol{x}_{\text{Ext}} \in \mathcal{C}'$.

With the knowledge that for any $\boldsymbol{x} \in \mathcal{C}$, the components corresponding to internal metabolites have to be zero, we can define an inverse mapping

$$\begin{aligned} g &: \mathcal{C}' \to \mathcal{C}, \\ &: \tilde{\boldsymbol{x}} \mapsto \begin{bmatrix} \tilde{\boldsymbol{x}}^T & \mathbf{0}^T \end{bmatrix}^T. \end{aligned}$$

This mapping is clearly a left- and right-inverse of $f$, and therefore $f$ must be a bijection. □

We can thus proceed with $H_{\text{Ext}}$ as the inequality description of the steady-state conversion cone.

##### 7.3.2 Splitting external metabolites

In Section 3.3, we explained why we need to split external metabolites into input- and output-metabolites in order to calculate all ECMs instead of only the extreme conversions. We also indicated that this splitting can be done during the preprocessing step. However, one can also choose (by using the argument `--splitting_before_polco False`) to calculate $H_{\text{Ext}}$ without splitting the metabolites, and split the metabolites afterwards. This choice does affect the size of $H_{\text{Ext}}$ and therefore the computational complexity of ECM-enumeration, but we could get no clear idea of when which method is favourable. We recommend the user to try both methods when working with large models, because we have observed that this option sometimes makes a large difference.

Let us now assume that the external metabolites are not yet split into input- and output-metabolites, and say, for simplicity, that all metabolites can both be consumed and produced, so that all need to be split. Then we can use $H_{\text{Ext}}$ to define

$$H_{\text{split}} = \begin{bmatrix} H_{\text{Ext}} & H_{\text{Ext}} \\ I & 0 \\ 0 & -I \end{bmatrix}. \tag{50}$$

Now let us compare the cones that are described by these inequality representations:

$$\mathcal{C}_{\text{Ext}} = \{\boldsymbol{x} \in \mathbb{R}^{|\text{Ext}|} \mid H_{\text{Ext}}\boldsymbol{x} \geq \mathbf{0}\}, \tag{51}$$

$$\mathcal{C}_{\text{split}} = \{\tilde{\boldsymbol{x}} \in \mathbb{R}^{2|\text{Ext}|} \mid H_{\text{split}}\tilde{\boldsymbol{x}} \geq \mathbf{0}\}. \tag{52}$$

We can define a mapping $f$ from $\mathcal{C}_{\text{Ext}}$ to $\mathcal{C}_{\text{split}}$ by

$$\begin{aligned} f \;&:\; \mathcal{C}_{\text{Ext}} \to \mathcal{C}_{\text{split}}, \\ &:\; \boldsymbol{x} \mapsto \tilde{\boldsymbol{x}} = \left[\max(x_1, 0), \cdots, \max(x_{|\text{Ext}|}, 0), \min(x_1, 0), \cdots, \min(x_{|\text{Ext}|}, 0)\right]^T, \end{aligned}$$

We can show that this mapping is well-defined. Let $\boldsymbol{x} \in \mathcal{C}_{\text{Ext}}$, we have to check if $\tilde{\boldsymbol{x}}$ satisfies $H_{\text{split}}\tilde{\boldsymbol{x}} \geq 0$. First of all, it is clear from the definition of $f$ that $\tilde{\boldsymbol{x}}$ satisfies the sign constraints that form the two lower rows of $H_{\text{split}}$. Further, the first rows of $H_{\text{split}}$ give

$$\begin{aligned} H_{\text{split}}^{\text{top}}\tilde{\boldsymbol{x}} &= H_{\text{Ext}}\left[\max(x_1, 0), \cdots, \max(x_{|\text{Ext}|}, 0)\right)^T + H_{\text{Ext}}\left(\min(x_1, 0), \cdots, \min(x_{|\text{Ext}|}, 0)\right]^T \\ &= H_{\text{Ext}}\left[\max(x_1, 0) + \min(x_1, 0), \cdots, \max(x_{|\text{Ext}|}, 0) + \min(x_{|\text{Ext}|}, 0)\right]^T, \\ &= H_{\text{Ext}}\boldsymbol{x} \geq \mathbf{0}. \end{aligned}$$

This shows that any steady-state conversion in $\mathcal{C}_{\text{Ext}}$ is still a steady-state conversion in $\mathcal{C}_{\text{split}}$. The following Theorem shows that the ECMs of $\mathcal{C}_{\text{Ext}}$ can be calculated by computing the minimal generator set of $\mathcal{C}_{\text{split}}$.

**Theorem 7.** *There is a bijection between the set of Elementary Conversion Modes of $\mathcal{C}_{Ext}$ and the minimal generator set of $\mathcal{C}_{split}$.*

The proof of this theorem is very similar to the proof of Theorem 3 in Section 3.3. Therefore, we will not repeat it here.

If we compute a minimal generator in $\mathcal{C}_{\text{split}}$, we can thus map this back to an ECM in $\mathcal{C}_{\text{Ext}}$. For this, we use the left-inverse of $f$:

$$\begin{aligned} g &: C_{\text{split}} \to C_{\text{Ext}}, \\ &: \tilde{\boldsymbol{x}} \mapsto \boldsymbol{x} = \left[\tilde{x}_1 + \tilde{x}_{|\text{Ext}|+1}, \ldots, \tilde{x}_{|\text{Ext}|} + \tilde{x}_{2|\text{Ext}|}\right]^T. \end{aligned}$$

##### 7.3.3 Removing redundant inequalities

The inequality description of $\mathcal{C}_0$, gathered in the matrix $H$, does not contain redundant inequalities, because the `polco`-software returns a minimal set. However, we have used $H$ to define $H_{\text{Ext}}$ and then $H_{\text{split}}$. Often, and mostly when there are relatively many internal metabolite columns that are dropped, these operations cause inequalities to become redundant. In theory, this causes no problem for the next steps in the ECM-enumeration. However, when $H_{\text{split}}$ has many rows, the next Double Description step is much slower. We can therefore speed up the ECM-computation by removing redundant rows. For this, we can use either the `redund`-program from `lrslib` [?] on small sets of rows, or our own algorithm for redundancy removal. Our algorithm has the advantages of being parallelizable if `ecmtool` is used with `mpiexec` (see the user guide in Section 11), and of reporting a counter which indicates the progress of the redundancy-removal. By default, we thus use our algorithm. We will shortly discuss how this works.

We call a row redundant if it can be written as a conical combination of other rows, i.e.,

$$\boldsymbol{A}_{i\bullet} = \sum_j \lambda_j \boldsymbol{A}_{j\bullet}, \quad \text{where } \lambda_j \geq 0. \tag{53}$$

This row is called redundant for the following reason. If for all $j$ such that $\lambda_j > 0$ we have $\boldsymbol{A}_{j\bullet}\boldsymbol{x} \geq \boldsymbol{0}$, then automatically $\boldsymbol{A}_{i\bullet}\boldsymbol{x} = \sum_j \lambda_j \boldsymbol{A}_{j\bullet}\boldsymbol{x} \geq 0$. So, the inequality implied by $\boldsymbol{A}_{i\bullet}$ does not further constrain the cone. We can thus remove all rows for which we can find a conical combination as in Equation (53)[1]. We detect these redundant rows by solving a Linear Program for each row:

$$\begin{aligned} \underset{\boldsymbol{\lambda}}{\text{maximize}} \quad & \sum_{i \neq j} \lambda_i \\ \text{subject to} \quad & A^T \boldsymbol{\lambda} = \boldsymbol{A}_{i\bullet}^T \\ & \lambda_i \geq 0. \end{aligned} \tag{54}$$

[1] It is important that we first make sure that any duplicate rows are removed, since otherwise the procedure could remove both of them.

This problem is exactly of the form described in Section 4, so we can use our custom LP-solver to solve it. In Section 4.5 we described that for this LP-solver to work, we need to select a maximal set of linearly independent columns of $A^T$ as a starting basis. This basis should contain the column $A_i^T$ that we want to test for redundancy. Finding such a starting basis is a relatively complex computational task. However, we use the same matrix $A^T$ for each redundancy test, and we can therefore use almost the same starting basis. We should only make sure that we replace one of the columns in this basis by $\boldsymbol{A}_{i\bullet}^T$. In Section 4.5 we describe how we accomplish this.

It is important to note that this redundancy removal only works when the cone generated by the columns of $A^T$ is pointed. If not, we use the strategy that we developed for the removal of cycles to make the cone pointed. We will very shortly discuss the strategy here too, but it is so similar to the removal of cycles that we refer to Section 5.5 for details.

If there is a $\mathbf{0} \neq \boldsymbol{\lambda} \geq \mathbf{0}$ such that $A^T\boldsymbol{\lambda} = \mathbf{0}$, then the Linear Program in (54) is unbounded. We solve this by selecting one $\lambda_j > 0$ and take the corresponding row $\boldsymbol{A}_{j\bullet}$. Since the rows of $A$ correspond to inequalities, we can see that $\boldsymbol{A}_{j\bullet}\boldsymbol{x} \geq \mathbf{0}$ and $0 \leq \sum_{k\neq j} \lambda_k \boldsymbol{A}_{k\bullet}\boldsymbol{x} = -\lambda_j \boldsymbol{A}_{j\bullet}\boldsymbol{x}$, so that $\boldsymbol{A}_{j\bullet}$ in fact gives an equality constraint: $\boldsymbol{A}_{j\bullet}\boldsymbol{x} = 0$. This means that we can add or subtract $\boldsymbol{A}_{j\bullet}$ to other rows without affecting the inequality representation. We choose to pick a nonzero entry of $\boldsymbol{A}_{j\bullet}$, say $A_{jl}$ and cancel the $l$-th entry from each row in $A$. The row $\boldsymbol{A}_{j\bullet}$ itself will for now be stored. We can repeat this procedure until the resulting cone is pointed. Then, we can use the redundancy-removal outlined above, and after that we add the found equality constraints again.

##### 7.4 ineq($\mathcal{C}$) $\stackrel{\text{DD}}{\to}$ gen($\mathcal{C}$) = ECMs

In this last step, we start with the found inequality representation of the steady-state conversion cone. Because we have split all metabolites, this cone is contained in one orthant. This automatically implies that the cone must be pointed. We can thus simply apply the Double Description method (again using fractions) to find a minimal generating set of the steady-state conversion cone. After undoing the splitting of external metabolites (as mentioned in Section 7.3.2) this yields the Elementary Conversion Modes, finally.

#### 8 The direct method

The starting point of the direct method is the generator description of the (non-steady-state) conversion cone, $\mathcal{C}_0$, defined in (45) in Section 6. We assume that cycles have been removed using the method described in Section 5.5 such that this cone is pointed. The remaining task is thus to impose the steady-state constraints $\dot{c}_i = 0$ for all internal metabolites $i$. Whereas the indirect method computes an inequality representation of $\mathcal{C}_0$ before imposing the steady-state constraints, the direct method will impose these constraints 'directly' on the set of generators. As far as we know, this direct intersection method is unconventional. It has probably not been used before because it is usually slower than the indirect intersection method. It however avoids the (too) high memory usage that the indirect method sometimes suffers from.

The direct method proceeds by an iterative procedure. We start by gathering the generators from the generator description of $\mathcal{C}_0$ in a matrix $R^{(0)}$, and then impose an additional steady-state constraint in each iteration. We assume for now that we have ordered the metabolites such that there are $K$ internal metabolites with indices $\{1, \dots, K\}$. In the first step, we impose the constraint $\dot{c}_1 = 0$. The result is a new cone, called $\mathcal{C}^{(1)}$ which again has a set of generators, which we gather in the matrix $R^{(1)}$. We continue to add the constraints until we end up with the generators gathered in $R^{(K)}$ of the steady-state conversion cone: $\mathcal{C} = \mathcal{C}^{(K)}$.

In the following we will denote by $\boldsymbol{r}_j \in R$ the $j$-th column of the matrix $R$.

#### 8.1 Imposing one steady-state constraint

The $i$-th iteration of the algorithm starts with the matrix $R^{(i-1)}$ of non-redundant columns. These columns can be interpreted as conversions that satisfy the steady-state constraints for all metabolites with index smaller than $i$. The entry $R_{ij}^{(i-1)}$ thus indicates the net production of metabolite $i$ in the $j$-th conversion. During this step, we will impose the constraint $\dot{c}_i = 0$, which thus means that we should make sure that $R_{ij}^{(i)} = 0$ for all $j$. For this, similar to the Double Description method (Section 1.5), we compute the index sets

$$\begin{aligned} J^{+} &= \{j \in \{1, \dots, n\} : R_{ij}^{(i-1)} > 0\}, \\ J^{-} &= \{j \in \{1, \dots, n\} : R_{ij}^{(i-1)} < 0\}, \\ J^{0} &= \{j \in \{1, \dots, n\} : R_{ij}^{(i-1)} = 0\}. \end{aligned} \tag{55}$$

We are certain that the vectors corresponding to $J^0$ will be minimal generators of $\mathcal{C}^{(i)}$: they satisfy the constraint $\dot{c}_i = 0$ and cannot be written as a conical combination of the other vectors, since otherwise they would not have been in $R^{(i-1)}$. Furthermore, each pair of indices $j_+ \in J^+$, $j_- \in J^-$ with their corresponding vectors $\boldsymbol{r}_{j_+}, \boldsymbol{r}_{j_-}$ generate a possible candidate: $\hat{\boldsymbol{r}} = R_{ij_+}^{(i-1)} \boldsymbol{r}_{j_-}^{(i-1)} - R_{ij_-}^{(i-1)} \boldsymbol{r}_{j_+}$. We do not have to consider combinations of more vectors, since these can always be written as combinations of the pairs.

We could get a generating set of $\mathcal{C}^{(i)}$ by taking the union of all $\hat{\boldsymbol{r}}$ with the vectors from $J^0$, but this set would not be minimal. Therefore, we need a test to determine which candidates to keep, analogous to the adjacency test that is used in the Double Description method, Section 1.4. The adjacency test of the Double Description method however uses information about which inequalities are satisfied with equality by the generators to determine if a pair is adjacent or not. Since we intersect with equalities instead of inequalities, we cannot directly use this method. We would like to find an alternative way to determine the adjacency of two rays, using only the ray representation $R$.

The next section provides this alternative adjacency test. In short: we test whether two rays are adjacent by considering a point in between them. If this point can also be formed by a conic combination that includes other rays than the original two, then they are not adjacent.

##### 8.1.1 A geometric adjacency test

Let

$$\boldsymbol{x} = \frac{1}{4}\boldsymbol{r}_{j_+} + \frac{3}{4}\boldsymbol{r}_{j_-}$$

and consider the linear program, LP$(\boldsymbol{r}_{j_+}, \boldsymbol{r}_{j_-})$:

$$\begin{aligned} \underset{\boldsymbol{\lambda}}{\text{maximize}} \quad & \sum_{j \notin \{j_+, j_-\}} \lambda_i \\ \text{subject to} \quad & R\boldsymbol{\lambda} = \boldsymbol{x} \\ & \lambda_i \geq 0. \end{aligned} \tag{56}$$

Note that the LP is always feasible, and has optimal value at least 0, since we have the initial solution $\bar{\boldsymbol{\lambda}}$ with $\bar{\lambda}_{j_+} = 1/4$, $\bar{\lambda}_{j_-} = 3/4$ and zero in the other coordinates. We will prove in the following theorem that this LP can be used as an adjacency test. The reason for choosing 1/4 and 3/4 is that this ascertains that the 'target' $\boldsymbol{x}$ is not close to the zero vector. In fact, in `ecmtool` we normalize the rays before starting the LP such that the $L^1$-norm of all rays is equal to 1. We can then show with the reverse triangle inequality that

$$||\boldsymbol{x}|| = \left|\left|\frac{1}{4}\boldsymbol{r}_{j_+} - (-\frac{3}{4}\boldsymbol{r}_{j_-})\right|\right| \geq \left|\frac{1}{4}||\boldsymbol{r}_{j_+}|| - \frac{3}{4}||\boldsymbol{r}_{j_-}||\right| = \left|-\frac{1}{2}\right| = \frac{1}{2}. \tag{57}$$

In the following we will denote by $R$ the matrix of generators (columns $\boldsymbol{r}_j$), and by $A$ the inequality matrix of the same polyhedral cone $P(A)$. Recall from Section 1.4 that the zero set of a generator was defined as

$$Z(\boldsymbol{r}_j) = \{i : \boldsymbol{A}_{i\bullet}\boldsymbol{r}_j = 0\}.$$

**Theorem 8** (geometric adjacency test)**.** *The following are equivalent for extreme rays* $\boldsymbol{r}_{j_+}, \boldsymbol{r}_{j_-}$*:*

*(1)* $\boldsymbol{r}_{j_+}$ *and* $\boldsymbol{r}_{j_-}$ *are adjacent.*

*(2)* $Z(\boldsymbol{r}_{j_+}) \cap Z(\boldsymbol{r}_{j_-}) \subseteq Z(\boldsymbol{r}_k) \implies \boldsymbol{r}_k \sim \boldsymbol{r}_{j_+}$ *or* $\boldsymbol{r}_k \sim \boldsymbol{r}_{j_-}$*.*

*(3) LP*$(\boldsymbol{r}_{j_+}, \boldsymbol{r}_{j_-})$ *has optimal value 0.*

*Proof.* The equivalence of (1) and (2) is just the definition of adjacency described in Section 1.4.

**(2)** $\implies$ **(3)**. Suppose (2) holds, so $Z(\boldsymbol{r}_{j_+}) \cap Z(\boldsymbol{r}_{j_-}) \subseteq Z(\boldsymbol{r}_k) \implies \boldsymbol{r}_k \sim \boldsymbol{r}_{j_+}$ or $\boldsymbol{r}_k \sim \boldsymbol{r}_{j_-}$. Consider any extreme ray $\boldsymbol{r}_k$ that is not equivalent to $\boldsymbol{r}_{j_+}$ or $\boldsymbol{r}_{j_-}$. We know that $Z(\boldsymbol{r}_k)$ does not contain $Z(\boldsymbol{r}_{j_+}) \cap Z(\boldsymbol{r}_{j_-})$, so there is some index $i \in Z(\boldsymbol{r}_{j_+}) \cap Z(\boldsymbol{r}_{j_-})$ that is not in $Z(\boldsymbol{r}_k)$. This means $\boldsymbol{A}_{i\bullet}\boldsymbol{r}_k > 0$.

Suppose $\boldsymbol{\lambda}$ is a feasible solution for LP$(\boldsymbol{r}_{j_+}, \boldsymbol{r}_{j_-})$, so $R\boldsymbol{\lambda} = \boldsymbol{x}$. Now consider

$$\boldsymbol{A}_{i\bullet} R\boldsymbol{\lambda} = \sum_j \boldsymbol{A}_{i\bullet}\boldsymbol{r}_j \lambda_j \geq \boldsymbol{A}_{i\bullet}\boldsymbol{r}_k \lambda_k. \tag{58}$$

The last inequality holds because each ray $\boldsymbol{r}_j$ is in the cone; hence $\boldsymbol{A}_{i\bullet}\boldsymbol{r}_j \geq 0$, and also $\lambda_j \geq 0$ (one of the LP constraints). At the same time

$$\boldsymbol{A}_{i\bullet} R\boldsymbol{\lambda} = \boldsymbol{A}_{i\bullet}\boldsymbol{x} = \boldsymbol{A}_{i\bullet}(\frac{1}{4}\boldsymbol{r}_{j_+} + \frac{3}{4}\boldsymbol{r}_{j_-}) = 0, \tag{59}$$

since $i \in Z(\boldsymbol{r}_{j_+})$ and $i \in Z(\boldsymbol{r}_{j_-})$. Combining (58) and (59) with $\boldsymbol{A}_{i\bullet}\boldsymbol{r}_k > 0$ gives $\lambda_k = 0$. Because $\boldsymbol{r}_k$ was any extreme ray not equivalent to $\boldsymbol{r}_{j_+}$ or $\boldsymbol{r}_{j_-}$, and $\boldsymbol{\lambda}$ was any feasible solution, this means that we will always have $\sum_i \lambda_i - \lambda_a - \lambda_b = 0$, hence (3) holds.

**(3)** $\implies$ **(2)**. For a contrapositive proof, assume (2) does not hold. Then there is some ray $\boldsymbol{r}_k$ such that $Z(\boldsymbol{r}_{j_+}) \cap Z(\boldsymbol{r}_{j_-}) \subseteq Z(\boldsymbol{r}_k)$. By definition $\boldsymbol{x} = 1/4\boldsymbol{r}_{j_+} + 3/4\boldsymbol{r}_{j_-}$. Therefore, for any row $\boldsymbol{A}_{i\bullet}$,

$$\boldsymbol{A}_{i\bullet}\boldsymbol{x} = \boldsymbol{A}_{i\bullet}(\frac{1}{4}\boldsymbol{r}_{j_+} + \frac{3}{4}\boldsymbol{r}_{j_-}) = \frac{1}{4}\boldsymbol{A}_{i\bullet}\boldsymbol{r}_{j_+} + \frac{3}{4}\boldsymbol{A}_{i\bullet}\boldsymbol{r}_{j_-}. \tag{60}$$

Because $\boldsymbol{r}_{j_+}$ and $\boldsymbol{r}_{j_-}$ are in the cone, $\boldsymbol{A}_{i\bullet}\boldsymbol{r}_{j_+} \geq 0$ and $\boldsymbol{A}_{i\bullet}\boldsymbol{r}_{j_-} \geq 0$. Thus $\boldsymbol{A}_{i\bullet}x = 0$ if and only if $\boldsymbol{A}_{i\bullet}\boldsymbol{r}_{j_+} = \boldsymbol{A}_{i\bullet}\boldsymbol{r}_{j_-} = 0$, hence

$$Z(\boldsymbol{x}) = Z(\boldsymbol{r}_{j_+}) \cap Z(\boldsymbol{r}_{j_-}). \tag{61}$$

Consider the line segment

$$L(t) = t\boldsymbol{x} + (1-t)\boldsymbol{r}_k \tag{62}$$

for $t \in [0, 1]$. This is the line segment from $\boldsymbol{r}_k$ to $\boldsymbol{x}$. In Figure S8 we have schematically drawn the situation. Take any index $l$ not in $Z(\boldsymbol{x})$. Since $\boldsymbol{A}_{l\bullet}\boldsymbol{x} > 0$, we have, for any $t \in (0,1]$,

$$\boldsymbol{A}_{l\bullet}L(t) = \boldsymbol{A}_{l\bullet}t\boldsymbol{x} + \boldsymbol{A}_{l\bullet}(1-t)\boldsymbol{r}_k > 0. \tag{63}$$

Because $\boldsymbol{A}_{l\bullet}L(t)$ is continuous with respect to $t$, there is an $\varepsilon_l > 0$ such that $\boldsymbol{A}_{l\bullet}L(1+\varepsilon_l) > 0$. Define $\varepsilon$ as the minimum of all the $\varepsilon_l$ obtained in this way, then

$$\boldsymbol{A}_{l\bullet}L(1+\varepsilon) > 0 \text{ for all } l \text{ not in } Z(\boldsymbol{x}) \tag{64}$$

On the other hand, any index $m \in Z(\boldsymbol{x})$ is also in $Z(\boldsymbol{r}_k)$, since $Z(\boldsymbol{x}) = Z(\boldsymbol{r}_{j_+}) \cap Z(\boldsymbol{r}_{j_-}) \subseteq Z(\boldsymbol{r}_k)$. This yields

$$\boldsymbol{A}_{m\bullet}L(1+\varepsilon) = \boldsymbol{A}_{m\bullet}(1+\varepsilon)\boldsymbol{x} - \boldsymbol{A}_{m\bullet}\varepsilon\boldsymbol{r}_k = 0 \text{ for all } m \in Z(\boldsymbol{x}). \tag{65}$$

Let $\boldsymbol{\alpha} = L(1+\varepsilon)$, it follows from (64) and (65) that $\boldsymbol{\alpha} \in P(A)$ and $Z(\boldsymbol{\alpha}) = Z(\boldsymbol{x})$. We can write

$$\boldsymbol{\alpha} = L(1+\varepsilon) = (1+\varepsilon)\boldsymbol{x} - \varepsilon\boldsymbol{r}_k, \tag{66}$$

so that

$$\boldsymbol{\alpha} + \varepsilon\boldsymbol{r}_k = (1+\varepsilon)\boldsymbol{x}, \tag{67}$$

and

$$\frac{\boldsymbol{\alpha}}{1+\varepsilon} + \frac{\varepsilon\boldsymbol{r}_k}{1+\varepsilon} = \boldsymbol{x}. \tag{68}$$

Because $\boldsymbol{\alpha}$ is a point in $P(A)$, it can be written as a conic combination of rays, say $\boldsymbol{\alpha} = \sum_i \lambda_i \boldsymbol{r}_i$. But then $\frac{1}{1+\varepsilon}\boldsymbol{\lambda} + \frac{\varepsilon}{1+\varepsilon}\hat{\boldsymbol{e}}_k$ is a feasible solution to the LP in (56), with objective value

$$\sum_{i \notin j_+, j_-} \frac{1}{1+\varepsilon}\lambda_i + \frac{\varepsilon}{1+\varepsilon} \geq \frac{\varepsilon}{1+\varepsilon} > 0. \tag{69}$$

So, LP$(\boldsymbol{r}_{j_+}, \boldsymbol{r}_{j_-})$ does not have optimal value 0. By contrapositivity, this implies that whenever the optimal value is 0, $\boldsymbol{r}_{j_+}$ and $\boldsymbol{r}_{j_-}$ must be adjacent. □

This theorem showed that solving the right LP can serve as an adjacency test for extreme rays. The following theorem in addition shows that if the rays are not adjacent, the original rays cannot be part of the optimal solution to the LP. To be precise: if the rays $\boldsymbol{r}_{j_+}, \boldsymbol{r}_{j_-}$ are not adjacent, then at some point the contribution $\lambda_{j_+}$ or $\lambda_{J_-}$ must become zero. This is important to make the adjacency test numerically robust, because the change in $\boldsymbol{\lambda}$ has to be of size at least 1/4 which means that it is easy to distinguish from round-off errors. Moreover, it provides an early exit strategy if we use our LP-solver described in

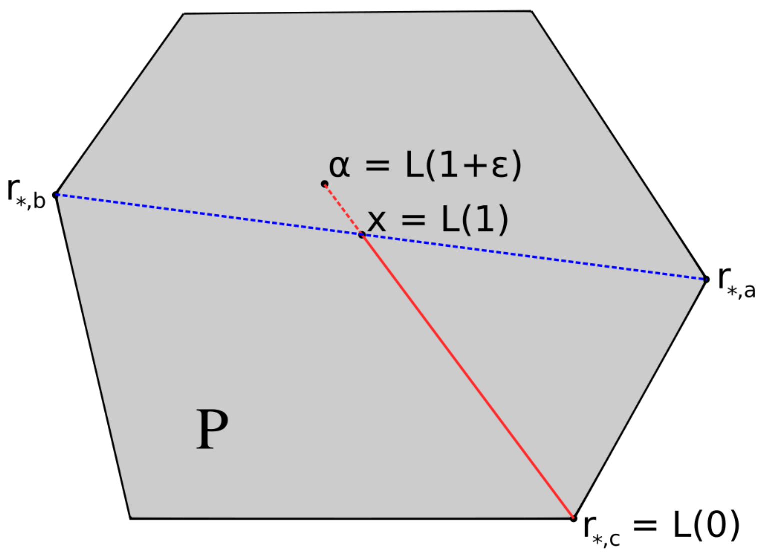

Figure S8: Illustration for the proof of **(3)** $\implies$ **(2)**. The point $\boldsymbol{\alpha}$ will always be inside $P$, indicating a feasible solution to the LP with objective strictly greater than 0. The grey area might not look like a cone at first, but consider it as a cross section of one.

Section 4: at the point that $\lambda_{j_+}$ or $\lambda_{j_-}$ becomes zero, we know that the rays are non-adjacent, and we can stop the LP.

**Theorem 9.** *Let $\boldsymbol{\mu}$ be the optimal solution to LP($\boldsymbol{r}_{j_+}, \boldsymbol{r}_{j_-}$). Then at least one of $\mu_{j_+}$ and $\mu_{j_-}$ is equal to zero.*

*Proof.* Suppose that both $\mu_{j_+} > 0$ and $\mu_{j_-} > 0$. We will show that this contradicts the optimality of $\boldsymbol{\mu}$. Besides $\boldsymbol{\mu}$ we know of one more feasible solution: the vector $\boldsymbol{\nu}$ with all zeroes except for $\nu_{j_+} = 1/4$ and $\nu_{j_-} = 3/4$. Since both $\boldsymbol{\mu}$ and $\boldsymbol{\nu}$ are feasible solutions, we have $R(\boldsymbol{\mu} - \boldsymbol{\nu}) = 0$. Now consider $\bar{\boldsymbol{\mu}} = \boldsymbol{\mu} + \delta(\boldsymbol{\mu} - \boldsymbol{\nu})$ for some small $\delta > 0$. This $\bar{\boldsymbol{\mu}}$ satisfies

$$R\bar{\boldsymbol{\mu}} = R(\boldsymbol{\mu} + \delta(\boldsymbol{\mu} - \boldsymbol{\nu})) = R\boldsymbol{\mu} + \delta R(\boldsymbol{\mu} - \boldsymbol{\nu}) = R\boldsymbol{\mu} + \mathbf{0} = \boldsymbol{x}. \tag{70}$$

The only positions in $(\boldsymbol{\mu} - \boldsymbol{\nu})$ that could be negative are $j_+$ and $j_-$, since $\boldsymbol{\nu}$ is zero everywhere else. But we know that $\mu_{j_+}, \mu_{j_-} > 0$, so if we pick $\delta$ small enough then $\bar{\mu}_{j_+}, \bar{\mu}_{j_-} \geq 0$, so that

$$\bar{\boldsymbol{\mu}} \geq \mathbf{0}. \tag{71}$$

Together (70) and (71) show that $\bar{\boldsymbol{\mu}}$ is a feasible solution. Now let us denote by Obj($\boldsymbol{\mu}$) the objective value corresponding to the solution $\boldsymbol{\mu}$. We can see that

$$\text{Obj}(\bar{\boldsymbol{\mu}}) = \sum_{i \notin \{j_+, j_-\}} \bar{\mu}_i = \sum_{i \notin \{j_+, j_-\}} \mu_i + \delta(\mu_i - \nu_i) = (1+\delta) \sum_{i \notin \{j_+, j_-\}} \mu_i > \sum_{i \notin \{j_+, j_-\}} \mu_i = \text{Obj}(\boldsymbol{\mu}). \quad (72)$$

This would contradict the optimality of $\boldsymbol{\mu}$, so it must be that in the optimal solution $\mu_{j_+} = 0$ or $\mu_{j_-} = 0$. □

##### 8.1.2 Performing the adjacency tests

Recall that one iteration of the direct method involves finding a minimal generating set of the cone $\mathcal{C}^{(i)}$ from the generating set $R^{(i-1)}$ of $\mathcal{C}^{(i-1)}$. For that, we determined for all generating vectors in $R^{(i-1)}$ the sign of $\dot{c}_i$, and used that to construct the sets $J^+, J^-, J^0$ (see (55)). All pairs with one ray from $J^+$ and one ray from $J^-$ give a potential candidate that we should test for redundancy. The candidate will only be non-redundant if the original rays $\boldsymbol{r}_{j_+}, \boldsymbol{r}_{j_-}$ are adjacent, according to the adjacency test defined above. To test all these candidates, we thus have to perform $|J^+||J^-|$ Linear Programs.

Fortunately, this LP is exactly of the form described in Section 4. We thus already have an efficient way to solve the LP. For this LP-solver to work, we do need to select a maximal set of linearly independent columns of $R$ as a starting basis. This basis should contain the rays $\boldsymbol{r}_{j_+}, \boldsymbol{r}_{j_-}$ that constitute an initial solution. Finding this starting basis is computationally relatively complex. Note however, that we use the same matrix $R$ for each redundancy test, and we can therefore use almost the same starting basis, $B$. We should only make sure that we replace two of the columns in $A_B$ by $\boldsymbol{r}_{j_+}, \boldsymbol{r}_{j_-}$. In Section 4.5 we describe how we can add one such column. For this we need to solve $A_B\boldsymbol{x} = \boldsymbol{r}_{j_+}$. Call the basis where this column is added $A_B^+$. Now, we should still add the second column, and for that solve $A_B^+\boldsymbol{x} = \boldsymbol{r}_{j_-}$.

In `ecmtool` we optimize this even further. We start by selecting a basis matrix $A_B$ once, and immediately calculate its LU-decomposition. This LU-decomposition can be used to solve $A_B\boldsymbol{x} = \boldsymbol{r}_{j_+}$. We solve this system for all possible $\boldsymbol{r}_{j_+}$, giving a set of bases $A_B^+$. For each basis in this set we also calculate the LU-decomposition once. These can then be used to solve the system $A_B^+\boldsymbol{x} = \boldsymbol{r}_{j_-}$ for all possible $\boldsymbol{r}_{j_-}$, yielding all the bases that we will need. This strategy requires us to calculate $1 + |J^+|$ LU-decompositions, instead of $|J^+||J^-|$, which induces a relevant computational speedup.

Another important optimization that decreases the computational time needed by the direct method, is that we perform the above described LPs using floats instead of fractions. We can do this because each LP will only give a boolean output, indicating if a pair of rays is adjacent, or not. We use fractions again when the adjacent rays are combined to form a generator that satisfies the steady-state constraints, so that the eventual ECM-computation remains exact.

#### 8.2 The order of imposing steady-state constraints

As mentioned above, in each iteration we impose one of the steady-state constraints $\dot{c}_i = 0$. The order in which these constraints are imposed has a large effect on the total computation time. This is similar to the sensitivity of the Double Description method to changing the order in which inequalities are added [?]. It is however unclear which order of equality intersection minimizes the computation time. Therefore, we offer several heuristics as options using the argument `--sort_order`.

The sorting order that usually performs well is `--sort_order min_adj`. Here, we first sum how many metabolites are adjacent in the metabolic network, i.e., connected by one reaction. Then, for each $i$, we sum how many metabolites would become adjacent if we would connect all reactions that produce metabolite $i$ to the reactions that consume metabolite $i$. As such, we try to estimate the number of adjacencies that are added, i.e., the difference between the number of adjacent metabolites after and before the removal of metabolite $i$. Note that the number of added adjacencies can be negative, because metabolite $i$ is deleted and can thus no longer be adjacent to any metabolite. If this sorting order is chosen, `ecmtool` picks the metabolite that minimizes the number of added adjacencies. The intuition behind this sort order is that it would lead to summarizing possible modules in the metabolic network before connecting these to the rest of the network. Indeed, if we use this sorting order, the last steps are often not the largest.

An alternative heuristic that `ecmtool` offers is `--sort_order min_lp`. This selects the metabolite $i$ that requires the minimal number of LPs. Recall that to impose the $i$-th steady-state constraint, we need to solve $|J^+||J^-|$ Linear Programs, where the size of the index sets $J^+, J^-$ is given by how many of the current generators produce/consume metabolite $i$ (see 55). This heuristic thus selects metabolites that are produced and consumed by few reactions. This has the disadvantage of removing the 'easy' metabolites first from the network, which could lead to a very difficult step later on.

#### 8.3 Parallelization

Since the adjacency tests are independent of each other, and we commonly need to perform millions of them to eliminate a single metabolite (for genome-scale networks) this algorithm is highly suitable for parallel processing. We have implemented this using `mpi4py` [?].

In Table 1 and Figure S9 we show the relative speed-up compared to using a single CPU for the `e_coli_core`-model. It shows close to linear gains. This is a smaller network, where the most LPs done in a single step is around $10e5$, so for genome scale networks we can expect the scaling to continue up to hundreds or more CPUs.

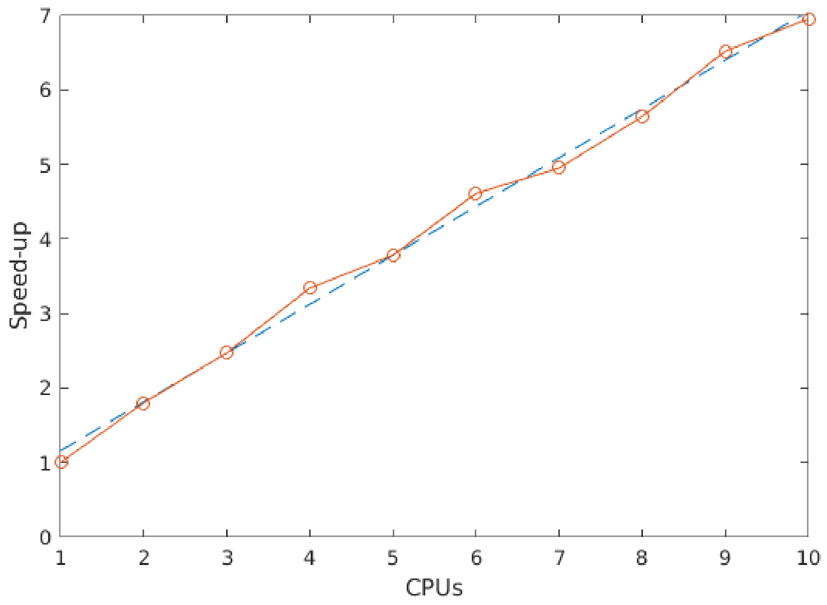

Figure S9: CPU count vs speed-up (red) and linear least-squares fit (blue, dashed). The fit is $y = 0.655x + 0.503$.

Table 1: Speed-up with different processor counts for *E. coli* core

| **CPUs** | 2 | 3 | 4 | 5 | 6 | 7 | 8 | 9 | 10 |
|---|---|---|---|---|---|---|---|---|---|
| **Speed-up rel. to 1 CPU** | 1.80 | 2.47 | 3.34 | 3.78 | 4.61 | 4.95 | 5.64 | 6.52 | 6.95 |

### 9 Validation of `ecmtool`

In this section, we describe how we used a `Matlab`-script to validate that the ECMs calculated by `ecmtool` for the `e_coli_core`-network satisfy

1. each ECM is an elementary vector
2. each steady-state conversion must be a conical combination of ECMs

1. According to the definition of ECMs given in Section 2, we can prove that each ECM is elementary by showing that it cannot be written as a positive sum of the other ECMs without the production of any external metabolite being canceled. We tested this with the `Matlab`-script `lp_ecms_efm.m`, which is available as a Supplementary File.

For each ECM, $\boldsymbol{x}$, we first gather all other ECMs that are in the same orthant as columns in a matrix

$R$. Then, we solve the following LP:

$$\begin{aligned} \underset{\boldsymbol{\lambda}}{\text{minimize}} \quad & \sum_i \lambda_i \\ \text{subject to} \quad & \begin{bmatrix} R \\ -R \end{bmatrix} \boldsymbol{\lambda} = \begin{bmatrix} \boldsymbol{x} + tol \\ -\boldsymbol{x} + tol \end{bmatrix} \\ & 0 \leq \lambda_i \leq 10^3. \end{aligned} \tag{73}$$

If the ECM is indeed an elementary vector, this LP should be infeasible. However, since the LP in `Matlab` is solved using floats, we should pay attention if round-off errors do not lead to incorrect results. Therefore, we allow for the decomposition to be off by a certain tolerance $tol$. If we allowed for an error of $10^{-7}$ none of the ECMs could be decomposed, indicating that they are all elementary. However, when we allowed for a larger error, more ECMs could be decomposed. This gives an indication of how close a conical combination of different ECMs can come to replacing an ECM. In Figure S10 we show the distribution of error margins that are needed to decompose an ECM into different ECMs. We see that most of the ECMs cannot be written as a combination of others unless one allows for an error of $10^{-5}$ or larger. This indicates that the ECMs are indeed all elementary vectors.

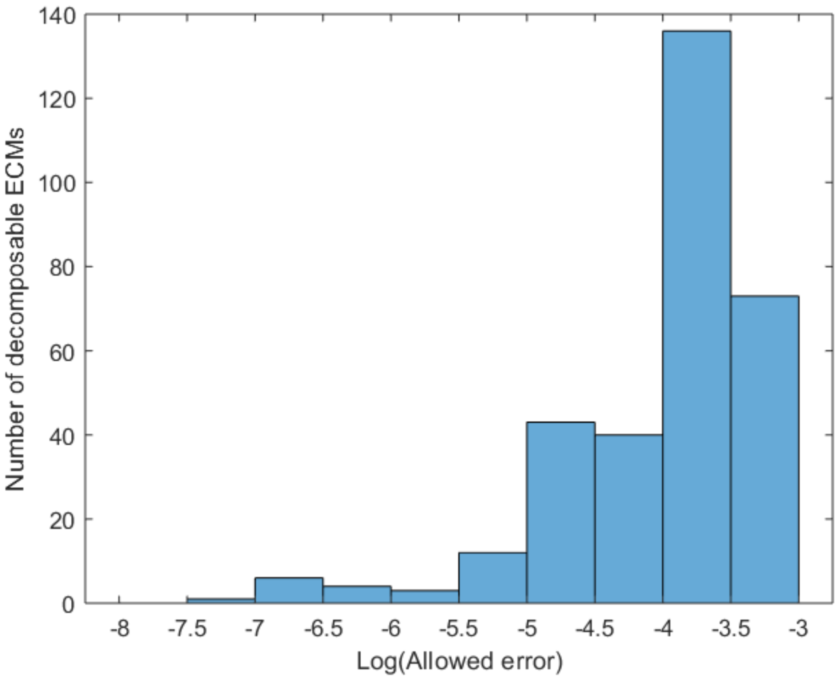

Figure S10: The distribution of error margins, $tol$ that are needed to decompose an ECM into different ECMs.

2. It is of course hard to validate that any steady-sate conversion is a conical combination of ECMs. We therefore chose to use the set of Elementary Flux Modes calculated by `efmtool`. This set spans all possible steady-state flux combinations that the model contains. For each EFM, we then calculated its overall conversion, and tried to write this conversion as a combination of ECMs, using a similar LP as in (73). If we allowed for an error of $10^{-7}$, then each EFM-based-conversion could be decomposed into ECMs. This error margin was necessary because the results from `efmtool` were affected by round-off errors, while the ECMs are calculated using fractions and are therefore exact.

When we multiply the EFMs with the stoichiometry matrix, we map them into the conversion cone. Many of the EFMs will end up in the interior of the cone, thus leading to conversions that are combinations of ECMs. However, for each ECM there must be at least one EFM that leads to exactly that conversion. As a last sanity check, we tried to validate this. We took the decompositions of EFMs into ECMs obtained in the LPs above, and checked if each ECM occurs at least once as the sole decomposing vector of an EFM. We say that an ECM is a decomposing vector of an EFM if it is used more than some 'support tolerance'. In Figure S11, we vary this tolerance. We see that at a support tolerance of $10^{-2}$, all ECMs have at least one EFM of which they are the sole decomposing ECM. Since both ECMs and EFMs were normalized before this validation, this implies that for all ECMs there is an EFM-based-conversion that is 99% equal to the ECM. This indicates that this EFM actually corresponds to the same conversion as the ECM; the 1% is caused by round-off errors and the allowed tolerance *tol* in the LP.

1. Description of validation on the `e_coli_core`-network

2. Figures of error distribution in decomposing EFMs

### 10 ECM-analyses several networks

In the main text we report on ECM computations for the `e_coli_core`-network [?], the `iIT341`-network [?], and the `iJR904`-network [?]. The full results and the run scripts that were used to obtain these results have been uploaded as supplementary files, and can also be found in the Github-folder:
`https://github.com/SystemsBioinformatics/ecmtool`, specifically in the subfolder
`results_and_corresponding_runscripts/`.

The Elementary Conversion Modes for the rhizobial bacteroids were calculated on the model `iCS320` by Schulte et al. The results and run scripts are reported in their work [?].

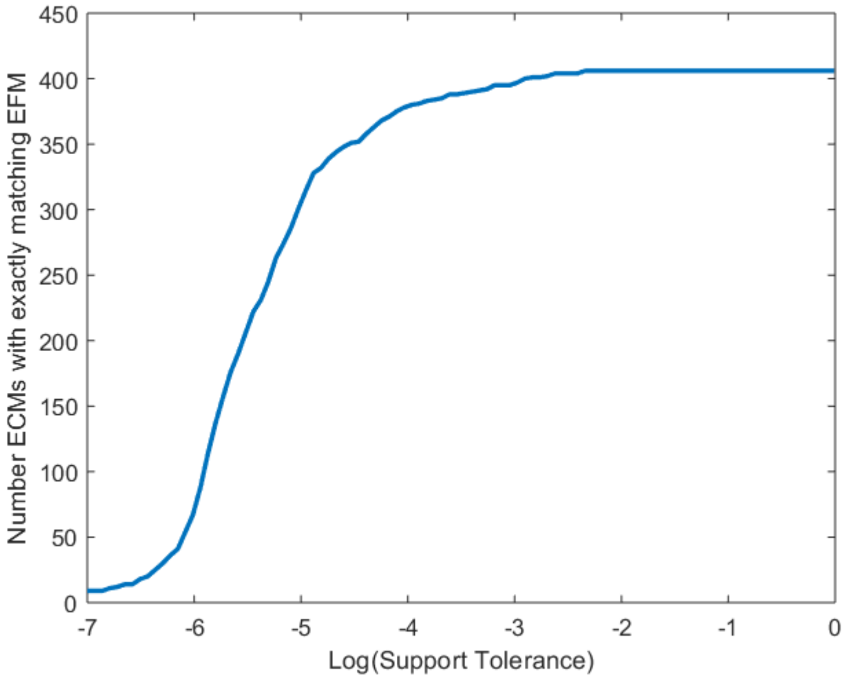

Figure S11: The cumulative number of ECMs that are identified as the sole decomposing ECM of at least one EFM, when we vary the cut-off at which we mark an ECM as 'decomposing'.

#### 10.1 Creating subnetworks of `e_coli_core`

In one of the figures of the main text we describe how the number of ECMs and EFMs behaves for various subnetworks of the `e_coli_core`-model. These subnetworks were created with a Python-script that we called `subnetwork_creator.py`, which is attached as a supplementary file. We created the subnetworks via an iterative procedure. The smallest subnetwork is constructed by taking only the active reactions in the FBA-solution. After that, we made a series of knockout-models. We deleted one of the active reactions, and again ran a Flux Balance Analysis. This knockout-model necessarily had a new set of active reactions. We took the union of these active reactions with the original active reactions to create our second model. Then, we again deleted a reaction, did another FBA, and took the active reactions. As such, we created subnetworks of increasing size for which we could compute the ECMs.

#### 10.2 Clustering the ECM results for visualisation

We have clustered some of the ECM-enumeration results for visualization purposes. All `R`-scripts are made available as supplementary files. We first made sure that the set of ECMs was no larger than a few thousand. For some models, comprising hundreds of thousands ECMs, we had to take a subset of ECMs with a certain property, for example growth-supporting ECMs. Given this set, we created a distance matrix that contains the $L^1$-distance between all pairs of ECMs. For some models, we chose to weigh the $L^1$-distance so that some metabolites are considered more important than others. On this distance matrix, we performed hierarchical clustering. The metabolites were ordered from top to bottom by the number of ECMs that used the metabolite as an output minus the number of ECMs that used the metabolite as an input.

To visualize the clustered ECMs we used two options. For some models we converted all coefficients to a ternary scale, showing only whether the metabolite in the ECM was taken up, left untouched, or secreted. For other models we used a shifted logarithmic scale. To be precise, we converted the stoichiometric coefficients according to:

$$x = \begin{cases} \log\left(\frac{x}{shift_{pos} \cdot minpos}\right) & \text{if } x > 0, \\ 0 & \text{if } x = 0, \\ -\log\left(\frac{x}{shift_{neg} \cdot maxneg}\right) & \text{if } x < 0, \end{cases} \tag{74}$$

where $shift_{neg}, shift_{pos}$ are parameters smaller than 1, and $minpos, maxneg$ are respectively the smallest positive and the largest negative coefficient occurring in the ECMs. This transformation was necessary to visualize all differences in the coefficients occuring in the ECMs, because these coefficients span many orders of magnitudes and are both positive and negative. However, we should emphasize that this transformation can be used for visualization purposes only.

### 11 User guide

#### 11.1 Prerequisite ingredients for ECM-computation

To compute the ECMs, one needs to provide at least an SBML-model. From the SBML-model, the following will be extracted by `ecmtool`

1. a stoichiometry matrix,
2. reversibility information of all reactions,
3. information on which metabolites are external or internal,
4. information on whether external metabolites can be produced, consumed or both

###### Important notes for the correct parsing of SBML-files by ecmtool

It should be checked carefully that `ecmtool` has parsed the model corresponding to the user's intentions. By far the most issues that users may have with ecmtool are **due to incorrect parsing of the SBML-**

**file**. Because several conventions exist for storing several features of the model, `ecmtool` cannot comply with all of them. When `ecmtool` is used as a standalone command line tool the parsing result can be checked by running `ecmtool` with the arguments `--print_reactions True` and `--print_metabolites True`, as described in subsection 11.2.1 below. When `ecmtool` is used as a Python library, the parsing result is available in the variable of the `network` class. Important to check are at least:

1. **reversibility information of all reactions**. We use the convention that reactions that are marked as irreversible can *only run in the forward direction.* Reactions can thus *not be backward irreversible.* In this case, the direction of the reaction should be swapped.

2. **internal/external-information of metabolites**. We use the convention that the metabolite-IDs of external metabolites are marked by `_e`. The user can change this with the argument `--external_compartment`. In addition, exchange reactions and external metabolites are recognized using functionality from the `cbmpy`-library, but this might not catch all.

3. **directionality information of external metabolites**. Based on the direction and reversibility of exchange reactions we determine whether a metabolite can be used as an input, an output or as both. This is what is most often parsed erroneously, due to conflicting conventions about when to set a reaction as reversible/irreversible in relation to its flux bounds.

`Ecmtool` can be used in two different modes: either as a standalone command line tool, or as a Python library for your own scripts. This section describes how to install and use both modes.

#### 11.2 Mode 1: standalone command line tool

In this mode, you can call ecmtool like a normal program from your command line. It reads metabolic networks in the SBML format, and writes resulting ECMs into a CSV file for later analysis. Most researchers will use this method. For running `ecmtool` on computing clusters efficiently, see the Advanced Usage section in this readme.

###### Installation

- Download and install Python. `Ecmtool` is compatible with python 3.x. Ensure both python and its package manager *pip* are added to your PATH environment variable. If this last step is omitted, an error like the following will be thrown when you try to run python: *'python' is not recognized as an internal or external command [..].*

- Download the latest `ecmtool` source through *git clone*, or as a zip file from `https://github.com/tjclement/ecmtool`.

- Open a command prompt, and navigate to the ecmtool directory (e.g. *cd C:\Users\You\Git\ecmtool*, where the path should be replaced with the path `ecmtool` was downloaded to).
- Install the dependencies in requirements.txt inside the ecmtool directory (e.g. by running *pip install -r requirements.txt*).
- Linux only: install *redund* of package `lrslib` (e.g. by running *apt install lrslib*).

**Running**

`Ecmtool` can be ran by executing

```
python3 main.py {model_path <path/to/model.xml> [arguments]
```

from the command line, after navigating to the ecmtool directory as described above. The possible arguments and their default values are printed when you run `python main.py --help`. After execution is done, the found conversions have been written to file (default: *conversions.csv*). The first row of this CSV file contain the metabolite IDs as read from the SBML model.

##### 11.2.1 Optional arguments

- `--model_path, type=str, default='models/active_subnetwork_KO_5.xml`. Relative or absolute path to an SBML model (.xml-file)
- `--direct, type=str2bool, default=False`. Enable to intersect with equalities directly. Direct intersection works better than indirect when many metabolites are hidden, and on large networks (default: False)
- `--compress, type=str2bool, default=True`. Perform compression to which the conversions are invariant, and reduce the network size considerably (default: True)
- `--out_path, default='conversion_cone.csv'`. Relative or absolute path to the .csv file you want to save the calculated conversions to (default: `conversion_cone.csv`)
- `--add_objective_metabolite, type=str2bool, default=True`. Add a virtual metabolite containing the stoichiometry of the objective function of the model (default: true).
- `--print_metabolites, type=str2bool, default=True`. Print the names and IDs of metabolites in the (compressed) metabolic network (default: true)
- `--print_reactions, type=str2bool, default=False`. Print the names and IDs of reactions in the (compressed) metabolic network (default: true)

- `--print_conversions, type=str2bool, default=True`. Print the calculated conversion modes (default: true)
- `--external_compartment, type=str, default='e'`. String indicating how the external compartment in metabolite-ids of SBML-file is marked. Please check if external compartment detection works by checking metabolite information before compression and with `--print metabolites true`)
- `--auto_direction, type=str2bool, default=True`. Automatically determine external metabolites that can only be consumed or produced (default: true)
- `--inputs, type=str, default=''`. Comma-separated list of external metabolite indices, as given by `--print_metabolites true` (before compression), that can only be consumed
- `--outputs, type=str, default=''`. Comma-separated list of external metabolite indices, as given by `--print_metabolites true` (before compression), that can only be produced. If inputs are given, but no outputs, then everything not marked as input is marked as output. If inputs and outputs are given, the possible remainder of external metabolites is marked as both
- `--hide, type=str, default=''`. Comma-separated list of external metabolite indices, as given by `--print_metabolites true` (before compression), that are transformed into internal metabolites by adding bidirectional exchange reactions
- `--prohibit, type=str, default=''`. Comma-separated list of external metabolite indices, as given by `--print_metabolites true` (before compression), that are transformed into internal metabolites without adding bidirectional exchange reactions. This metabolite can therefore be used as neither input nor output.
- `--tag, type=str, default=''`. Comma-separated list of reaction indices, as given by `--print_reactions true` (before compression), that will be tagged with new virtual metabolites, such that the reaction flux appears in ECMs.
- `--hide_all_in_or_outputs, type=str, default=''`. String that is either empty, input, or output. If it is input or output, after splitting metabolites, all inputs or outputs are hidden (objective is always excluded.
- `--iterative, type=str2bool, default=False`. Enable iterative conversion mode enumeration (might help on large, dense networks) (default: false)

- `--only_rays, type=str2bool, default=False`. Enable to only return extreme rays, and not elementary modes. This describes the full conversion space, but not all biologically relevant minimal conversions. See: Clement, 2020 and Urbanczik, 2005.
- `--verbose, type=str2bool, default=True`. Enable to show detailed console output (default: true)
- `--splitting_before_polco, type=str2bool, default=True`. Enables splitting external metabolites by making virtual input and output metabolites before starting the computation. Setting to false would do the splitting after first computation step. Which method is faster is complicated and model-dependent. (default: true)
- `--redund_after_polco, type=str2bool, default=True`. (Indirect intersection only) Enables redundant row removal from inequality description of dual cone. Works well with models with relatively many internal metabolites, and when running parrallelised computation using MPI (default: true)
- `--scei, type=str2bool, default=True`. Enable to use SCEI compression (default: true)
- `--sort_order, type=str, default='min_adj'`. Order in which internal metabolites should be set to zero during direct intersection. Default is to minimize the added adjacencies, other options are: `min_lp`, `max_lp_per_adj`, `min_connections`
- `--intermediate_cone_path, type=str, default=''`. Filename where intermediate cone result can be found. If an empty string is given (default), then no intermediate result is picked up and the calculation is done in full.
- `--manual_override, type=str, default=''`. (Advanced option). Index indicating which metabolite should be intersected in first step. Can be used in combination with `--intermediate_cone_path` to pick a specific intersection at a specific step.

**Example**

```
python3 main.py --model_path models/e_coli_core.xml --auto_direction true --
    out_path core_conversions.csv
```

#### 11.3 Mode 2: Python library

`Ecmtool` can also be used as a separate programming interface from within your own Python code. To do so, install `ecmtool` using *pip* (e.g. *pip install ecmtool*). The most crucial method is ecmtool.conversion_cone:get_conversion_cone(), which returns the ECMs of a given stoichiometric matrix. For information on how to use advanced features like SBML parsing, network compression, and metabolite direction estimation, please see ecmtool/main.py.

**Example**

```python
from ecmtool.network import extract_sbml_stoichiometry
from ecmtool.conversion_cone import get_conversion_cone

network = extract_sbml_stoichiometry('models/sxp_toy.xml', add_objective=True)
stoichiometry = network.N

ecms = get_conversion_cone(stoichiometry, network.external_metabolite_indices(),
 network.reversible_reaction_indices(), network.input_metabolite_indices(),
 network.output_metabolite_indices())

```

#### 11.4 Advanced usage

After testing how the tool works, most users will want to run their workloads on computing clusters instead of on single machines. This section describes some of the steps that are useful for running on clusers

**Doubling direct enumeration method speed**

The direct enumeration method can be sped up by compiling our LU decomposition code with Cython. The following describes the steps needed on Linux, but the same concept also applies to Mac OS and Windows. First make sure all dependencies are satisfied. Then execute:

```
python3 setup.py build_ext --inplace

mv _bglu* ecmtool/
```

**Running on a computing cluster with mpiexec**

For example:

```
mpiexec -n 4 python3 main.py --model_path models/e_coli_core.xml
```

#### Examples of run commands and necessary computing power

In this part, we provide some descriptions on how the presented results were obtained. All ECMs that were computed are supplied as supplementary files.

##### *Escherichia coli*-model: e_coli_core

This model, downloadable from `bigg.ucsd.edu` [?], is excellent for getting to know the workings of `ecmtool`, because the runtime is quite short. An example runscript is given by:

```
python3 main.py --model_path models/e_coli_core.xml --auto_direction true --
    direct false --splitting_before_polco true
```

It is good to get a feel for the different enumeration options, such as `--direct`, `--splitting_before_polco` and `--redund_after_polco`. The enumeration should always give the ECMs that can also be found in the file `conversions_ecolicore.csv`.

In the main text, we also show the ECMs obtained for this model when all outputs were hidden. This can be achieved by running

```
python3 main.py --model_path models/e_coli_core.xml --direct False --
    hide_all_in_or_outputs output
```

In fact, this command provides a shortcut to focus on only inputs, but one could also obtain this result with giving all indices of output metabolites to the `--hide`-argument.

If run with the argument `--print_reactions true`, `ecmtool` prints an indexed list of reactions before starting the computation. This can be used if a specific reaction is of interest. For example, in the main text we showed results in which the activity of the pyruvate dehydrogenase-reaction was reported. In the printed list one can see that this is reaction 50. We can therefore run

```
python3 main.py --model_path models/e_coli_core.xml --direct False --
    hide_all_in_or_outputs output --tag 50
```

##### *Helicobacter pylori*-model: iIT341

The `iIT341`-model is also available at `bigg.ucsd.edu`. The enumeration of all ECMs of this model is quite computationally intense. The results shown in Supplementary Figure 1, and made available in `iIT_minII_fullconversioncone.zip`, were calculated on a Linux-based virtual computing cluster with the command:

```
mpiexec -n 4 python3 main.py
--model_path models/iIT341.xml --direct false
```

```
--inputs 139,262,28,294,300,306,314,231,35,350,259,261,22,356,334,93,293,271
--out_path iIT.csv
--outputs 16,26,29,33,39,40,59,65,75,81,90,100,110,145,171,174,212,223,224,232,
234,235,239,252,253,255,263,265,269,276,277,279,280,283,284,286,291,296,302,308,
312,319,320,323,325,329,331,336,341,342,344,345,352,358,361,366,368,370,372,293
--splitting_before_polco false
```

Running with `mpiexec -n 4`, implies that some of the tasks are spread over 4 computation cores. For this model, it turned out to be beneficial to use the indirect method, `--direct false`. When we use indirect intersection, only the redundancy removal can be parallelised. The indices for the `inputs`- and `outputs`-arguments were determined by first running `ecmtool` with the default argument `--print_metabolites true`. In the printed list with indices and metabolites, we could find all metabolites that were mentioned by the model developers [?] in a minimal medium. We used `--splitting_before_polco false`, simply because this seemed to enable faster progress. For now, we cannot really determine when it is wise to use this option or not, so that trial and error is the best we can do. This computation took several hours to run.

If one is mostly interested in the conversion of inputs to biomass, it might be sufficient to calculate the ECMs in the network while all outputs are hidden. In Figure 5 of the main text, we show the results obtained by this command:

```
--model_path models/iIT341.xml --direct false
--inputs 139,262,28,294,300,306,314,231,35,350,259,261,22,356,334,93,293,271
--out_path iIT.csv
--outputs 16,26,29,33,39,40,59,65,75,81,90,100,110,145,171,174,212,223,224,232,
234,235,239,252,253,255,263,265,269,276,277,279,280,283,284,286,291,296,302,308,
312,319,320,323,325,329,331,336,341,342,344,345,352,358,361,366,368,370,372,293
--hide_all_in_or_outputs output
```

So, we added the argument –hide_all_in_or_outputs output, and removed `--splitting_before_polco false`. The first ensures that all output-metabolites are hidden. Hiding these output metabolites is only possible if metabolites that are both in- and output are already split before the enumeration step. Therefore, the argument `--splitting_before_polco false` is overridden by `ecmtool` anyhow.

##### *Escherichia coli*-model: iJR904

This large *E. coli*-model can also be downloaded from `bigg.ucsd.edu`. We used it to calculate all relations between glucose and oxygen consumption, and biomass production. These relations are shown in Figure 6 of the main text. The computation must be done using the direct method, and on many computation cores. We have used 20 nodes each of which had 16 computing cores, and the computation took 14.5 hours. The full jobscript that we used was:

```bash
#!/bin/bash
#SBATCH -N 20
#SBATCH -t 48:00:00

module load 2019
module load Python/3.6.6-intel-2019b
export PATH=$PATH:~/lrslib/lrslib-070

mpiexec python3 ~/ecmtool/main.py --model_path ~/ecmtool/models/iJR904.xml
--inputs 374,542,551,589,660
--hide 542,589,660,3,6,20,27,65,68,77,120,122,124,130,133,136,144,146,
164,167,169,173,180,189,192,195,197,206,211,219,223,229,233,236,242,244,247,
249,260,265,268,280,286,288,305,315,324,328,333,336,338,340,349,352,354,356,
358,360,363,366,376,378,380,382,388,393,395,397,399,402,404,413,419,424,427,
431,436,447,452,455,459,462,466,474,476,479,481,490,493,497,500,502,504,506,
509,510,513,515,517,525,532,534,536,542,545,547,549,555,562,583,589,592,603,
611,616,623,633,638,643,649,651,660,662,668,674,680,682,688,692,695,697,699,
704,707,709,711,714,742,745,747,750,752,755,759
--out_path ~/output/iJR.csv
--direct true
```

One might notice that the lists of metabolites to hide can become quite long, and therefore it might be tedious to manually compile these. We therefore first used `ecmtool` as a Python library, and created a small script that returned lists of the metabolite indices that should be hidden.

##### *Saccharomyces cerevisiae*-model: iND750

This model, also downloaded from `bigg.ucsd.edu`, was analysed in the paper that originally introduced ECMs [?]. We found that the import and export of many external metabolites was prohibited by the authors. This is also possible with the `ecmtool`-argument `--prohibit`. It makes sure that only the ECMs are returned that do not involve one of the prohibited metabolites. Although this functionality enables the calculation of some ECMs for very large models, it might not be very biologically reasonable.

Our runscript and the computed ECMs can be found in the supplementary files `iND750_runscript.txt` and `iND750_indirect.csv`.
